# Reticulate Evolution and Plastome Restructuring Shape Asian *Passiflora* Diversification

**DOI:** 10.64898/2026.09.23.753946

**Authors:** Zhou Zi Yu, Shook Ling Low, Niu Hong Bin, Song Shi Jie, Wu Fu Chuan, Shen Jian Yong, Dong Hui, Jiang Qiu Yu, Landrein Sven

**Affiliations:** Center for Gardening and Horticulture, Xishuangbanna Tropical Botanical Garden, Chinese Academy of Sciences, Xishuangbanna 666303, China; Forest Research Institute Malaysia 52109 Kepong, Selangor Darul Ehsan, Malaysia; Funing County Shizhuang Crop Pest and Disease Monitoring Station; Fairylake Botanical Garden 160 Xianhu Road, Luohu District, Shenzhen 518004 Guangdong

## Abstract

Asian Passifloraceae occupy a distinctive evolutionary and ecological position within the family despite their relatively low species diversity compared with the Neotropics. Many Asian representatives are highly localized, small and often ephemeral climbers that persist seasonally through rhizomes, tuberous stems or dormant seeds, while considerable morphological plasticity may obscure species limits and evolutionary relationships. Xishuangbanna, southwestern China, lies at the convergence of Southeast Asian, Indian-Himalayan and southern Chinese floristic elements and represents an important regional centre of Passifloraceae diversity, with two species of *Adenia* and five species of *Passiflora*, including the recently described *P. xishuangbannaensis* and *P. menghaiensis*.

We investigate the evolutionary relationships, plastome evolution and population genomics of these taxa using complete plastome assemblies, a concatenated dataset of 68 selected plastid protein-coding genes, 356 reconstructed nuclear CoDing Sequences (CDS), and Genotyping-By-Sequencing (GBS) across the known wild distribution of *P. xishuangbannaensis*. Plastid, concatenated nuclear and nuclear coalescent phylogenies were strongly incongruent and differed from published classifications based on morphology and earlier phylogenies (*ITS*, *ncpGS*, *ctyGS*, *ndhF* and *trnL*-*F*). Gene-tree discordance and phylogenetic-network inference indicated widespread incomplete lineage sorting together with introgression. The two statistically best-supported network models placed *P. menghaiensis*, which showed high genetic similarity to the Indian *P. napalensis*, as a sister lineage involved in independent reticulation events associated with *P. altebilobata*, *P. xishuangbannaensis* and *P. sumatrana*. A pronounced biogeographic pattern was evident within *Passiflora* supersection *Disemma*, particularly in the *P. siamica*-*P. cochinchinensis* lineage across the Hua Line, whereas relationships among other Asian *Passiflora* blurred this boundary. This pattern suggests that diversification and reticulation within the Asian lineage may have overlapped temporally with the development of the biogeographic divide represented by the Hua Line. The concentration of early-diverging lineages around Xishuangbanna and the eastern Himalayan-Indian region further points to this area as an important centre in the early diversification of Asian *Passiflora*. Within this context, the highly localized *P. xishuangbannaensis* may represent either a relictual of an early Asian lineage or the surviving product of an ancient reticulate complex, contrasting with lineages such as *P. siamica* that subsequently expanded farther across Southeast Asia.

Comparative plastome analyses revealed extensive structural and gene-content evolution within subgenus *Decaloba*, including pseudogenization, gene and intron losses, and repeated Inverted-Repeat (IR) expansions and contractions; substantial plastome modification also characterized supersection *Disemma*. *Passiflora sumatrana* exhibited a particularly distinctive IR expansion extending into the *accD*-*pafII* region, potentially characterizing this lineage. *Adenia* plastomes were generally more structurally conservative, although *A. penangiana* displayed an IR-boundary contraction involving the *rps19*-*rpl2* region. Mitochondrial CDS and non-coding regions provided the best-resolved phylogenetic signal and revealed strong discordance with the chloroplast phylogeny, consistent with maternal mitochondrial inheritance and variable plastid transmission, potentially involving paternal or biparental inheritance and heteroplasmy.

GBS analyses of *P. xishuangbannaensis* using complementary distant-reference, de novo and closely related-reference approaches showed that mapping to *P. organensis* provided the clearest SNP distribution and population resolution. ADMIXTURE, PCA and *F_ST_* analyses revealed two strongly geographically structured populations with additional substructure and substantial differentiation (*F_ST_* = 0.112). Linkage disequilibrium based demographic reconstruction using GONE2 indicated declining effective population size (*N*_e_) toward the present in both populations, with a particularly abrupt recent decline in Pop1 and a more gradual decline in Pop2. Few candidate Self-Incompatibility (SI) genes could be confidently reconstructed, including limited recovery of highly variable docking regions. However, the intracellular region of an S-locus Receptor Kinases (SRK) contained a distinctive non-synonymous substitution predicted to alter protein configuration without major disruption of overall folding, potentially contributing to compatibility differences between populations.

Together, these results reveal a complex evolutionary history combining historical dispersal, incomplete lineage sorting, reticulation, morphological plasticity, plastome structural evolution and strong population differentiation. The narrow distributions and recent demographic decline of these newly recognized Xishuangbanna taxa emphasize their conservation importance, while broader Asian taxon sampling and deeper genomic sequencing will be essential to resolve the evolutionary history of this discrete and poorly known Passifloraceae radiation.

## INTRODUCTION

*Passiflora* L. is one of the largest genera of Passifloraceae, comprising more than 500 species and reaching its greatest diversity in the Neotropics. Against this predominantly American radiation, the comparatively small Old-World assemblage is biogeographically and evolutionarily distinctive. Twenty-two species have traditionally been recognized from Indochina, Southeast Asia and the Australasian members of subgenus *Decaloba* (DC.) Rchb. supersection *Disemma* (Labill.) J.M.MacDougal & Feuillet, a lineage whose systematic position was debated for more than two centuries (Krosnick & Freudenstein, 2005; Krosnick, 2006; Krosnick et al., 2013). Current circumscription recognizes three sections within *Disemma*: the Asian and Southeast Asian section *Octandranthus* Harms, the Australian section *Disemma*, and the monotypic section *Hollrungiella* Harms. from Papua New Guinea (Krosnick, 2006; Krosnick et al., 2013).

Section *Octandranthus* is particularly important for understanding the Asian radiation of *Passiflora*. Krosnick (2006) treated 17 species distributed from India and China through Southeast Asia and identified China as its principal centre of diversity. The lineage is characterized most consistently by a reduced first-order inflorescence axis with higher-order cymose branching and two series of coronal filaments, whereas many vegetative and floral characters; including leaf shape, nectary position and type, and reproductive organ number are highly variable (Krosnick, 2006). The morphological characters plasticity is striking: *P. cochinchinensis* Spreng. exhibits the unusual subopposite to opposite leaf arrangement, while *P. siamica* Craib., *P. tonkinensis* W.J.de Wilde and *P. perakensis* Hallier F. may vary in stamen and carpel number, a condition exceptional within *Passiflora* (Krosnick, 2006; Krosnick et al., 2006). Such character variation, together with repeated convergence in floral form, helps explain the historical instability of classifications based primarily on morphology (**Figure 1**).

**Figure 1.**
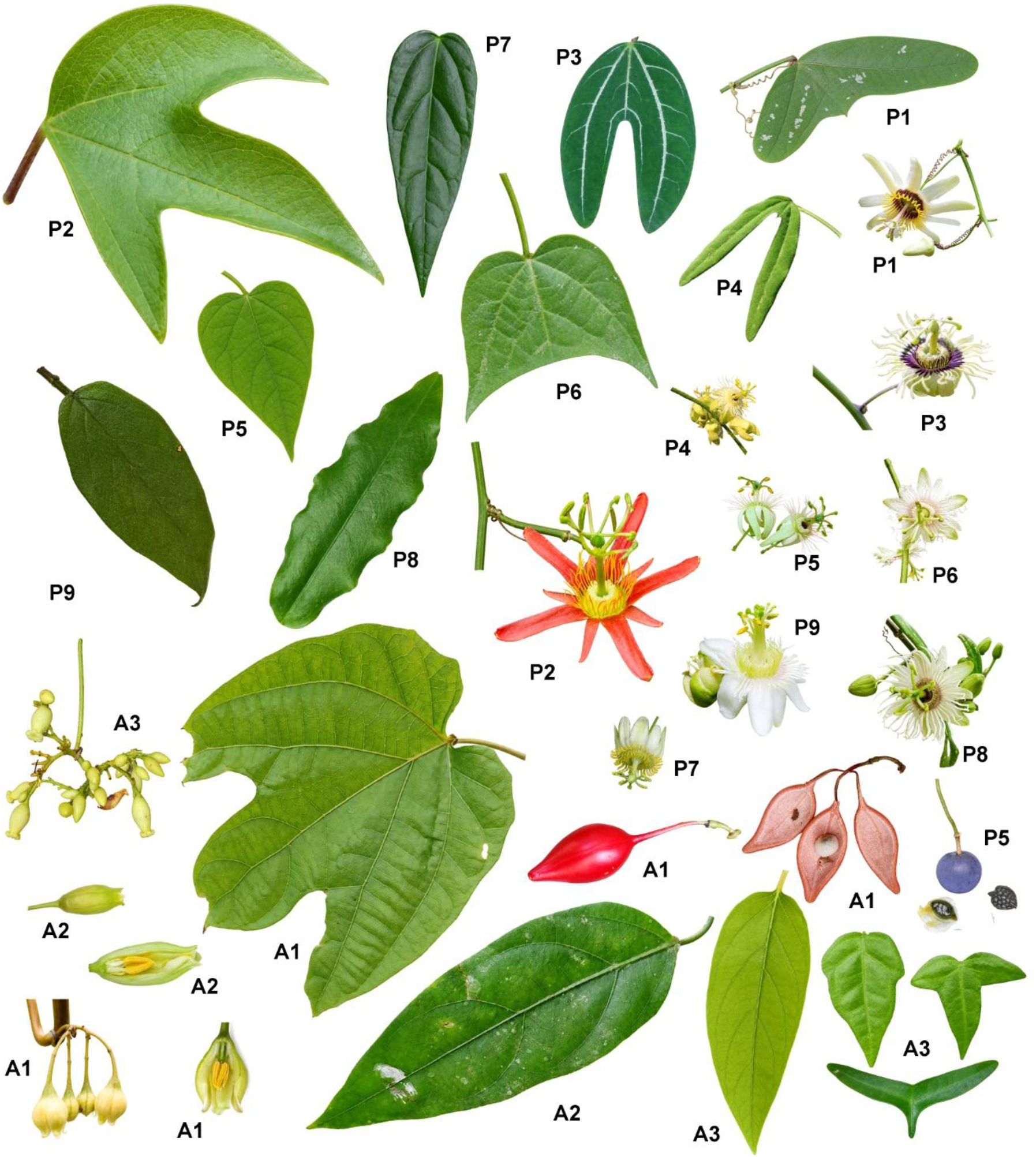
Morphological diversity in Asian *Adenia* and *Passiflora* supersection *Disemma*. **A1-A3**, *Adenia*: **A1**, *A. cardiophylla -* leaf, inflorescence, male flower with outer perianth removed, capsule, and seed; **A2**, *A. penangiana* - leaf, male flower, and male flower with outer perianth removed; **A3**, *A. cordifolia* - juvenile and adult leaves, inflorescence (Borneo, Sabah). **P1-P9**, *Passiflora*: **P1**, *P. biflora* - leaf and flower; **P2**, *P. cinnabarina* - leaf and flower; **P3**, *P. xishuangbannaensis* - leaf and flower; **P4**, *P. altebilobata* - leaf and flowers; **P5**, *P. menghaiensis* - leaf and flower; **P6**, *P. sumatrana* - leaf and flower; **P7**, *P. kwangtungensis* - leaf and flower (Guangdong, Yangchun); **P8**, *P. cochinchinensis* - leaf and flower (Kadoorie Farm and Botanic Garden, Hong Kong); **P9**, *P. siamica* - leaf and flower. All photographs by Sven Landrein, taken in Xishuangbanna unless otherwise indicated. Leaves and flowers are shown to scale relative to one another.

Molecular studies fundamentally changed the interpretation of these Old-World taxa. Analyses of nuclear and plastid loci demonstrated that it is monophyletic and nested within subgenus *Decaloba*, rather than representing an early-diverging lineage of *Passiflora* (Krosnick & Freudenstein, 2005; Krosnick, 2006). Broader sampling subsequently confirmed strong support for *Disemma* and section *Octandranthus*, while showing that the New-World *P. multiflora* L. is closely allied to the Old-World radiation (Krosnick et al., 2013). Using 148 taxa and four molecular markers (*nrITS*, *ncpGS*, *trnL–F* and *ndhF*) Krosnick et al. (2013) established one of the most comprehensively sampled phylogenetic frameworks for subgenus *Decaloba* section *Disemma*. Nevertheless, locus-specific conflicts remained, and unexpected plastid placements occurred probably due to paralogy, heteroplasmy or hybridization. Consequently, the evolutionary history of the Asian clade is particularly suited to an explicitly cytonuclear approach rather than inference from a single genomic compartment.

Southwestern China is central to this problem. Yunnan contains a large proportion of the Chinese diversity of section *Octandranthus*, and continuing fieldwork has revealed taxa with extremely restricted distributions (**Figure 2**). *Passiflora xishuangbannaensis* Krosnick was described as a Chinese endemic by Krosnick (2005), and *P. menghaiensis* X.D.Ma, L.C.Yan & J.Y.Shen was subsequently described from Menghai, Xishuangbanna (Ma et al., 2019). With the latter discovery, 14 species of section *Octandranthus* were recognized from China. Ma et al. (2019) further highlighted the concentration of native species in Yunnan and distinguished *P. menghaiensis* from morphologically similar *P. kwangtungensis* Merr. and *P. napalensis* Wall. by its two-flowered inflorescences, corona coloration, reduced branching and puberulous fruit. The diagnostic key presented by these authors further illustrates the mosaic distribution of characters among Chinese *Octandranthus*, including entire versus lobed leaves, petiolar and laminar nectaries, variable reproductive-organ number, pubescence and inflorescence architecture. Cytogenetic investigation has additionally shown *P. xishuangbannaensis*, *P. menghaiensis* and other native species from southwestern China to be diploid (2n = 12), while emphasizing the scarcity of detailed cytogenetic information for Asian *Passiflora* (Qian et al., 2023).

**Figure 2.**
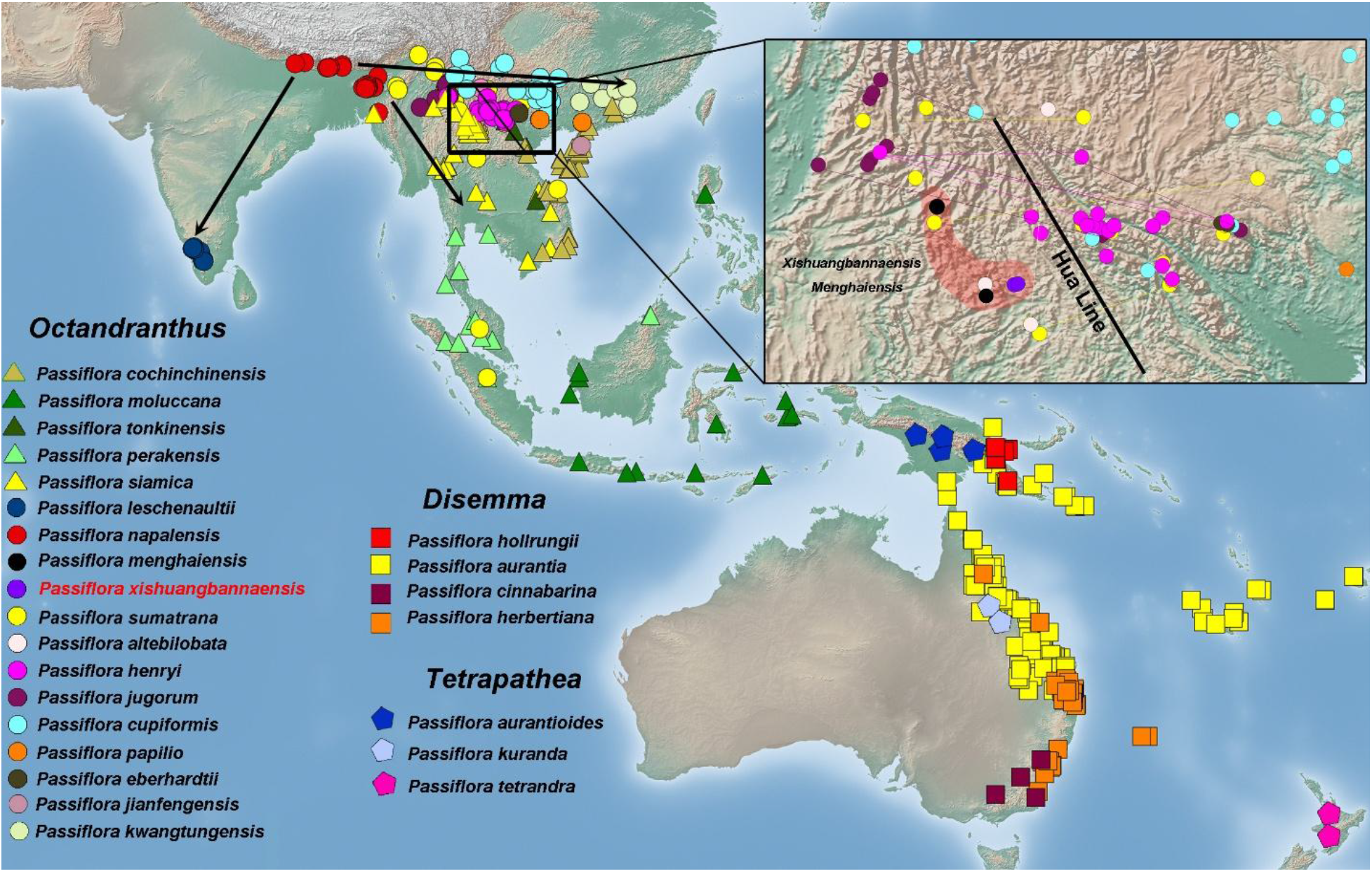
Geographic distribution of Asian *Passiflora*. Distribution of all known Asian *Passiflora* belonging to supersection *Disemma* and the historically recognized supersection *Hollrungia* as well as supersection *Tetrapathea* (DC.) P.S.Green. Supersection *Hollrungia* has subsequently been subsumed within *Disemma*, while supersection *Tetrapathea* has been redefined to include two additional species. Locality records were compiled primarily from Krosnick (2006) and georeferenced using a Phyton script from the locality information provided in specimen records; ambiguous or anomalous localities were manually verified and corrected where necessary. The Hua Line (Zhu, 2011), a proposed biogeographical boundary between southern and tropical southeastern Yunnan approximately following the Lixianjiang River is indicated. Although several taxa cross this boundary (connecting lines), the overall distribution shows marked geographical structuring, with differentiation among predominantly Indo-Malaysian, East/Southeast Asian, and Australasian elements. The distribution map and topographic layers were prepared and plotted using DIVA-GIS.

Despite these advances, relationships within the Asian radiation remain incompletely resolved. Earlier phylogenies were based on relatively few loci, and even the expanded four markers analysis of Krosnick et al. (2013) was primarily designed to resolve deeper infrageneric relationships within subgenus *Decaloba*, rather than recent divergences within *Octandranthus*. The same study documented locus-specific conflicts and warned that unexpected plastid placements may reflect missing data, heteroplasmy, paralogy or hybridization. Hao and Wu (2021) sequencing and phylogenetic analysis of the chloroplast genome of *P. xishuangbannaensis* extended study of the group to the genomic level and placed the species within subgenus *Decaloba*, but limited taxonomic sampling left finer-scale relationships among closely related Asian taxa insufficiently resolved. Thus, short evolutionary distances, Morphological plasticity, sparse genomic sampling and potential reticulation create precisely the conditions under which concatenated plastid or single-locus trees may provide an incomplete representation of species history.

Plastome evolution adds a second dimension to this problem. *Passiflora* is unusual among angiosperms for the structural dynamism of its chloroplast genomes. Comparative studies have documented extensive inversions and translocations, gene and intron losses, and major expansions and contractions of the inverted repeats, with particularly pronounced structural variation in subgenus *Decaloba* (Shrestha et al., 2019; Cauz-Santos et al., 2020, 2025). Cauz-Santos et al. (2020) sequenced 20 new plastomes, including 18 *Passiflora* species, and documented almost 50 kb of variation in plastome size, extensive inversions, IR expansions and contractions, lineage-specific gene losses, and complete loss of one IR in *P. capsularis* L. and *P. costaricensis* Killip. More recently, Cauz-Santos et al. (2025) analysed 61 taxa and confirmed pronounced variation in genome size, gene content, repeat composition, inversions and IR boundaries, particularly within *Decaloba*. Tests of selection additionally detected lineage-specific positive selection in genes associated with photosynthesis, gene expression and metabolism, including *clpP* and *petL* (Cauz-Santos et al., 2025). These structural features provide potentially informative evolutionary characters but also reinforce the need to compare plastid histories with independently inherited nuclear loci.

Interpretation of plastid phylogenies is further complicated by the unusual diversity of plastid inheritance in *Passiflora*. Experimental crosses have demonstrated maternal, paternal and biparental chloroplast transmission within the genus (Muschner et al., 2006; Hansen et al., 2007; Shrestha et al., 2021). Hansen et al. (2007) found predominantly paternal chloroplast inheritance in interspecific hybrids, contrasting with predominantly maternal inheritance in intraspecific crosses, in which biparental transmission was also detected. More extensive experiments involving 45 interspecific crosses subsequently demonstrated that inheritance patterns are partly lineage dependent: subgenera *Passiflora* and *Astrophea* (DC.) Mast predominantly showed paternal transmission, whereas subgenus *Decaloba* showed predominantly maternal or biparental inheritance (Shrestha et al., 2021). Importantly, heteroplasmy resulting from biparental inheritance was principally detected in cotyledons and first leaves and was subsequently lost through plastid sorting, so that mature plants generally retained only one detectable parental plastid type (Shrestha et al., 2021). Consequently, a plastome recovered from an adult individual may preserve only one component of a reticulate evolutionary history. Discordance between plastid and nuclear phylogenies in *Passiflora* therefore cannot automatically be attributed to phylogenetic error or incomplete lineage sorting; hybridization, differential plastid transmission and subsequent sorting provide biologically plausible mechanisms by which nuclear and plastid evolutionary histories may become decoupled.

The biogeographic history of *Passiflora* has received comparatively little attention relative to its extensive systematic study. The first broad molecular dating analysis of the genus inferred ancient divergence among its principal subgenera, approximately 32–38 Ma, followed by considerably younger diversification within individual lineages (Muschner et al., 2012). Old-World *Passiflora* do not represent an early-diverging lineage of the genus, but are nested within subgenus *Decaloba*, with supersection *Disemma* comprising distinct Asian and Austral-Pacific lineages distributed from the Indian subcontinent and southern China through Southeast Asia to Australasia (Krosnick & Freudenstein, 2005; Krosnick et al., 2006). This distribution is particularly interesting in southwestern China, where major geological and climatic changes associated with the Himalayan uplift, displacement of the Indochina and Lanping–Simao blocks, and development of the Asian monsoon contributed to the assembly of the modern southern Yunnan flora. A major floristic boundary, the Hua Line, separates the more Indo-Malaysian flora of southern Yunnan from that of southeastern China, although its influence and timing vary among tropical plant lineages (Zhu, 2011). Phylogeographic reconstruction of another rainforest liana, *Eleutharrhena macrocarpa* Diels (Forman), demonstrated substantial Miocene differentiation across this region and dispersal into southern Yunnan, illustrating how present distributions may preserve signatures of these historical geographical divisions (Song et al., 2022). The geographical distribution of Asian *Passiflora*, particularly supersection *Disemma*, therefore provides an opportunity to test whether diversification and dispersal across southern China and Indochina similarly correspond to this biogeographic boundary.

Reproductive biology may provide an additional dimension to the conservation of geographically restricted *Passiflora*. Self-incompatibility (SI) has been demonstrated in several species, and reproductive studies indicate that its genetic control may involve both sporophytic and gametophytic components (Suassuna et al., 2003; Madureira et al., 2014). However, the molecular determinants of self versus non-self-recognition in *Passiflora* remain unresolved. The *Passiflora organensis* Gardner reference genome provided the first genomic investigation of candidate SI-associated genes, identifying 54 proteins with similarity to S-locus Receptor Kinase (SRK) and S-Locus Glycoprotein (SLG) proteins, including a genomic cluster containing adjacent SRK and SLG-like candidates (Costa et al., 2021). These candidates were identified using Brassicaceae S-locus sequences and partial *P. edulis* Sims. sequences, and therefore their functional involvement in *Passiflora* SI remains to be demonstrated. Nevertheless, because reproductive compatibility can become particularly important in small and genetically depauperate populations, variation associated with candidate SI loci represents a potentially relevant component of the conservation genetics of narrowly distributed species such as *P. xishuangbannaensis*.

*Passiflora xishuangbannaensis* is a highly localized endemic restricted to Xishuangbanna and recognized in China as a Plant Species with Extremely Small Populations (PSESP), (Yang et al., 2020). Surveys of all known localities subsequently recorded only 38 wild individuals, with remaining populations threatened particularly by agricultural and road disturbance (Meng et al., 2021). Earlier XTBG surveys had found fewer than 100 plants distributed among only three populations, two of which occurred outside protected areas. Conservation efforts at Xishuangbanna Tropical Botanical Garden (XTBG) have included vegetative propagation, grafting, ex situ cultivation and experimental reintroduction: more than 100 plants were initially propagated, and 60 individuals were reintroduced into Mengyang Nature Reserve in 2019 for subsequent monitoring. However, difficulties in sexual reproduction and the extremely small number of surviving wild individuals emphasize the need to understand the species remaining genetic diversity, population structure and evolutionary history before further reinforcement or reintroduction are undertaken.

Here, we integrate complete plastome comparisons, plastid phylogenomics, reconstructed nuclear loci and population-level genomic data to investigate the evolutionary history of Asian *Passiflora*, with particular emphasis on section *Octandranthus* and the Xishuangbanna endemics. We ask whether plastome structural evolution provides phylogenetically informative patterns within the Asian lineage; whether nuclear gene trees and species-tree and network analyses are concordant with plastid relationships or instead reveal incomplete lineage sorting and reticulate evolution; and how genetic diversity and population differentiation are structured within *P. xishuangbannaensis*, including variation associated with candidate self-incompatibility loci. By integrating genomic compartments and analytical scales, this study aims to refine the evolutionary framework of a poorly sampled Old-World radiation and provide genomic information relevant to the conservation of these narrowly distributed Chinese species.

## MATERIALS AND METHODS

### Plant material, DNA extraction and sequencing

Leaf material was collected from both cultivated plants maintained at the Xishuangbanna Tropical Botanical Garden (XTBG), Chinese Academy of Sciences (CAS), and from natural populations of *Passiflora xishuangbannaensis* in Xishuangbanna, Yunnan, China, Hong Kong, China and Borneo, Sabah, Malaysia. Depending on the material and subsequent analyses, leaves were either immediately frozen in liquid nitrogen or preserved in silica gel. Voucher specimens were deposited in the Herbarium of Xishuangbanna Tropical Botanical Garden (HITBC), Xishuangbanna, Yunnan, China, and the Herbarium of Kadoorie Farm and Botanic Garden (KFBG), Hong Kong, China. Total genomic DNA was extracted using the Magnetic Plant Genomic DNA Kit (TIANGEN, China).

For species-level phylogenomic analyses, genomic libraries were prepared and sequenced by Annoroad Gene Technology (Beijing, China) using the Illumina NovaSeq 6000 platform, generating 150-bp paired-end reads. Adapter sequences and low-quality reads were removed from the raw sequencing data using Trimmomatic (Bolger et al., 2014) prior to downstream analyses **(Supplementary Table S1)**.

For population genomic analyses of *P. xishuangbannaensis*, Genotyping-By-Sequencing (GBS) library preparation and sequencing were performed by Novogene (Tianjin, China), **(Supplementary Table S2)**. Genomic DNA quality was assessed by agarose gel electrophoresis, purity was evaluated using a NanoDrop spectrophotometer, and DNA concentration was quantified using Qubit. GBS libraries were generated by restriction-enzyme digestion, with MseI used as the primary restriction enzyme and a second restriction enzyme (selected during library optimization) used to regulate the number of recovered tags. Digested fragments were ligated to P1 and P2 adapters containing 6 bp sample-specific barcode sequences, followed by PCR amplification, fragment-size selection, pooling and purification using AMPure XP beads. Library quality and insert size were assessed using Qubit 2.0 and an Agilent 2100 Bioanalyzer, and effective library concentration was determined by quantitative real-time PCR prior to sequencing. Libraries were subsequently pooled and sequenced on an Illumina NovaSeq 6000 platform to generate 150-bp paired-end reads.

### Chloroplast genome and nuclear ribosomal cistron assembly and annotation

Complete chloroplast genomes were assembled from whole-genome Illumina paired-end sequencing data for the newly sequenced *Passiflora* and *Adenia* Forssk. accessions using GetOrganelle v1.7.7.1 (Jin et al., 2020), with Bowtie2 (Langmead & Salzberg, 2012) for read recruitment and SPAdes (Bankevich et al., 2012) for de novo assembly. The GetOrganelle embryophyte plastome database (-F embplant_pt) was used for plastid-read recruitment. Assemblies were performed using eight threads and up to 40 extension rounds. For accessions that did not produce a resolved plastome under the initial settings, alternative k-mer combinations and closely related seed plastomes were tested.

Resulting assembly graphs were inspected in Bandage (Wick et al., 2015) to identify the principal plastome path and distinguish it from short alternative branches or low-coverage spurious sequences. Assemblies were further assessed by comparison with plastomes of closely related species and, where necessary, through inspection of read-mapping support. Because plastomes are circular molecules and equivalent assemblies may differ in their starting coordinate or orientation, completed genomes were normalized to a common starting position and orientation prior to comparative analyses. Attention was given to the orientation of the Large Single-Copy (LSC), Small Single-Copy (SSC), and Inverted-Repeat (IR) regions to avoid interpreting alternative representations of the circular molecule as genuine structural rearrangements.

Completed plastomes were annotated using GeSeq (Tillich et al., 2017), with closely related *Passiflora* and *Adenia* plastomes used as references. Transfer RNA annotations were additionally checked using tRNAscan-SE (Chan et al., 2021). Automated annotations were subsequently subjected to extensive manual inspection in GenomeView (Abeel et al., 2012), including examination of gene positions and boundaries, coding-sequence integrity, exon-intron structure, duplicated and truncated genes, and putative pseudogenes. Problematic regions were compared directly among accessions and with closely related reference plastomes to evaluate and, where appropriate, correct inconsistent annotations. Attention was given to genes showing apparent duplication, truncation, internal stop codons or unusual intron structure, as well as genes located within or adjacent to the IRs. Putative gene losses and pseudogenes were manually verified by examining sequence length, reading-frame integrity, start and stop codons, exon-intron structure, genomic position and neighbouring gene order. Duplications associated with IRa and IRb and partial gene copies generated at IR boundaries were distinguished from pseudogenes. The multipart organization of the trans-spliced *rps12*, alternative initiation of *ndhD*, and differences resulting solely from intron or feature annotation were also considered. Genes were classified as putatively pseudogenized only when manual inspection supported substantial truncation, fragmentation or loss of coding integrity, while apparent gene losses were accepted only after examination of the corresponding genomic region excluded annotation failure or the presence of an unannotated homologous sequence.

Whole-plastome structural variation was investigated using progressiveMauve (Darling et al., 2010) to identify conserved locally collinear blocks and large-scale rearrangements among representative Passifloraceae plastomes. The positions of the LSC, SSC, IRa and IRb regions were determined from the final annotations, and the identities and orientations of genes occurring at or spanning their boundaries were manually verified in GenomeView. These curated annotations were subsequently used to compare plastome size and organization, gene content, gene loss and pseudogenization, and IR-boundary expansions and contractions among taxa.

The nuclear ribosomal cistron, comprising the 18S rRNA gene, ITS1, 5.8S rRNA gene, ITS2 and 26S rRNA gene, was assembled independently from the same whole-genome Illumina reads using GetOrganelle v1.7.7.1 with the nuclear ribosomal DNA database (-F embplant_nr). The resulting nrDNA sequences were checked for completeness and consistency before subsequent comparative and phylogenetic analyses.

### Phylogenetic analysis of plastome protein-coding genes, nuclear ribosomal cistron, nrITS, *cytGS*, *ncpGS*, *trnL–trnF* and *ndhF*

Phylogenetic relationships among the sampled *Passiflora* taxa were reconstructed using a concatenated dataset of 68 plastid protein-coding genes shared among the analysed plastomes **(Supplementary Table S3)**, together with the nuclear ribosomal ITS. The 68 plastid protein coding genes corresponded to the conserved coding regions used in previous comparative plastome phylogenetic analyses (Cauz-Santos et al., 2025).

Maximum-Likelihood (ML) phylogenetic analyses were conducted in IQ-TREE (Nguyen et al., 2015). The concatenated plastid dataset was analysed under a partitioned framework, with individual protein-coding genes treated as separate initial partitions. Best-fitting nucleotide substitution models and an optimized partitioning scheme were selected within IQ-TREE using ModelFinder (Kalyaanamoorthy et al., 2017) and partition merging (MFP+MERGE) reduced these 68 starting gene partitions to 10 optimized partitions. Branch support was assessed using 1000 ultrafast bootstrap replicates (Hoang et al., 2018). Bayesian phylogenetic inference of the plastid CDS dataset was additionally performed using MrBayes (Ronquist et al., 2012). To avoid the very slow convergence observed when all 68 genes were independently parameterized, MrBayes was run using a single unpartitioned GTR+Γ model, with two independent runs and four chains. The initial 25% of sampled trees were discarded as burn-in, and a majority-rule consensus tree was calculated from the post-burn-in samples. Convergence between independent runs was assessed from the stabilization of likelihood values and model parameters and by verifying that the average standard deviation of split frequencies had reached an acceptably low value. Posterior Probabilities (PP) calculated from the post-burn-in trees were used as Bayesian measures of branch support.

Previous phylogenetic analyses of supersection *Disemma*, particularly the comprehensive sampling of Krosnick (2006), included nearly all recognized species, apart from the subsequently described *Passiflora menghaiensis* and *P. kwangtungensis*. To integrate the newly sampled taxa into this broader phylogenetic framework and permit direct comparison with previous studies, five markers used in earlier phylogenetic analyses were examined: nuclear ribosomal ITS (nrITS), the low-copy nuclear genes *cytGS* and *ncpGS*, and the plastid regions *trnL*-*trnF* and *ndhF*. Plastid markers were extracted from the annotated plastome assemblies, whereas *cytGS* and *ncpGS* were reconstructed from the original Illumina reads in *Passiflora menghaiensis*. Sequences generated in the present study were combined with homologous sequences retrieved from GenBank (Sayers et al., 2024), **(Supplementary Table S4)**. Each marker was aligned independently using MAFFT (Katoh & Standley, 2013) and the resulting alignments were manually inspected prior to phylogenetic analysis. Maximum-likelihood phylogenies were inferred separately for each marker using IQ-TREE, with the best-fitting nucleotide substitution model selected using ModelFinder and branch support assessed by bootstrap analyses. Individual marker trees were subsequently compared with each other and with the 68-gene plastid CDS phylogeny to assess phylogenetic congruence and identify topological conflicts among plastid, nuclear ribosomal and low-copy nuclear markers.

Phylogenetic hypotheses obtained from the complete plastid CDS matrix, the Bayesian analysis, and the individual nuclear ribosomal cistron, ITS, *cytGS*, *ncpGS*, *trnL*-*trnF* and *ndhF* datasets were compared to evaluate the robustness of the recovered relationships and to identify phylogenetic discordance among genomic compartments.

### Mitochondrial phylogenetic analysis

Mitochondrial reads from eight *Passiflora* accessions were mapped against the *P. edulis* mitochondrial reference genome (MT140634; 680480 bp) using BWA-MEM (Li & Durbin, 2009). Consensus sequences were generated after coverage filtering (depth <5x were masked). The annotated reference contained 41 CDS features. Coding regions were extracted and concatenated, yielding a common callable CDS matrix of approximately 29.3 kb. Because mitochondrial coding sequences showed relatively limited variation, non-coding regions were additionally examined. Annotated gene regions were excluded from the reference and intergenic positions callable as unambiguous A/C/G/T in all eight accessions were retained. This produced 855 callable intergenic blocks ≥100 bp, with a maximum block length of 1785 bp and a total aligned length of 269898 bp. The intergenic matrix contained 3997 variable sites, of which 1027 were parsimony-informative (1.48% and 0.38% of sites, respectively). Pairwise divergence was consistently greater in intergenic than coding regions, generally by approximately 1.3-2.0 fold.

Coding and intergenic matrices were concatenated into a 299391 bp mitochondrial matrix and analysed using maximum likelihood in IQ-TREE, with coding and intergenic sequences treated as separate partitions and partition-specific substitution models selected by ModelFinder. Branch support was estimated using ultrafast bootstrap replicates. Bayesian inference was subsequently performed in MrBayes using separate models for the two partitions. Two independent MCMC runs were performed and stopped after 2 million generations following convergence. Convergence was assessed using the average standard deviation of split frequencies and Effective Sample Sizes (ESS).

### Nuclear gene recovery and phylogenomic analyses

Nuclear coding loci were recovered using the annotated *Passiflora organensis* genome (Costa et al., 2021) as the initial reference. Illumina paired-end reads of *Passiflora perakensis* were mapped with BWA-MEM and alignments processed with SAMtools (Li et al., 2009). Variants were called using bcftools (Danecek et al., 2021) with minimum mapping and base-quality thresholds of 20. Positions with sequencing depth <5× and regions overlapping indels were masked before construction of sample specific consensus sequences. Annotated CDS sequences were extracted from the *P. organensis* annotation using gffread (Pertea & Pertea, 2020). Where multiple transcript isoforms were annotated for a gene, the longest CDS isoform was retained.

Illumina whole-genome resequencing data from eight *Passiflora* accessions (Bor10, kfbg21, SRR13805787, sven30, sven31, sven32, sven33 and sven50) were used to recover orthologous nuclear coding regions. Reads were mapped with BWA-MEM against *Passiflora perakensis* derived nuclear reference generated using the annotated *P. organensis* genome (PRJNA690087), selected because of its higher depth sequencing. Alignments were coordinate-sorted with SAMtools, and variants were called with bcftools mpileup/call using minimum mapping and base-quality thresholds of 20. Positions with sequencing depth <5× and regions containing indels were masked, whereas heterozygous SNPs were retained as IUPAC ambiguity codes in sample-specific consensus sequences. Annotated CDS coordinates were transferred to the consensus sequences and coding regions were extracted using gffread, retaining the longest transcript when multiple transcripts were present. Genes were evaluated according to sequence recovery, with loci containing ≥90% resolved sequence and represented in at least six of the eight taxa initially retained; loci with insufficient recovery or excessive missing data were subsequently excluded **(List of selected genes in Supplementary Table S5)**.

The resulting nuclear loci were aligned individually and used for both concatenated and gene-tree-based phylogenetic analyses. For the concatenated analysis, the trimmed alignments were combined into a nucleotide supermatrix, with missing loci represented by Ns and each locus initially treated as an independent partition. Maximum-likelihood phylogenetic analysis was performed in IQ-TREE using ModelFinder with partition merging (MFP+MERGE) to optimize the partitioning scheme and select the best-fitting substitution models, resulting in nine optimized partitions. Branch support was assessed using 1000 ultrafast bootstrap (UFBoot) replicates and 1000 SH-like approximate likelihood-ratio test (SH-aLRT) replicates.

### Gene-tree, coalescent and discordance analyses

Maximum-likelihood gene trees were reconstructed independently for each of the 356 nuclear loci using IQ-TREE. ModelFinder was used to select the best-fitting nucleotide substitution model for each locus, and branch support was estimated using 1000 ultrafast bootstrap replicates. To reduce the influence of poorly resolved relationships on species tree reconstruction, branches with UFBoot support <50% were collapsed before coalescent analysis. The resulting gene trees were analysed under the multispecies coalescent using ASTRAL-IV (Zhang et al., 2018), employing the more thorough search option (-R). Branch support and gene tree concordance were evaluated using local posterior probabilities and quartet support statistics. The resulting ASTRAL species tree was compared with the concatenated nuclear topology to identify relationships characterized by substantial gene tree conflict.

Gene-tree discordance was additionally examined using PhyParts (Smith et al., 2015) to quantify concordant and conflicting gene tree topologies around branches of the ASTRAL species tree. Selected discordant relationships (α–γ) were subsequently examined using PhyTop (Than et al., 2008) to compare the frequencies of alternative quartet topologies. Under an ILS-only expectation, the two alternative discordant topologies are expected to occur at approximately equal frequencies; significant asymmetry between them was therefore interpreted as evidence inconsistent with ILS alone and used to identify relationships for subsequent phylogenetic-network analyses.

### Phylogenetic network analyses

Potential reticulate evolution was investigated using the 356 nuclear gene trees with the maximum pseudo-likelihood method InferNetwork_MPL implemented in PhyloNet v3.8.2 (Than et al., 2008). Models allowing a maximum of one, two or three reticulation events (H = 1, H = 2 and H = 3) were analysed, together with a zero-reticulation model (H = 0) representing the bifurcating-tree baseline. For each reticulation level, 20 independent searches were conducted to reduce sensitivity to local optima, and the highest-scoring network was retained. For each inferred reticulation, inheritance probabilities (γ) were extracted directly from the PhyloNet extended Newick output. Networks were converted to Dendroscope compatible extended Newick format and visualized using Dendroscope 3 (Huson & Scornavacca, 2012). Alternative network models were compared using their maximum pseudo-log-likelihood scores, with higher values indicating a more favourable balance between model fit and complexity. Alternative network models were also compared using the Akaike Information Criterion (AIC) and Bayesian Information Criterion (BIC) to evaluate improvement in model fit while accounting for increasing network complexity. AIC and BIC were calculated for the H = 1, H = 2 and H = 3 models, with lower values indicating a more favourable balance between model fit and complexity. Because InferNetwork_MPL is based on maximum pseudo-likelihood rather than a conventional full likelihood, AIC and BIC comparisons were interpreted as relative criteria for model selection rather than formal likelihood-ratio tests. Network topology, inheritance probabilities and model-selection results were subsequently evaluated together with the ASTRAL species tree, concatenated nuclear phylogeny, gene-tree discordance patterns and morphological evidence.

### GBS data processing, population structure and genetic diversity

Genotyping-By-Sequencing (GBS) data from *Passiflora xishuangbannaensis* were analysed to investigate population structure, genetic diversity, demographic history and variation associated with candidate self-incompatibility genes. Because no chromosome-level reference genome was available for the species, three analytical strategies were evaluated: mapping against *P. edulis*, reference-free locus assembly, and mapping against the more closely related *P. organensis* genome. Quality-filtered paired-end reads were first aligned to the *P. edulis* reference genome (GCA_002156105 https://ngdc.cncb.ac.cn/gwh/Genome/557/show) using BWA-MEM v0.7.19, alignments were processed with SAMtools, and variants were called with BCFtools. Because this strategy provided insufficient representation of shared loci among individuals after population-level filtering, it was not retained for downstream analyses. To provide a reference-independent assessment, the reads were subsequently analysed de novo with Stacks (Catchen et al., 2013), using ustacks for individual locus assembly, cstacks to construct the catalogue of homologous loci and sstacks to match individual loci to the catalogue; population-level filtering was then applied to the resulting dataset, which was used as an independent assessment of population structure and for examination of highly similar genotypes. The final reference-based analysis used the approximately 256 Mb *P. organensis* genome. Reads were aligned with BWA-MEM and BAM files processed with SAMtools, after which reference-based loci and genotypes were reconstructed with Stacks v2.68 gstacks using the Marukilow genotype model (var_alpha = 0.01, gt_alpha = 0.05). The final dataset comprised 39 individuals. SNPs were filtered with the Stacks populations module by requiring loci to occur in at least 70% of individuals within a population (-r 0.7), excluding variants with minor allele frequency <0.05 (--min-maf 0.05) and loci with observed heterozygosity >0.60 (--max-obs-het 0.6). Pop1 comprised samples 1–27, and Pop2 samples 29–43. VCF (Danecek et al., 2011), PLINK (Purcell et al., 2007) and PHYLIP datasets were generated, and linkage disequilibrium pruning for analyses requiring approximately independent markers was performed in PLINK v1.9 using -- indep-pairwise 50 5 0.2. As an independent variant-calling procedure, variants were also called directly from the reference-aligned BAM files with BCFtools and filtered using QUAL ≥ 20 and DP ≥ 5; this dataset was maintained separately from the reference-based Stacks population-genetic dataset. The latter was used for subsequent analyses of population structure, genetic differentiation and diversity, demographic history, genotype concordance and candidate self-incompatibility regions.

Population structure was investigated using principal component analysis (PCA) and model-based ancestry estimation with ADMIXTURE (Alexander et al., 2009). Multiple values of K were evaluated (Pritchard et al., 2000), and the most strongly supported clustering solutions were assessed using cross-validation together with the ΔK approach of Evanno et al. (2005). Individual ancestry coefficients were visualized as admixture proportions and compared with the PCA clustering pattern. Basic population-genetic diversity statistics were calculated from the filtered genotype dataset using PLINK, including the Number of alleles (Na), effective Number of alleles (Ne), Observed Heterozygosity (Ho), Expected Heterozygosity (He), and the Percentage of Polymorphic Loci (PPL). Genetic differentiation among populations was evaluated using pairwise (*F_ST_*), with particular emphasis on the comparison between Pop1 and Pop2; sample 28 was retained separately as Pop2 because of its intermediate genetic position. Linkage disequilibrium pruning was performed in PLINK using --indep-pairwise 50 5 0.2 before analyses requiring approximately independent markers. Analysis of molecular variance (AMOVA), (Excoffier et al., 1992) was performed on the LD-pruned SNP dataset (36470 SNPs) to partition genetic variation among and within the two populations. Significance of population differentiation (ΦST) was assessed using 999 permutations.

### Spatial visualization of population structure

To examine the spatial distribution of genetic structure, individual ancestry coefficients obtained from the population structure analyses were associated with the geographic coordinates of each sampled individual. Ancestry proportions were extracted for both the K = 2 and K = 3 models and combined with sample identifiers, latitude and longitude. Geographic and ancestry data were compiled into comma-separated files containing the sample identifier, geographic coordinates, sampling group and corresponding ancestry coefficients (Q values). All 39 individuals included in the final genomic dataset were successfully matched to geographic coordinates and ancestry estimates.

The resulting files were imported into QGIS (QGIS Development Team, 2026), with sampling localities plotted using their longitude and latitude coordinates. Individual ancestry proportions were represented geographically as pie charts, with each sector proportional to the corresponding Q ancestry coefficient. Separate maps were produced for K = 2 and K = 3. Esri satellite imagery was used as the background layer to provide fine-scale geographic context for the sampled populations. Maps were displayed at approximately 1:15000, allowing the spatial relationships among sampled individuals and surrounding landscape features to be visualized while retaining satellite imagery. Final maps were exported at high resolution for figure preparation.

### Clone detection and pairwise genetic relatedness

Potential clonality and close genetic relatedness among *Passiflora xishuangbannaensis* individuals were investigated using the final *P. organensis* reference GBS dataset. The PLINK dataset comprised 39 individuals and 4951 SNPs. Initial pairwise Identity-By-Descent (IBD) estimates were calculated in PLINK v1.9 using the genome procedure, providing Z0, Z1, Z2 and PI_HAT estimates for all pairs of individuals.

Because GBS datasets can contain substantial locus-specific missing data, SNP missingness was subsequently quantified. Mean locus-level missingness across the 4951 SNP dataset was 45.3%, and 4745 of 4951 SNPs (95.8%) had greater than 5% missing data. Consequently, PI_HAT estimates were not used alone to identify clonally identical individuals, because differences in locus representation among samples and population structure could influence conventional IBD estimates.

Clone detection was therefore evaluated more directly using pairwise genotype concordance. Genotypes were exported from PLINK and all possible pairwise comparisons among the 39 individuals were performed, yielding 741 individual pairs. For each pair, only SNP loci for which a genotype call was available in both individuals were included. Loci missing in either individual were excluded from that particular comparison. Genotype concordance was calculated as the proportion of jointly genotyped loci at which the two individuals possessed identical diploid genotype calls. The number of jointly genotyped SNPs, identical genotypes, discordant genotypes and percentage concordance were recorded for every pair. A complete 39 × 39 pairwise concordance matrix and heatmap was generated.

### Demographic history

Historical and recent changes in effective population size (*N*_e_) were reconstructed independently for the two principal genetic populations of *Passiflora xishuangbannaensis* using complementary site-frequency-spectrum and linkage disequilibrium-based approaches. Long-term demographic history was inferred with Stairway Plot 2 (Liu & Fu, 2020) from folded Site-Frequency Spectra (SFS) generated separately for Pop1 and Pop2, with folding used because ancestral and derived allele states could not be reliably determined. A mutation rate of 1.0 x 10^-8^ substitutions site^-1^ generation^-1^ and a generation time of 24 years were specified to convert inferred demographic times to calendar years, and the resampling procedure implemented in Stairway Plot 2 was used to characterize uncertainty around the reconstructed trajectories.

Recent demographic history was independently investigated using linkage disequilibrium with GONE2 v2.0 (Santiago et al., 2025). Reads were mapped to the *P. organensis* reference genome and variants jointly called with BCFtools using minimum mapping and base qualities of 20. Only biallelic SNPs were retained, genotypes with depth (<3) were treated as missing, and each population was filtered independently to retain loci genotyped in at least 90% of individuals and with minor allele frequency ≥0.05, thereby minimizing LD generated by population structure. Because the *P. organensis* reference is scaffold-based, GONE2 analyses were restricted to major scaffolds with experimentally supported chromosome-arm assignments. Scaffold 1 and scaffold 2 represented chromosomes 1 and 4, respectively, while scaffolds 7+3 and scaffolds 4+8 were combined according to their published assignments to reconstruct chromosomes 5 and 6. Physical distances within scaffolds were preserved and gaps between chromosome arms were approximated from cytologically estimated chromosome lengths; the resulting coordinates therefore represent approximate linkage coordinates rather than complete chromosome assemblies. Genotypes and reconstructed coordinates were converted to PLINK PED/MAP format. GONE2 analyses were conducted separately for Pop1 and Pop2 under the standard unphased diploid model (g0). No LD pruning was applied because demographic inference in GONE2 relies directly on linkage disequilibrium among markers, and, in the absence of a species-specific recombination map, a constant recombination rate of 1.0 cM Mb^-1^ was assumed. Robustness to stochastic optimization was evaluated using five independent runs with random seeds 111, 222, 333, 444 and 555 while keeping all other parameters constant, and consistency among replicate trajectories was used to assess the stability of recent *N*_e_ inference.

### Self-incompatibility receptor analysis

Variation in the putative stigma-side self-incompatibility receptor was investigated using the S-locus candidate annotations reported for the *Passiflora organensis* genome. These candidates had been identified from homology to S-locus glycoproteins and S-locus receptor kinases and their characteristic domain architecture. We focused on scaffold7_size10217246.2019, a receptor-like candidate associated with the characteristic B-lectin, S-locus glycoprotein and PAN extracellular domains, a transmembrane region and an intracellular kinase domain. Population variation was examined in the two principal *P. xishuangbannaensis* populations (Pop1, n = 25; Pop2, n = 13). Genotypes with read depth <3 were masked, and variants were retained only when callable in at least 60% of individuals in each population (≥15 individuals in Pop1 and ≥8 in Pop2). Differentiation between populations was estimated using the Weir and Cockerham (*F_ST_*) estimator implemented in VCFtools, and allele frequencies were compared between populations. The coding sequence of the .2019 receptor candidate was reconstructed and translated to verify reading-frame integrity and was aligned to the reference genomic region to determine the coding consequence of differentiated variants. The focal SNP was localized to CDS position 2206, corresponding to genomic position 9912928 on *P. organensis* scaffold 7, and codon reconstruction was used to determine the resulting amino-acid substitution. Because this substitution occurred within the predicted intracellular kinase region, its potential structural effect was explored using AlphaFold (Jumper et al., 2021) and implemented through ColabFold (Mirdita et al., 2022) predictions of residues 560-810. Models were generated for the alternative protein variants and compared in terms of predicted fold and model confidence.

## RESULTS

### Plastome structural variation, gene content and inverted-repeat boundaries

Plastome structure varied markedly among the sampled *Adenia* and *Passiflora*, particularly through changes in the extent of the Inverted Repeats (IRs) and corresponding variation in the Large Single-Copy (LSC) region. Among the *Adenia* plastomes for which the quadripartite structure was resolved, genome sizes were relatively similar, ranging from 164680 bp in *A. penangiana* (Wall. ex G.Don) W.J.de Wilde to 165364 bp in *A. mannii* Engl.. The SSC regions were comparatively stable (13780-13855 bp), whereas greater variation occurred in the LSC and IR regions. *Adenia mannii* possessed an LSC of 87176 bp and IRs of 32204 bp, *A. cardiophylla* Engl. an LSC of 93076 bp and IRs of 30283 bp, and *A. penangiana* an LSC of 90213 bp and IRs of 30326 bp. Gene content was also highly conserved, with 109-110 unique genes, including 75-76 protein-coding genes, 30 tRNAs and four rRNAs, with *rpl20* absent in *A. penangiana*. The IR showed a minor contraction in *A. penangiana* and was localized to the rps19-rpl2 interval (event 6), shifting the IR/LSC boundary relative to the other sampled *Adenia* and representing a lineage-specific structural change (**Table 1**).

**Table 1.**
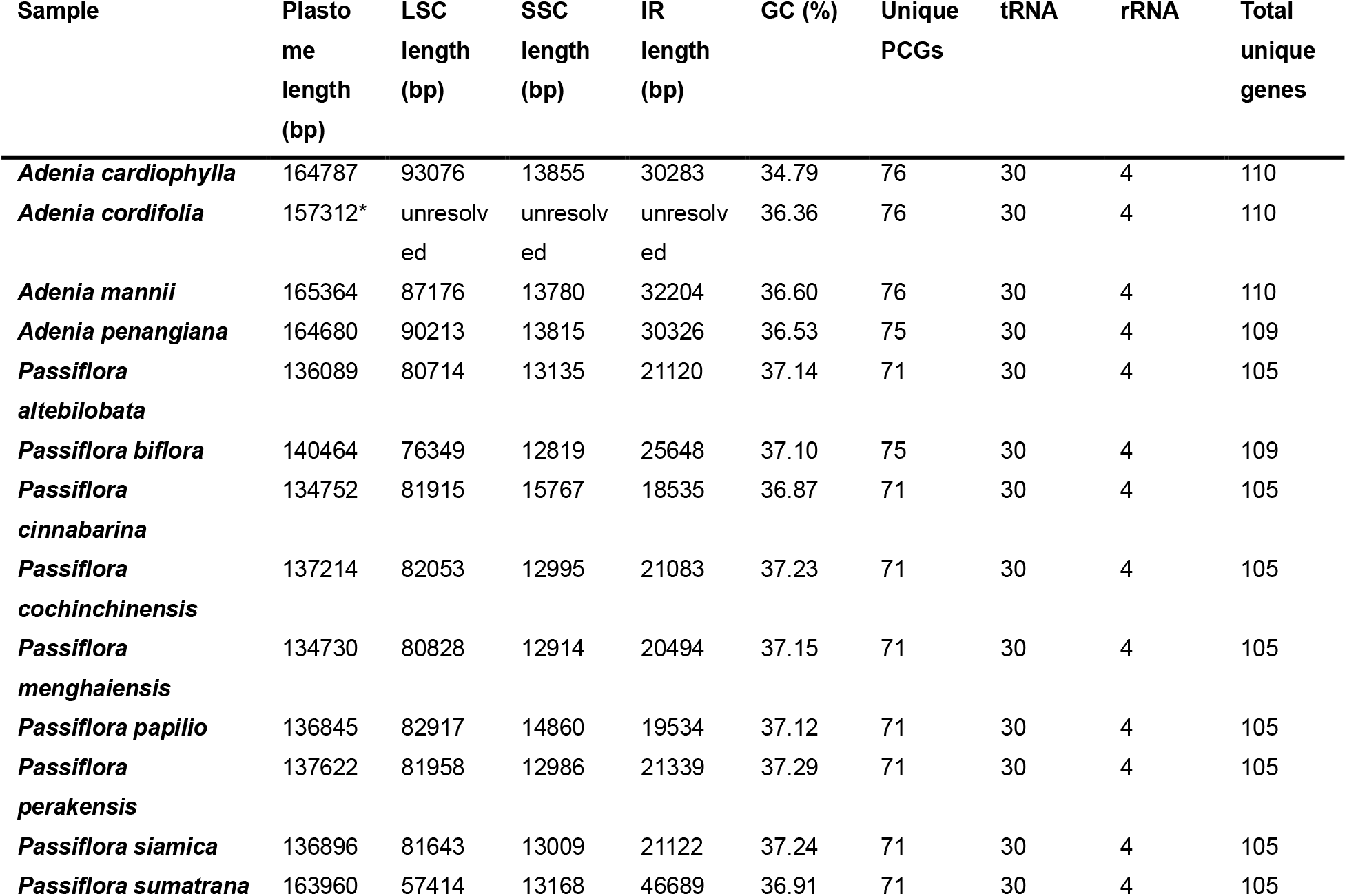
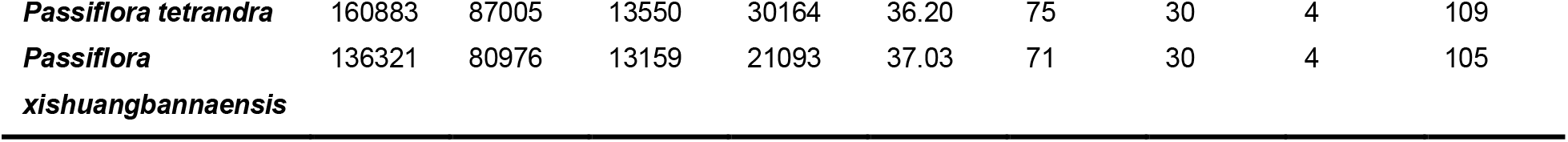
Plastome characteristics of the sampled *Adenia* and *Passiflora* species. Plastome length, large single-copy (LSC) and small single-copy (SSC) region lengths, inverted-repeat (IR) length, GC content, and numbers of unique protein-coding genes (PCGs), tRNA genes, rRNA genes and total unique genes are shown. Unresolved values indicate regions for which boundaries could not be reliably determined. * incomplete assembly (2 gaps)

*Passiflora tetrandra* exhibited a comparatively large plastome of 160883 bp, comprising an 87005 bp LSC, a 13550 bp SSC and two 30164 bp IRs. It contained 109 unique genes, including 75 protein coding genes, 30 tRNAs and four rRNAs. Most species of supersect. *Disemma* possessed considerably smaller plastomes. *P. altebilobata* Hemsl., *P. cochinchinensis*, *P. menghaiensis*, *P. perakensis*, *P. siamica* and *P. xishuangbannaensis* plastome size ranged from 134730 to 137622 bp. Their LSC regions were relatively uniform, ranging from 80714 to 82053 bp, as were their SSC regions (12914-13159 bp), while IR length ranged from 20494 bp in *P. menghaiensis* to 21339 bp in *P. perakensis*. *Passiflora xishuangbannaensis*, for example, possessed a 136321 bp plastome comprising an 80976 bp LSC, a 13159 bp SSC and two 21093 bp IRs (**Table 1**).

Two members of supersect. *Disemma* represented the extremes of structural variation. *Passiflora cinnabarina* Lindl. had the smallest plastome examined (134752 bp) and strongly contracted IRs of 18535 bp, together with an 81915 bp LSC and a comparatively enlarged SSC of 15767 bp. In contrast, *P. sumatrana* Blume possessed a 163960 bp plastome characterized by an exceptionally large IR expansion and a correspondingly reduced LSC. Its LSC was only 57414 bp, whereas IRa and IRb reached 46661 and 46717 bp, respectively, with an SSC of 13168 bp. The junction and phylogenetic reconstruction localized this expansion to the *accD-psaI* interval (event M), resulting in the incorporation into the IRs of a substantial region located within the LSC in the other sampled members of supersect. *Disemma*. This lineage-specific expansion therefore accounted for the combination of exceptionally large IRs and a strongly reduced LSC in *P. sumatrana* (**Table 1**).

Gene content also differed between the major lineages examined. Supersect. *Disemma* shared a reduced complement of plastid genes relative to *Adenia*, *P. tetrandra* Banks ex DC. and *P. biflora* Lam.. Seven genes or gene families were lost or pseudogenized in the lineage: *accD*, *rpl20*, *rpl22*, *rpl32*, *rps7*, *rps16* and *ycf1*/*ycf2*. No additional gene losses were detected among the sampled species within supersect.

*Disemma*, indicating that this reduced gene complement was shared across the section. In contrast, loss or pseudogenization of rpoA, observed in *P. tenuiloba* Engelm. and *P. suberosa* L., was not shared by supersect. *Disemma*. Accordingly, all sampled *Disemma* species contained 105 unique genes, comprising 71 protein-coding genes, 30 tRNAs and four rRNAs. GC content was conserved, ranging from 36.87% to 37.29%. Thus, the substantial variation in plastome size within supersect. *Disemma* occurred without further changes in unique gene number and was associated predominantly with structural changes affecting the IR and LSC regions (**Figure 3B**).

**Figure 3.**
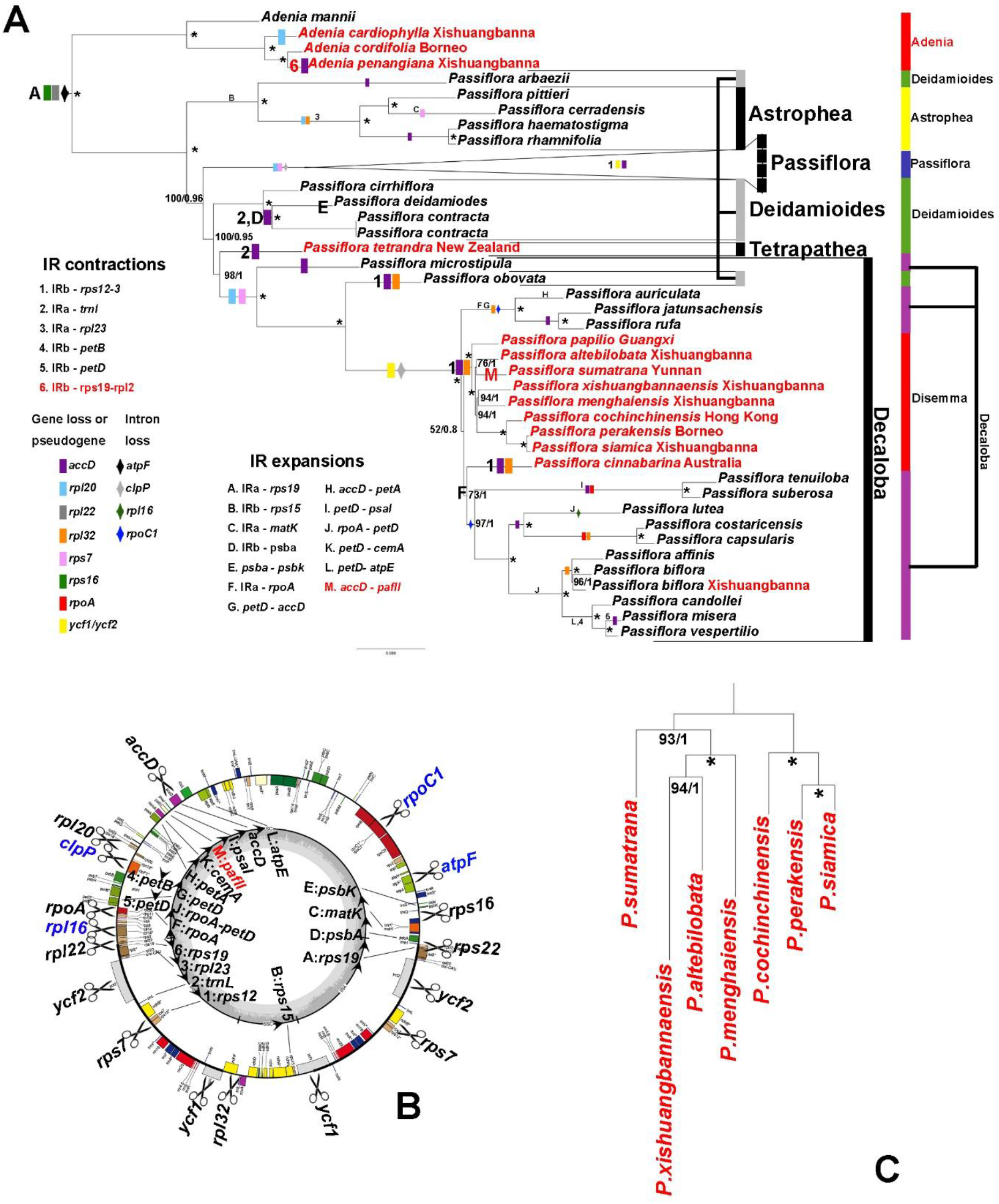
Plastome structural evolution and organellar phylogenetic relationships in Passifloraceae. (A) Maximum-likelihood/Bayesian plastid phylogeny showing relationships among *Adenia* and *Passiflora* accessions, with major plastome structural changes mapped onto the corresponding branches. Colored symbols indicate inferred gene losses or pseudogenization and intron losses, whereas letters and numbers denote expansions and contractions of the inverted repeat (IR) regions, respectively, as defined in the accompanying legends. Major *Passiflora* lineages are indicated on the right. Newly sequenced accessions and Asian taxa are highlighted in red, with their geographic origins indicated. Node values represent branch support, and asterisks denote maximally supported nodes. Terminal branches of the large subgenus *Passiflora* were omitted for clarity. (B) Representative plastome map illustrating the approximate positions of the gene losses or pseudogenization, intron losses, and IR-boundary expansions and contractions identified in the comparative analysis. Blue indicates intron loss rather than genes loss or pseudogene (C) Mitochondrial phylogeny of the focal Asian *Passiflora* taxa inferred from a concatenated matrix comprising mitochondrial protein-coding genes and non-coding regions; node labels indicate branch support and asterisks denote maximally supported nodes. Together, the analyses highlight extensive lineage-specific organellar genome evolution, including repeated IR-boundary shifts and recurrent plastid gene loss or pseudogenization, particularly within subgenus *Decaloba*, while allowing comparison of plastid and mitochondrial evolutionary histories.

ProgressiveMauve comparisons further revealed pronounced variation in plastome organization across the sampled Passifloraceae. Although large portions of the plastomes remained homologous among taxa, the number, orientation and arrangement of locally collinear blocks varied substantially. Several plastomes retained broadly comparable arrangements of the major homologous blocks, whereas others contained large blocks in reverse orientation. These inversions involved substantially larger portions of the plastome than the localized structural changes associated with shifts in individual IR boundaries. Overall, the structural comparisons revealed a combination of large-scale inversions and rearrangements, extensive variation in IR boundaries, and localized gene loss or pseudogenization across the sampled lineages **(Supplementary Figure S6 & Figure S7)**

### Phylogenetic relationships inferred from plastid protein-coding genes

The final concatenated plastid CDS dataset comprised 69 taxa, 68 protein-coding genes and 55217 aligned nucleotide positions. Missing data were extremely low (0.086%). Of the aligned sites, 43762 were constant, 11378 were variable and 7474 were parsimony-informative, representing 13.54% of the alignment.

Within *Passiflora* supersection *Disemma* section *Octandranthus*, the plastid phylogeny recovered several distinct lineages. *Passiflora altebilobata* and *P. sumatrana* formed a moderately supported clade (76% BP/1.0 PP), representing one of the weaker relationships within the section. *Passiflora xishuangbannaensis* and *P. menghaiensis* recovered as closely related, with strong support for the node uniting them (94% BP/1.0 PP). This lineage was in turn sister to the well-supported clade comprising *P. cochinchinensis*, *P. siamica* and *P. perakensis* (94% BP/1.0 PP). *Passiflora papilio* was recovered as sister to the remaining members of the section with maximal support (100% BP/1.0 PP). The Australian *P. cinnabarina* occupied a comparatively poorly resolved position relative to supersect. *Disemma* lineages. The deeper node associated with its placement received only moderate bootstrap support (73% BP), despite a Bayesian posterior probability of 1.0. Given this discrepancy between support measures and the very limited representation of Australian section *Disemma* in the present sampling, its precise relationship to the Asian lineages should be regarded as unresolved rather than as evidence for a definitive phylogenetic affinity.

Relationships among the sampled *Adenia* were more clearly resolved. *Adenia penangiana* from Xishuangbanna and the Bornean *A. cordifolia* were recovered as the closest relatives, with maximal support (100% BP/1.0 PP), while *A. cardiophylla* was successively related to this pair and *A. mannii* was more distant within the sampled *Adenia*.

Overall, the plastid phylogeny strongly resolved several terminal relationships within supersect. *Disemma*, particularly the *P. cochinchinensis*-*P. siamica*-*P. perakensis* group and the close relationships among *P. papilio*, *P. xishuangbannaensis* and *P. menghaiensis*, whereas support decreased substantially toward some of the deeper relationships connecting these lineages. The topology provided the phylogenetic framework for mapping plastome gene loss and pseudogenization, structural rearrangements, and IR expansions and contractions across the sampled lineages (**Figure 3A**).

The combined mitochondrial matrix produced a substantially better resolved phylogeny than the mitochondrial CDS only analysis. Maximum-likelihood support was high for the principal internal relationships, with bootstrap values of 93-100% within the lineage comprising *P. xishuangbannaensis*, *P. altebilobata*, *P. menghaiensis* and *P. sumatrana*. *Passiflora xishuangbannaensis* was recovered as sister to *P. altebilobata* (94% BP/1.0 PP), with *P. menghaiensis* successively sister to this pair (100% BP/1.0 PP), followed by *P. sumatrana* (93% BP/1.0 PP). This lineage was not recovered as sister to the other principal lineage, comprising *P. cochinchinensis*, *P. perakensis* and *P. siamica*, which was itself strongly supported (100% BP/1.0 PP; **Figure 3C**). Thus, the mitochondrial phylogeny provided strong resolution of relationships within these principal lineages while recovering a topology markedly different from that inferred from the plastid data.

### Nuclear ribosomal cistron, nrITS and individual-marker phylogenies

The nuclear ribosomal cistron phylogeny provided an independent estimate of relationships among the newly sampled Asian *Passiflora*, but showed substantially lower overall resolution and branch support than the plastid phylogeny. The greatest uncertainty involved the relationships among *P. sumatrana*, *P. altebilobata*, *P. xishuangbannaensis* and *P. menghaiensis*, whose placements also differed from those recovered from the plastid dataset. In contrast, the grouping of *P. cochinchinensis*, *P. perakensis* and *P. siamica* was consistently recovered (100% BP), with strong support for the more terminal relationships **(Supplementary Figure S8)**. The lower resolution of the nuclear ribosomal phylogeny likely reflects the much smaller amount of phylogenetic information provided by a single linked ribosomal locus compared with the concatenated plastid dataset.

The expanded nrITS analysis, incorporating the broader taxonomic sampling available from previous studies, provided considerably stronger resolution for several terminal relationships but little support for many of the deeper branches within supersect. *Disemma*. *Passiflora menghaiensis* was recovered as sister to *P. napalensis* with maximal support (100% BP/1.0 PP). *Passiflora xishuangbannaensis* was placed separately from this pair, similar to the plastid phylogeny but with no support. *Passiflora altebilobata* and *P. kwangtungensis* occurred in the same part of the tree but their relationship was not resolved with meaningful support. Conversely, *P. cochinchinensis*, *P. perakensis* and *P. siamica* remained associated within a comparatively well-supported lineage, providing one of the most consistent phylogenetic signals across the different datasets **(Supplementary Figure S9)**.

The individual marker analyses based on *cytGS*, *ncpGS*, *trnL*-*trnF* and *ndhF* revealed considerable topological variation. The corresponding tanglegram **(Supplementary Figure S10)** showed that only a limited number of terminal relationships were consistently recovered across markers. The *P. cochinchinensis-P. siamica*-*P. perakensis* lineage was relatively stable in the nuclear markers, whereas the *trnL*-*trnF* and *ndhF* phylogenies did not recover the same relationship and displayed substantially greater topological discordance. Numerous additional differences were apparent among deeper relationships, particularly between the nuclear and plastid markers. However, most of these conflicting deeper nodes received very low support, especially in the *trnL*-*trnF* and *ndhF* trees. Consequently, the pronounced crossing pattern observed in the tanglegram should not itself be interpreted as evidence of genuine phylogenetic conflict at these deeper nodes. Rather, much of this apparent discordance reflects the limited phylogenetic signal of individual markers and their inability to resolve deeper relationships confidently. Only conflicts involving independently well-supported relationships can therefore provide meaningful evidence for alternative evolutionary histories.

For *P. menghaiensis*, nrITS, *trnL*-*trnF* and *ndhF* were recovered completely from the annotated assemblies. Recovery of the low copy nuclear markers was more limited because of sequencing depth. For *cytGS*, 463 of 668 bp (69.3%) were supported at ≥3× read depth, with a mean depth of 4.08×, whereas for *ncpGS*, 419 of 514 bp (81.5%) were supported at ≥3× read depth, with a mean depth of 5.78×. No sequence differences from *P. geminiflora* were detected at confidently covered positions in either locus, indicating very high sequence similarity between the two taxa for the recovered portions of these nuclear markers.

### Nuclear gene recovery and phylogenomic analysis

Mapping of the *Passiflora perakensis* paired-end reads to the *Passiflora organensis* reference genome resulted in 47.55% of reads mapping, including 47.06% primary mapped reads, while 43.02% were properly paired. Genome-wide mean sequencing depth was 26.53×. Coverage was uneven across the complete reference, with 50.68% of positions covered at ≥1×, 46.34% at ≥3×, 44.46% at ≥5× and 41.59% at ≥10×. In contrast, annotated protein-coding regions were substantially better represented. Across 31864656 CDS positions, mean sequencing depth was 35.48×, with 96.47%, 95.54%, 94.97%, and 93.74% of positions represented at ≥1×, ≥3×, ≥5× and ≥10×, respectively. Thus, the apparent incompleteness at the whole-genome level largely reflected poorly covered non-coding regions rather than loss of coding information.

Variant calling identified approximately 6.84 million SNPs relative to the *P. organensis* reference. Sample-supported alleles were incorporated into the consensus rather than being removed. Low-depth positions (<5×) and regions affected by indels were masked to N, producing a conservative reference-guided nuclear consensus containing approximately 44.5% unmasked genome-wide sequence.

The *P. organensis* annotation contained 40870 transcripts representing 25327 genes. To prevent alternative transcript isoforms from being treated as independent phylogenetic loci, the longest CDS isoform was retained for each gene. Among these 25327 representative CDS sequences, 21796 (86.1%) contained ≥90% resolved nucleotides, while 20887 (82.5%) contained ≥95% resolved nucleotides. The high recovery of coding sequences despite incomplete genome-wide coverage indicates that the *P. perakensis* resequencing dataset provides extensive nuclear sequence information suitable for gene-based phylogenomic analyses. Locus filtering and alignment

The initial occupancy filter yielded 669 candidate nuclear loci. Sequences containing >50% ambiguous bases (N) were removed at the sequence level, removing 273 sequences and retaining 5079 sequences. Following this step, 35 loci contained six taxa, 203 loci contained seven taxa and 431 loci contained all eight taxa. Six-taxon loci were excluded, leaving 634 loci containing seven or eight taxa. Individual loci were aligned with MAFFT and trimmed with ClipKIT using smart-gap mode.

Phylogenetic informativeness was assessed for each trimmed locus by counting variable and Parsimony Informative (PI) sites while ignoring gaps and ambiguous characters. Of the 634 loci, 541 contained at least one PI site, 445 at least two, 356 at least three, 220 at least five, 65 at least 10 and six at least 20 PI sites. To balance gene-tree informativeness and locus retention, loci with at least three PI sites were retained, resulting in a final dataset of 356 nuclear genes **(Supplementary Table S5)**. Among these, 234 loci contained all eight taxa and 122 contained seven taxa. Taxon representation among the 356 loci was: *Passiflora perakensis*, 356; *P. cochinchinensis*, 355; *P. xishuangbannaensis*, 355; *P. siamica*, 312; *P. menghaiensis*, 349; *P. altebilobata*, 298; *P. sumatrana*, 347; and *P. biflora*, 354.

### Nuclear phylogenetic reconstruction and gene-tree discordance

The ASTRAL-IV analysis recovered strong support for part of the nuclear phylogeny but substantial discordance at several deeper nodes. *Passiflora perakensis* and *P. siamica* formed a maximally supported clade (localPP = 1.0), and their relationship with *Passiflora cochinchinensis* was also maximally supported. The placement of *Passiflora sumatrana* relative to this group was strongly supported in the collapsed gene-tree analysis (localPP approximately 0.97). In contrast, relationships involving *Passiflora xishuangbannaensis*, *P. menghaiensis* and *P. altebilobata* were weakly to moderately supported, including local posterior probabilities of approximately 0.57 and 0.36, indicating substantial among-gene conflict (Figure 4B).

**Figure 4.**
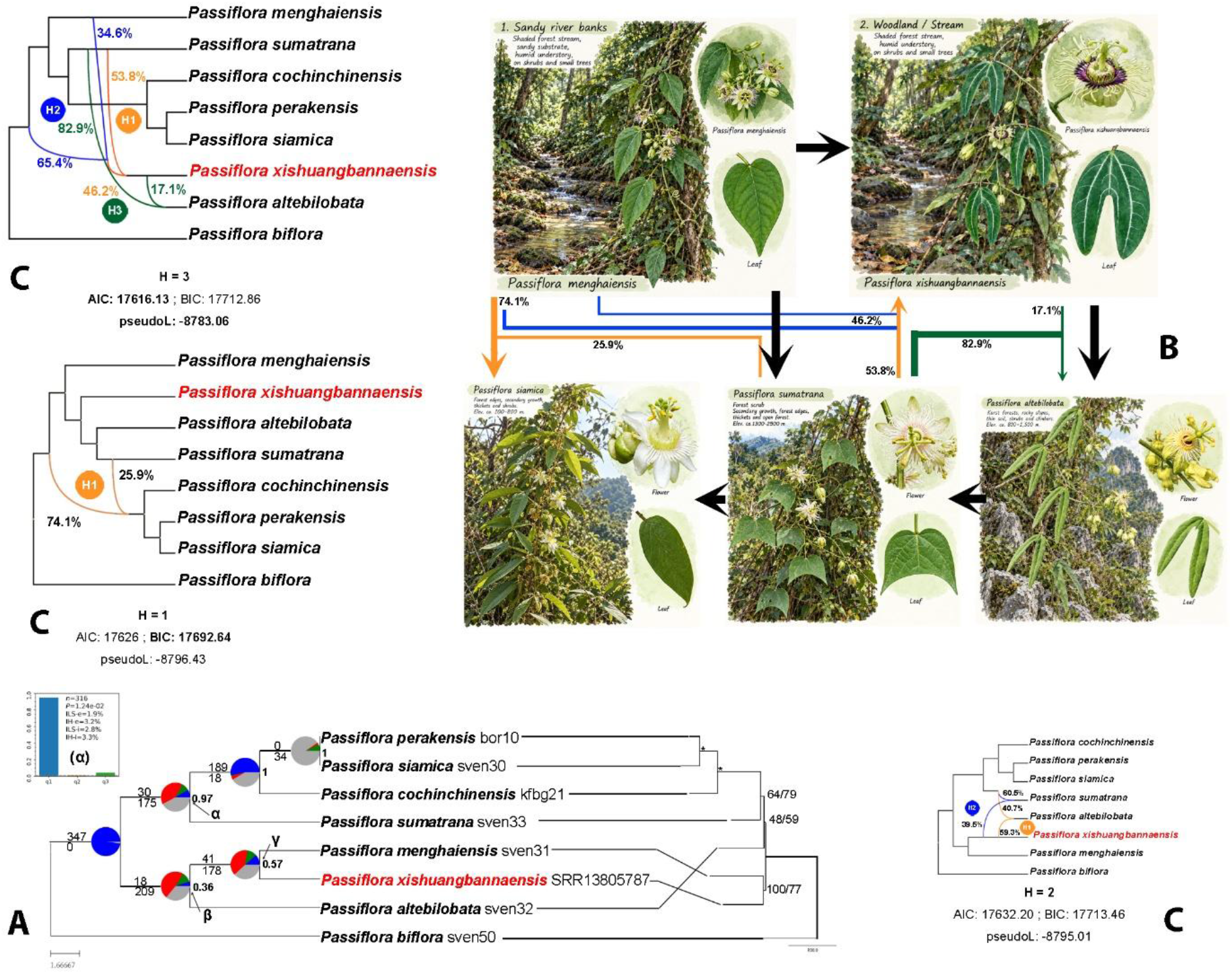
Nuclear phylogenomic conflict, reticulate evolution and ecological differentiation among Asian *Passiflora* supersect. *Disemma* sect. *Octandranthus*. (A) left; ASTRAL species tree inferred from nuclear loci, with gene-tree concordance and discordance summarized by PhyParts; values adjacent to internal branches indicate local posterior probabilities, and pie charts and associated counts summarize concordant and conflicting gene-tree topologies. The inset shows the PhyTop test of asymmetric discordance at the focal α relationship. Right; Maximum-likelihood tree inferred from the concatenated 356 locus nuclear dataset, with SH-aLRT/UFBoot support values shown at internal branches. (B) Botanical panels illustrate ecological and morphological differentiation among representative lineages, from sandy river-bank habitat in *P. menghaiensis* and shaded woodland streams in *P. xishuangbannaensis* to karst outcrops in *P. altebilobata* and forest-scrub/forest-edge habitats in *P. sumatrana* and *P. siamica*, respectively. Arrows indicate the proposed sequence of habitat and morphological differentiation discussed in the text. Botanical habitat panels are AI assisted schematic reconstructions generated with OpenAI image-generation tools using species photographs and habitat information supplied by the authors as morphological and ecological references. The generated illustrations were subsequently reviewed and corrected by the authors for consistency with the observed leaf morphology, floral morphology and habitat. They are intended solely as explanatory illustrations and do not represent primary observational data. (C) PhyloNet networks allowing one to three reticulation events (H = 1-3); coloured reticulation edges indicate inferred hybridization/introgression events and percentages indicate inheritance probabilities. AIC, BIC and pseudo-likelihood scores are given beneath each network. *Passiflora xishuangbannaensis* is highlighted in red.

### Gene-tree concordance and discordance

PhyParts analysis revealed substantial heterogeneity among the individual nuclear gene trees, consistent with the variable support observed in the ASTRAL species tree. The *P. perakensis*-*P. siamica*-*P. cochinchinensis* lineage showed comparatively strong gene-tree concordance, whereas considerably greater conflict was associated with the deeper relationships involving *P. sumatrana*, *P. xishuangbannaensis*, *P. menghaiensis* and *P. altebilobata*. At the α node, corresponding to the placement of *P. sumatrana* relative to the *P. cochinchinensis*-*P. perakensis*-*P. siamica* lineage, 30 gene trees supported the species-tree relationship, while 175 supported conflicting relationships. The β and γ nodes associated with *P. altebilobata* and the *P. xishuangbannaensis*-*P. menghaiensis* relationship also showed discordance: only 18 versus 209 and 41 versus 178 gene trees, respectively, supported the corresponding species-tree and conflicting relationships. These patterns closely paralleled the low ASTRAL local posterior probabilities at β (localPP = 0.36) and γ (localPP = 0.57), compared with strong support at α (localPP = 0.97), (**Figure 4B**).

The concatenated 356 locus, maximum likelihood analysis recovered *Passiflora perakensis* + *P. siamica* with maximal support (SH-aLRT/UFBoot = 100/100), and *P. cochinchinensis* with this pair was also maximally supported (100/100). *Passiflora xishuangbannaensis* and *P. menghaiensis* were recovered together with SH-aLRT/UFBoot support of 100/77. Deeper relationships remained less stable: the node associated with the placement of *Passiflora altebilobata* received weak support (48/59), whereas a *P. sumatrana* received moderate support this time (64/79). Thus, several relationships were strongly supported by the total concatenated nucleotide signal, while the coalescent analysis revealed pronounced discordance among loci at other nodes (**Figure 4B**).

### PhyTop analysis of gene-tree discordance

To further investigate the origin of gene-tree conflict, the relative frequencies of alternative topologies at selected discordant nodes (α–γ) were examined using PhyTop. At node α, 316 informative gene trees were available. The dominant topology accounted for approximately 95% of informative trees, whereas the two alternative topologies occurred at markedly lower and unequal frequencies (approximately 1% and 4%). This asymmetry was statistically significant (P = 0.0124), indicating that the discordance at this node departed from the symmetrical distribution expected under an ILS-only model. The estimated contributions of ILS and introgression/hybridization were low but consistently indicated a greater hybridization/introgression rather than ILS history (ILS-e = 1.9% versus IH-e = 3.2%; ILS-i = 2.8% versus IH-i = 3.3%), (**Figure 4B**). These results provided additional evidence that ILS alone may not fully account for the observed nuclear gene-tree discordance and motivated subsequent explicit phylogenetic-network analyses.

### Phylogenetic network inference

PhyloNet analyses supported increasingly complex reticulate solutions when additional reticulations were permitted. The best H = 1 network had a log pseudo-likelihood of −8796.430, whereas the best H = 2 network had a value of −8795.101. Allowing three reticulations produced a substantially improved maximum pseudo-likelihood of −8783.065. The zero-reticulation analysis provided a poorer fit (−8817.857), indicating that a strictly bifurcating model did not explain the observed distribution of gene-tree topologies as effectively as models incorporating reticulation (**Figure 4C**).

The H = 1 network recovered a single reticulation associated with the otherwise stable lineage containing *P. cochinchinensis*, *P. perakensis* and *P. siamica*. The two parental edges showed strongly asymmetric inheritance probabilities of approximately 74.1% and 25.9%, indicating a predominant ancestry from one lineage accompanied by a smaller contribution from the alternative ancestral lineage.

The H = 2 network recovered two independent reticulation events. The first showed estimated inheritance probabilities of approximately 59.3% and 40.7%, while the second showed contributions of approximately 60.5% and 39.5%. These additional reticulations involved branches within the portion of the network exhibiting substantial gene-tree discordance, including lineages associated with *P. xishuangbannaensis*, *P. altebilobata* and *P. sumatrana*.

The H = 3 network recovered three separate reticulation events, with inheritance probabilities of approximately 53.8/46.2%, 65.4/34.6%, and 82.9/17.1%, respectively. Although the H = 3 network achieved the highest raw pseudo-likelihood, the additional reticulations increased model complexity and therefore cannot individually be regarded as definitive evidence of three separate historical hybridization events. In particular, some of the gene-tree discordance involving *P. xishuangbannaensis* and *P. menghaiensis* had independently shown patterns compatible with ILS.

Network backbone topology also varied with the permitted number of reticulations. Notably, the H = 1 and H = 3 solutions recovered a comparatively basal placement of *P. menghaiensis* that was more consistent with existing taxonomic and morphological evidence than some of the alternative nuclear topologies. This placement is concordant with the comparatively simple leaf morphology as well as habitat preferences of *P. menghaiensis* and floral/stamen characters associated with the *P. perakensis* group. The correspondence between these network solutions and morphology and ecology provides independent support for this arrangement, although the variation among network backbones indicates that these relationships remain sensitive to the modelling of reticulation (**Figure 4A**).

### Comparison of the three GBS analytical strategies

Mapping against *P. edulis* demonstrated that this genome was too divergent to provide an optimal reference for *P. xishuangbannaensis*. Mapping success varied considerably among individuals and was low in representative samples. Only 24.92% of reads from sample 1 mapped to *P. edulis*, with 20.92% properly paired, while sample 17 reached 34.43% total mapping and 30.49% properly paired reads. Thus, approximately two-thirds to three-quarters of reads could fail to contribute to reference-based variant discovery in some individuals.

The reference-free Stacks analysis avoided this limitation and recovered approximately 70000-120000 loci per individual, producing a catalogue of approximately 66804 loci and a large SNP dataset of approximately 256037 SNPs. Importantly, PCA of this independent dataset revealed the same major genetic subdivision subsequently recovered using reference-based analyses.

Reference-based variant calling against the *P. organensis* genome identified 1193185 variant sites across the 39 retained *P. xishuangbannaensis* individuals, including 1058545 SNPs. Restriction to biallelic SNPs yielded 1048361 sites distributed across 321 reference scaffolds. To minimize genotype uncertainty associated with low sequencing depth, individual genotype calls supported by fewer than three reads (DP < 3) were masked as missing rather than removing the corresponding variant site. The agreement between two analytically different approaches indicates that the principal population subdivision was biological rather than an artefact of reference-genome choice. The *P. organensis* dataset was therefore used for the final population genomic analyses.

### Population structure and genetic diversity

PCA based on 4951 SNPs across 39 individuals revealed a pronounced subdivision of *Passiflora xishuangbannaensis* into two major genetic groups (**Figure 5**). ADMIXTURE analysis of the same 4951 SNP dataset likewise supported a primary division into two genetic clusters. STRUCTURE analysis, conducted using a more stringently LD-pruned subset of 681 SNPs across the same 39 individuals, independently corroborated this pattern. Across ten replicate STRUCTURE runs for each K from 1 to 6, the Evanno ΔK statistic showed a pronounced maximum at K = 2 (ΔK = 252.60), compared with substantially lower values at K = 3 (4.26), K = 4 (0.99), and K = 5 (21.23), **(Supplementary Figure S11)**. Thus, the analyses consistently supported two principal genetic populations, although K = 3 revealed additional finer-scale structure.

**Figure 5.**
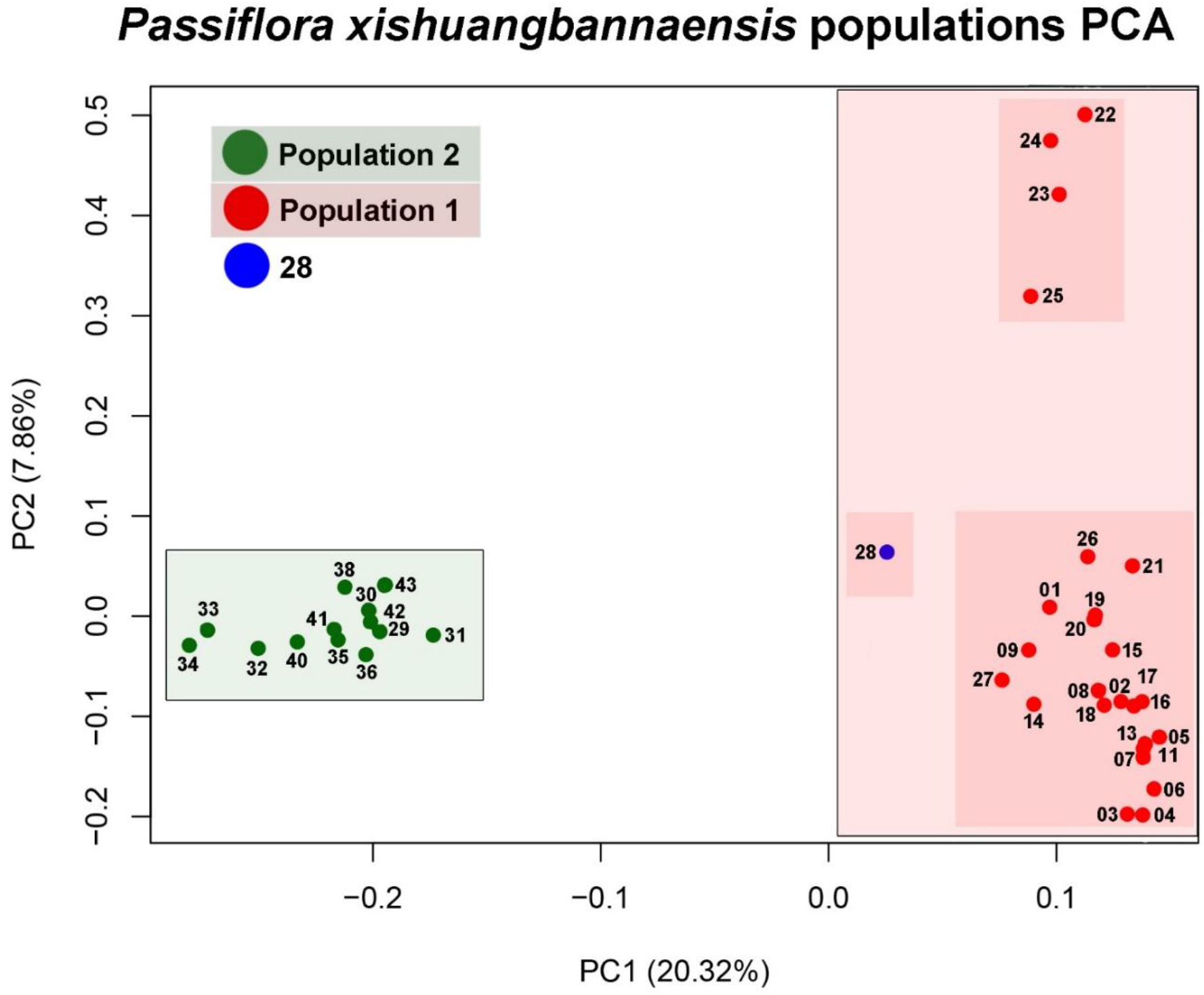
Principal component analysis (PCA) of genetic variation among 39 *Passiflora xishuangbannaensis* individuals. Individuals form two clearly differentiated genetic groups corresponding to Population 1 and Population 2, primarily separated along PC1, which explains 20.32% of the total variation. PC2 explains an additional 7.86% and reveals some within-population variation. Individual 28 occupies an intermediate position between the two principal genetic groups and is shown separately.

The two major populations were genetically differentiated, with a genome-wide *F_ST_* of approximately 0.112, indicating appreciable differentiation between the two geographic/genetic groups. Genetic diversity also differed between populations. Population 1 showed higher diversity (Na = 1.8868, Ne = 1.4159, H = 0.2443, I = 0.3755, PPL = 88.7%) than population 2 (Na = 1.6171, Ne = 1.3156, H = 0.1865, I = 0.2859, PPL = 61.7%), (**Table 2**).

**Table 2.** Genetic diversity and inbreeding estimates for the two populations of *Passiflora xishuangbannaensis*. Number of individuals, mean number of observed alleles (Na), effective number of alleles (Ne), Nei’s gene diversity (H), Shannon’s information index (I), mean PLINK inbreeding coefficient (F), and percentage of polymorphic loci (PPL) are shown for each population.

| Population | Individuals<br>total | $N_a$ | $N_e$ | $H$ | $I$ | $F$ | PPL |
| --- | --- | --- | --- | --- | --- | --- | --- |
| pop1 | 26 | 1.8868 | 1.4159 | 0.2443 | 0.3755 | 0.125 | 88.7 |
| pop2 | 13 | 1.6171 | 1.3156 | 0.1865 | 0.2859 | 0.136 | 61.7 |

Genome-wide PLINK estimates indicated positive mean inbreeding coefficients in both major populations. Mean individual F was 0.125 in population 1 (SD = 0.090; range -0.066-0.314) and 0.136 in population 2 (SD = 0.135; range 0.013-0.432). Population 2 therefore showed slightly higher mean inbreeding and substantially greater among-individual variation in F, including particularly high values in samples 41 and 42 (F = 0.392 and 0.432, respectively).

Analysis of MOlecular VAriance (AMOVA), based on 36470 LD-pruned SNPs from 38 individuals, revealed significant genetic differentiation between the two populations. Among-population differences accounted for 21.04% of the total molecular variance, whereas 78.96% occurred within populations (**Table 3**). The corresponding ΦST was 0.210 and was significant based on 999 permutations (P = 0.001). These results support substantial genetic structuring between the two geographically defined populations of *Passiflora xishuangbannaensis*, while indicating that most genetic variation remains distributed among individuals within populations.

**Table 3.** Analysis of molecular variance (AMOVA) between the two principal populations of *Passiflora xishuangbannaensis*. The analysis was based on 36470 LD-pruned SNPs from 38 individuals assigned to Pop1 (n = 25) and Pop2 (n = 13). Sample 28, which showed intermediate ancestry, was excluded. Genetic variation was partitioned into among and within population components. Population differentiation was significant (ΦST = 0.210; P = 0.001, 999 permutations). df, degrees of freedom; SS, Sum of Squares; MS, Mean Squares; Est. Var., Estimated Variance component.

| Source | df | SS | MS | Est.Var. | % |
| --- | --- | --- | --- | --- | --- |
| Among populations | 1 | 0.808273 | 0.808273 | 0.038752 | 21.04% |
| Within populations | 36 | 5.234826 | 0.14541 | 0.145412 | 78.96% |
| Total | 37 | 6.043099 | - | 0.184164 | 100.00% |

Taken together, PCA, ADMIXTURE, STRUCTURE, *F_ST_* and diversity statistics demonstrate substantial population structure within *P. xishuangbannaensis*, with population 2 exhibiting lower overall genetic diversity than population 1.

### Geographic distribution of genetic structure

Spatial visualization of ancestry coefficients revealed a strong geographic component to the genetic structure of *Passiflora xishuangbannaensis*. Under the K = 2 model, most individuals were strongly assigned to one of the two principal ancestry components, consistent with the major genetic subdivision detected independently by PCA and the population-structure analyses. Nevertheless, several individuals exhibited mixed ancestry, indicating that genetic differentiation was not absolute and suggesting either historical or ongoing gene flow between the principal genetic groups.

Mapping the K = 3 model provided additional resolution of fine-scale genetic variation. Although most individuals continued to show strong assignment to a predominant ancestry component, a subset contained appreciable contributions from a second or third component. Importantly, this variation occurred over a very restricted geographic scale, with genetically differentiated or admixed individuals occurring in close spatial proximity (**Figure 6**).

**Figure 6.**
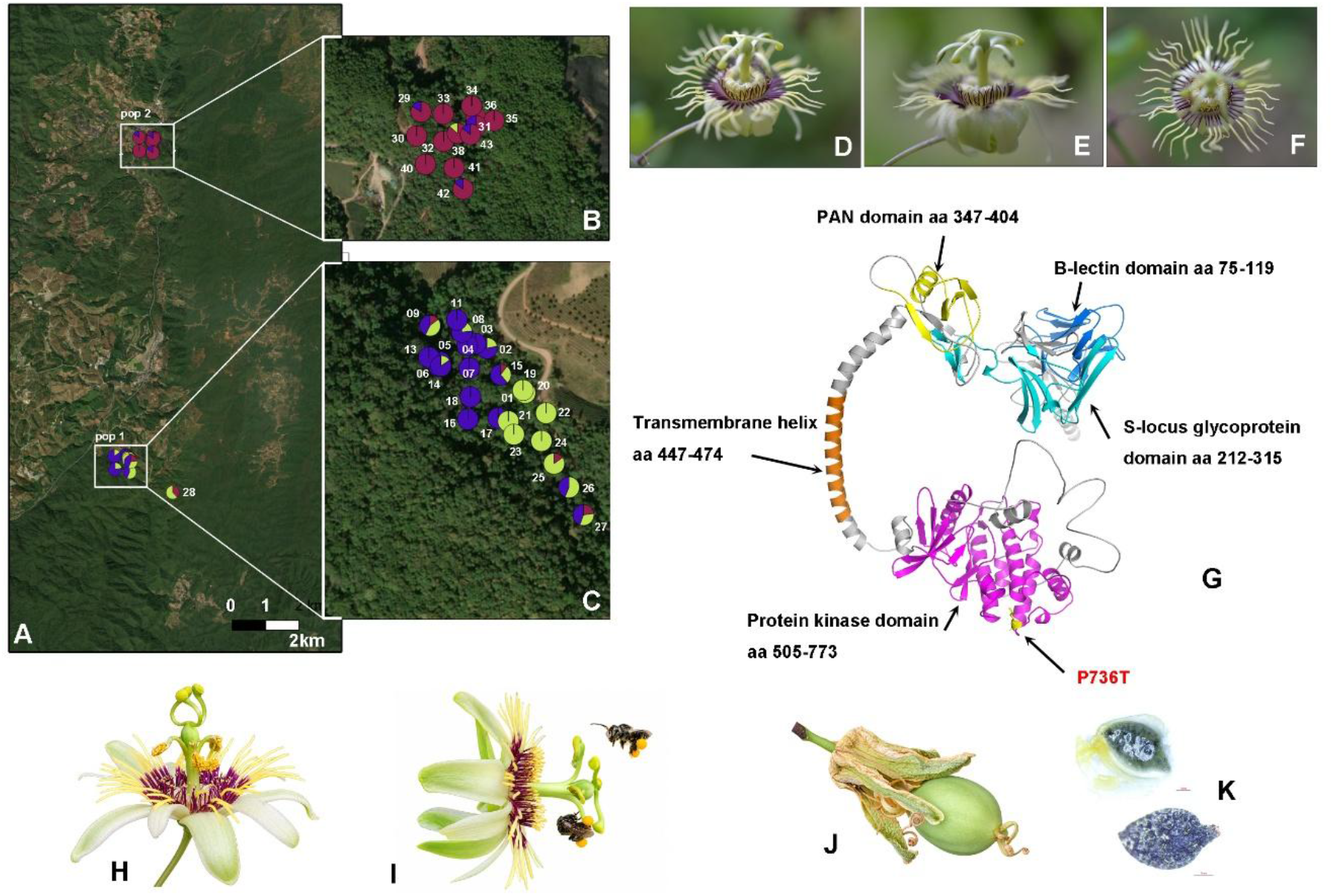
Population structure, floral biology and candidate self-incompatibility receptor in *Passiflora xishuangbannaensis*. (A–C) Geographic distribution and genetic structure of the two identified populations of *P. xishuangbannaensis*, with individual ancestry proportions shown as pie charts generated from the population-structure analysis and mapped in QGIS; enlarged views show populations 2 (B) and 1 (C). (D–F) Flowers of *P. xishuangbannaensis*, illustrating variation in floral aspect. (G) AlphaFold2/ColabFold-predicted structure of the candidate G-type lectin S-receptor-like kinase gene2019. Pfam-identified B-lectin (aa 75–179), S-locus glycoprotein (aa 212–315), PAN (aa 347–404), transmembrane (aa 447–474), and protein kinase (aa 505–773) regions are indicated; the population-differentiated P736T substitution is highlighted within the kinase domain. (H) Flower showing the stigmas held in an upward position. (I) Stingless bees visiting the flower; these small visitors were observed but considered ineffective pollinators. (J) Fruit produced following hand pollination. (K) Seed and associated aril. Photos sven Landrein.

The geographic pie-chart analysis therefore demonstrates that the genomic structure identified by PCA and ancestry analyses is spatially organized but cannot be explained simply by large geographic distances between sampling localities. Instead, substantial differences in ancestry composition occur among individuals separated by relatively short distances within the sampled landscape. The K = 3 map further illustrates the finer-scale subdivision that was obscured under K = 2, while retaining the same overall pattern of a dominant genetic division with localized admixture (**Figure 6**).

### Clone detection and pairwise genetic relatedness

Initial PLINK identity-by-descent analysis produced relatively high PI_HAT estimates for numerous pairs of individuals. The highest values included s19/s20 (PI_HAT = 0.7488), s11/s13 (0.7370), s16/s17 (0.7214), s17/s18 (0.7117), s03/s04 (0.7053), s40/s41 (0.7052), and s22/s23 (0.7051). Previously suspected highly similar pairs, s19/s21 and s22/s24, had PI_HAT values of 0.6908 and 0.6969, respectively, while s06/s07 showed a comparable value of 0.6897. The occurrence of similarly elevated PI_HAT values across numerous pairs, together with the high level of missing data in the GBS dataset, indicated that these estimates should not be interpreted directly as evidence of clonality.

Pairwise genotype concordance provided a more direct assessment by restricting each comparison to SNPs successfully genotyped in both individuals. No pair exhibited complete or near-complete genotype identity. The highest concordance was observed between s19 and s20, which shared genotype calls at 2338 SNPs, of which 2073 were identical and 265 differed, corresponding to 88.67% concordance. Other highly similar pairs included s11/s13 (88.48%), s03/s04 (88.05%), s22/s23 (87.85%), s16/s17 (87.45%), s17/s18 (87.45%), s22/s24 (87.31%), s40/s41 (87.22%) and s19/s21 (87.20%), **(Supplementary Figure S12)**.

The distribution of pairwise concordance showed that most comparisons had substantially lower similarity, broadly concentrated at approximately 55-75%, whereas a smaller group of pairs occurred at approximately 82-89% concordance. Thus, a subset of individuals was substantially more genetically similar than the majority of sampled pairs, consistent with close relatedness and/or fine-scale genetic structure (**Figure 7**).

**Figure 7.**
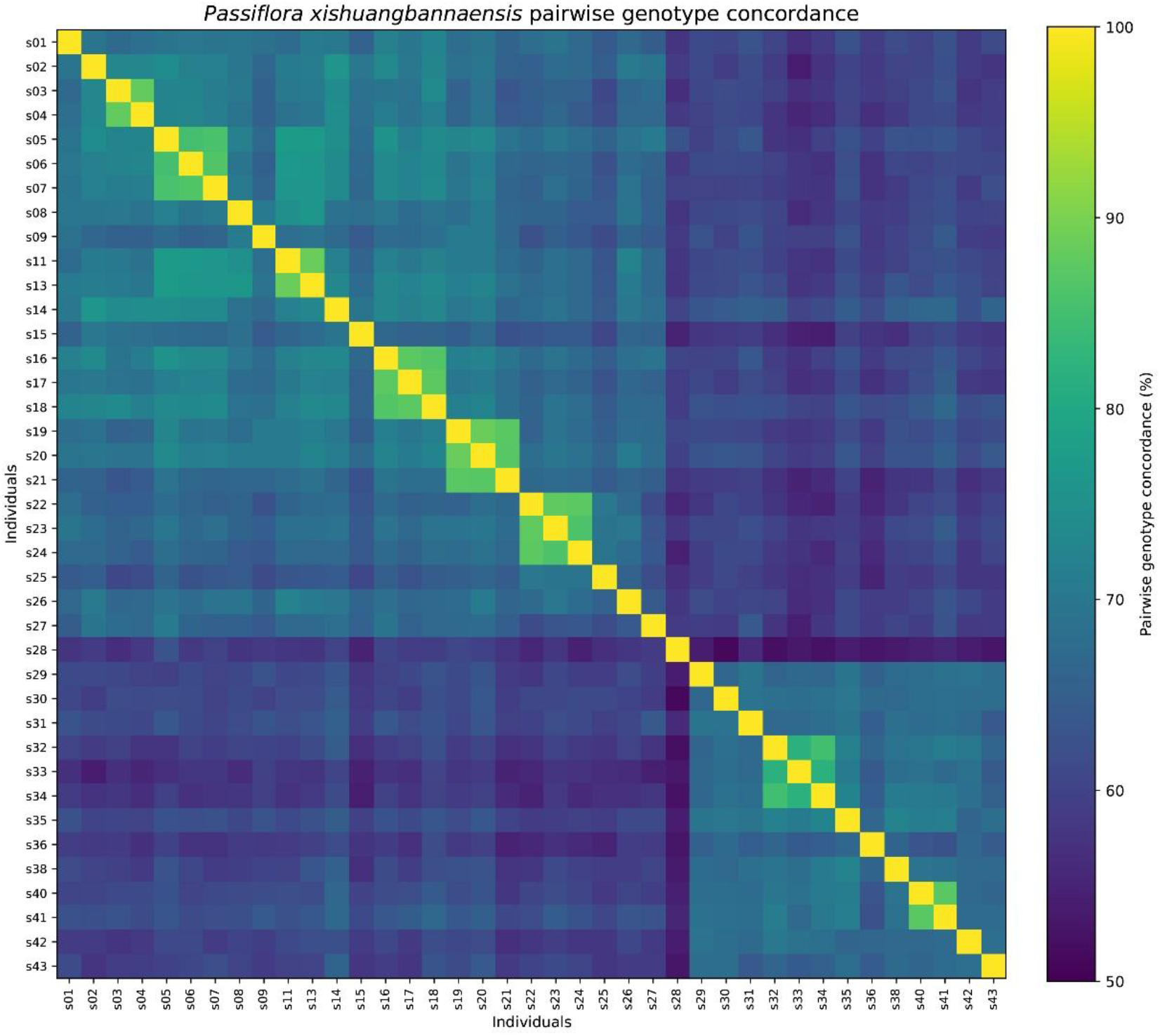
Pairwise genotype concordance among *Passiflora xishuangbannaensis* individuals. Heatmap showing pairwise genotype concordance (%) calculated from the GBS dataset. Warmer colours indicate greater genetic similarity between individuals. Several clusters of highly concordant individuals occur within populations, whereas concordance is generally lower between the two major genetic populations, consistent with pronounced population structure and fine-scale genetic relatedness within populations.

Importantly, however, even the most similar pair differed at 265 of 2338 jointly genotyped SNPs (11.3%). The genomic data therefore provided no evidence that any of the sampled individuals represented genetically identical clones or duplicate genotypes. Consequently, the previously suspected pairs s19/s21 and s22/s24 should not be classified as clones, and the 39 sampled individuals can be treated as distinct multilocus genotypes. The elevated similarity observed among several local groups instead suggests substantial relatedness among some individuals, potentially reflecting family structure or restricted local gene flow within populations.

### Recent demographic history

The four chromosomes GONE2 analysis recovered evidence of a substantial decline in recent effective population size in both populations, although the timing and shape of the decline differed markedly between them.

For population 1, five independent GONE2 runs converged closely on a demographic history characterized by an approximately stable historical effective population size of around 1070-1150 individuals between approximately 10 and 40 generations before present, followed by a pronounced recent contraction. Across replicate runs, *N*_e_ was approximately 1090-1155 at generation 10, 1100-1151 at generation 20, and 1067-1102 at generation 40. In contrast, estimated *N*_e_ declined to approximately 414-534 at generation 5 and only 105-109 at generation 1. Thus, the predominant population 1 solution indicates a particularly strong contraction occurring within approximately the last 5 to 10 generations (**Figure 8**).

**Figure 8.**
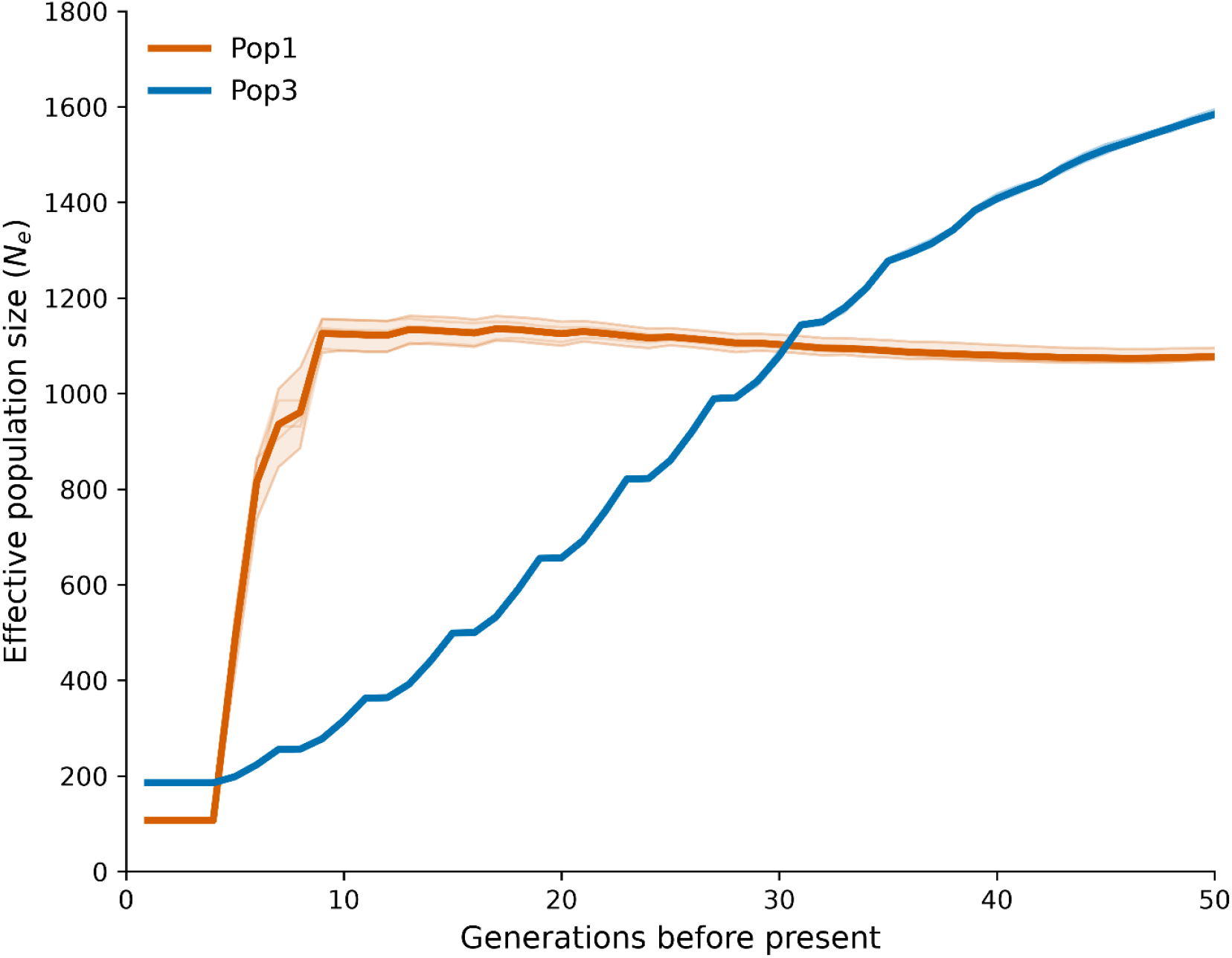
Recent demographic history of *Passiflora xishuangbannaensis* populations inferred using GONE2. Effective population size (*N*e) was reconstructed separately for Pop1 and Pop2 from linkage disequilibrium among GBS-derived SNPs mapped to the *P. organensis* reference genome. Analyses used 5402 SNPs for Pop1 and 5876 SNPs for Pop2 distributed across four reconstructed chromosome groups, a recombination rate of 1.0 cM Mb^-1^, and unphased diploid genotypes. Lines represent mean *Ne* estimates from five independent GONE2 runs with different random seeds, with generations before present increasing from left to right. Both populations show declining *Ne* toward the present, with a particularly abrupt recent decline in Pop1 and a more gradual decline in Pop2.

The demographic trajectory of population 2 was more gradual and exceptionally consistent among replicate GONE2 runs. Effective population size was approximately 1401-1418 at generation 40, declining to 1071-1091 at generation 30, 654-658 at generation 20, 313-322 at generation 10 and 197-199 at generation 5. The most recent estimate was only 184-187 individuals at generation 1. Population 2 therefore showed a progressive decline in *N*_e_ throughout the reconstructed recent demographic interval rather than the abrupt recent contraction inferred for population 1 (**Figure 8**).

Importantly, the five independent random seed replicates produced nearly identical estimates for population 2 and strongly concordant estimates for population 1, demonstrating that the principal demographic patterns were reproducible across stochastic GONE2 optimizations. One preliminary population 1 run using a different random seed (12345) converged on an alternative, smoother trajectory, indicating some optimization uncertainty for this population; however, all five subsequent independent replicates converged on the solution with a sharp recent decline.

Analyses based initially on only scaffolds 1 and 2 were substantially less stable, including an anomalously low population 1 estimate during the first four generations followed by an abrupt increase in estimated *N*_e_. Incorporation of the additional chromosome arm information for chromosomes 5 and 6 increased the analysed marker set from approximately 3200-3500 SNPs to 5402 SNPs in Pop1 and 5876 SNPs in population 2 and produced substantially more coherent demographic reconstructions. Sensitivity analyses further showed that removal of closely related population 1 individuals did not eliminate the inferred recent decline, indicating that the signal was not driven solely by the two highly related sample pairs.

Overall, the LD-based analyses therefore support recent reductions in effective population size in both geographically and genetically differentiated populations of *P. xishuangbannaensis*. Population 1 is characterized by an approximately stable older *N*_e_ followed by a sharp recent contraction, whereas population 2 exhibits a more gradual decline extending across several tens of generations. Despite their contrasting trajectories, both populations reached low inferred recent effective population sizes, suggesting substantial recent demographic reduction across the species sampled range.

In population 1 and 2, the Stairway Plot reconstruction indicated a long term decline beginning approximately 85000 years ago rather than a single abrupt population crash **(Supplementary Figure S13)**.

### Genetic variation in candidate self-incompatibility genes

Sequence reconstruction supported .2019 as an intact G-type lectin S-receptor-like kinase candidate. The reconstructed 619 bp CDS encoded an approximately 873 aa protein without internal stop codons and closely matched the 872 aa *Passiflora organensis* prediction, with the characteristic B-lectin-SLG-PAN-TM-kinase architecture. The differentiated SNP mapped to CDS position 2206 (genomic position 9912928 on the reverse strand of scaffold 7), where CCA becomes ACA produces a non-synonymous P736T substitution within the C-terminal kinase domain. AlphaFold modelling of residues 560-810 produced a well folded kinase domain for both alleles, with similar confidence (pTM = 0.85 for P736 and 0.86 for T736), no structural clashes, and a core Cα RMSD of 0.378 Å, indicating no major disruption of the kinase fold. The substitution nevertheless altered the local physicochemical environment, with the introduced Thr hydroxyl positioned approximately 3.25 Å from the Lys737 backbone N, leaving open the possibility of a localized functional effect. Population GBS coverage was insufficient to evaluate the complete receptor: only 243 of 2113 CDS positions were covered at ≥3× in ≥60% of both populations, corresponding approximately to residues 566-646 within the kinase region. Consequently, variation in the extracellular recognition domains could not be reliably assessed, and the functional significance of P736T remains to be experimentally established (**Figure 6**).

## DISCUSSION

### Reticulate evolution and the limits of a single species tree

The combined plastid (**Figure 1**), nuclear ribosomal **(Supplementary Figure S8)**, concatenated nuclear (**Figure 4**), multispecies-coalescent and network analyses (**Figure 4**) reveal an evolutionary history in Asian *Passiflora* supersect. *Disemma* sect. *Octandranthus* that cannot be reduced to a single, fully bifurcating tree. Some relationships were remarkably stable. In particular, *P. siamica*, *P. perakensis* and *P. cochinchinensis* formed a consistently supported lineage across plastid, nuclear ribosomal, concatenated and coalescent analyses. In contrast, the positions of *P. xishuangbannaensis*, *P. menghaiensis*, *P. altebilobata* and *P. sumatrana* changed among genomic compartments and analytical frameworks, and the corresponding nuclear nodes showed low concordance among gene trees. This contrast between a stable terminal lineage and a highly discordant set of short internal branches is characteristic of rapid diversification in which Incomplete Lineage Sorting (ILS) and interspecific gene flow have acted together (Maddison, 1997; Degnan & Rosenberg, 2009; Folk et al., 2017, 2018).

The distinction between these processes is important. ILS alone is expected to generate alternative gene-tree topologies around short internodes, and under the simplest multispecies-coalescent model the two minor topologies should occur at approximately equal frequencies. At the focal alpha node, however, the alternative topologies were significantly asymmetric (P = 0.0124), and PhyTop attributed a slightly larger component of discordance to introgression or hybridization than to ILS. PhyloNet likewise fitted the gene-tree distribution better when reticulation was permitted than under a zero-reticulation model. Together, these results reject an ILS-only explanation for all of the observed discordance. They do not, however, establish every reticulation edge as a unique historical hybridization event. The H = 3 model had the highest raw pseudo-likelihood, but additional reticulations inevitably improve fit and may absorb gene-tree error, unsampled lineages or other departures from the model. The H = 1 is therefore biologically informative alongside H = 3, rather than merely inferior versions of a definitive three-reticulation history (**Figure 4**).

This interpretation also explains why previous analyses based on nrITS and a few plastid or low-copy nuclear markers could resolve the limits of supersection *Disemma* and some terminal relationships but not the deepest relationships within section *Octandranthus* **(Supplementary Figure S10)**, (Krosnick & Freudenstein, 2005; Krosnick, 2006; Krosnick et al., 2013). A short locus may recover one of several legitimate genealogical histories, whereas concatenation can yield strong bootstrap support for the dominant aggregate signal even when many loci support alternatives. In the present study, the concatenated analysis strongly united *P. xishuangbannaensis* and *P. menghaiensis*, but the corresponding ASTRAL branch had only moderate local posterior support and strong gene-tree conflict (**Figure 4**). The apparent precision of a concatenated tree should consequently not be equated with historical simplicity (Degnan & Rosenberg, 2009). Conversely, weak support in the coalescent tree is biologically meaningful here: it identifies branches for which the sampled loci do not converge on a single history.

Cytonuclear discordance is especially plausible in *Passiflora* because plastid inheritance is unusually variable. Maternal, paternal and biparental transmission have all been demonstrated, and subgenus *Decaloba* frequently shows maternal or biparental inheritance followed by sorting of heteroplasmy during development (Muschner et al., 2006; Hansen et al., 2007; Shrestha et al., 2021). A mature plant may therefore retain only one plastid lineage after hybridization, while its recombining nuclear genome preserves contributions from both parental lineages. Organellar capture can then produce a well-supported plastid topology that differs from the predominant nuclear history, even when neither analysis is technically erroneous (Rieseberg & Soltis, 1991; Folk et al., 2017). The strong plastid association of *P. xishuangbannaensis* with *P. menghaiensis*, coupled with weaker and conflicting nuclear support, is consistent with this mechanism, although it does not identify the direction or timing of gene flow.

Reticulation itself does not vanish through time, but its recoverable signature can become obscured. Recombination breaks introgressed tracts into progressively smaller segments, ancestral alleles sort or are lost, and extinction or incomplete sampling removes potential parental lineages. Deep phylogenetic nodes may therefore appear cleaner than recent ones because only the dominant surviving signal remains, not because early evolution was non-reticulate. This provides a plausible explanation for why plastid data can resolve many deeper relationships in *Passiflora* while the recent Asian radiation remains difficult. It also answers a central question raised by these results: the alternative trees are not competing estimates from which one must simply select the ‘real’ phylogeny. For the stable *P. siamica*-*P. perakensis*-*P. cochinchinensis* lineage, a bifurcating representation is well supported. For the *P. xishuangbannaensis*-*P. menghaiensis*-*P. altebilobata*-*P. sumatrana* complex, the most realistic current representation is a set of strongly supported terminal relationships embedded in a reticulate and incompletely sorted history.

### Reference-guided phylogenomics for poorly sampled tropical lineages

The recovery of 356 informative nuclear genes from existing Illumina resequencing data demonstrates a practical route for phylogenomics in non-model plants **(Supplementary Table S5)**. Coding regions were recovered much more completely than the genome as a whole, and stringent depth masking, isoform selection and locus filtering produced a dataset capable of distinguishing stable branches from highly discordant ones. This strategy complements target-enrichment approaches such as Hyb-Seq and Angiosperms353, which offer standardized orthologous markers and efficient cross-study integration (Weitemier et al., 2014; Johnson et al., 2019). The similarity between 356 recovered loci and the 353 loci of Angiosperms353 is coincidental: the present genes were selected empirically from the *P. organensis* annotation according to coverage, occupancy and phylogenetic informativeness, whereas Angiosperms353 was designed from conserved single-copy orthologues sampled across angiosperms.

Reference-guided recovery is particularly valuable where probe kits, high-molecular-weight DNA or new field collections are difficult to obtain, as is often the case for rare tropical species. It also permits archived whole genome reads to be repurposed for phylogenemics. Its limitations are equally important: mapping favours regions similar to the reference, paralogues may be collapsed, divergent alleles may be missed, and low-coverage taxa contribute incomplete sequences. The method should therefore be presented as a cost effective bridge to denser genomic sampling, not a substitute for chromosome scale assemblies, long reads or purpose designed target capture. Adding *P. napalensis*, *P. leschenaultia* DC., *P. kwangtungensis* and broader population sampling of the focal species will provide the most decisive next test of the network hypotheses.

### Plastome restructuring as an additional record of lineage history

The plastome comparisons reinforce the exceptional structural dynamism already reported in *Passiflora*, especially subgenus *Decaloba* (Cauz-Santos et al., 2020, 2025). All sampled *Disemma* shared a reduced set of 105 unique genes and losses or pseudogenization affecting *accD*, *rpl20*, *rpl22*, *rpl32*, *rps7*, *rps16* and *ycf1/ycf2*, supporting a common structural background for the group (**Figure 3**). Within that background, however, repeated inversions and changes in IR boundaries generated marked differences in genome architecture without further changes in unique gene number. The results therefore distinguish older synapomorphic gene content changes from younger, lineage-specific rearrangements.

The most striking example was *P. sumatrana*, whose approximately 46.7kb IRs extended into the *accD*-*psaI* interval and reduced the LSC to only 57.4 kb. This plastome is almost 30 kb longer than those of most sampled *Octandranthus* despite retaining the same number of unique genes. At the other extreme, *P. cinnabarina* possessed strongly contracted IRs and the smallest plastome examined (**Table 1**). These observations extend earlier demonstrations that IR expansion, contraction and even loss have occurred repeatedly within *Passiflora* (Cauz-Santos et al., 2020). They also show why plastome length alone is a poor proxy for gene content or evolutionary advancement: genome size here primarily records changes in duplicated boundaries and arrangement.

Structural similarity should nevertheless be interpreted independently of plastid-tree identity. Large rearrangements can provide valuable rare genomic characters, but parallel boundary shifts are possible, and the plastome is inherited as a single linked unit. In a group with biparental transmission and organellar capture, an entire plastome structure can move between species together with its nucleotide genealogy. The distinctive *P. sumatrana* expansion may therefore be a useful diagnostic feature of that plastid lineage, but its distribution must be tested across additional individuals and related species before it is treated as a synapomorphy of a taxonomic group. The comparatively conservative *Adenia* plastomes provide an informative contrast, although the *rps19*-*rpl2* boundary contraction in *A. penangiana* demonstrates that lineage-specific IR change is not confined to *Passiflora* (**Table 1**).

### Biogeography, ecology and morphological evolution across the Hua Line

Southern Yunnan lies at a major floristic junction among tropical Asian, Indo-Himalayan, Indochinese and southern Chinese elements (Zhu, 1997, 2008, 2012). The Hua Line, broadly following the Lixianjiang river and associated geological transition, was proposed to distinguish the floras of southern and southeastern China (Zhu, 2011). The distribution of *Octandranthus* does not conform to an impermeable boundary (**Figure 2**): Although several lineages cross the Hua Line, the more recently evolved *P. siamica*-*P. cochinchinensis*-*P. perakensis* lineage closely follows this biogeographic boundary, spanning a broad distribution across the Indo-Malayan region and southern China. The concentration east of the line of *P. menghaiensis* and P. *xishuangbannaensis*, together with their phylogenetic connections to lineages occurring farther west or south, suggests that this regional boundary may have structured dispersal and secondary contact.

Because the present phylogeny was not time calibrated, the observed distributions cannot be assigned directly to a particular tectonic event or dated episode of river reorganization. The Hua Line should therefore be treated as a biogeographic hypothesis rather than a vicariant event demonstrated by our data. Wider sampling, especially *P. napalensis*, *P. leschenaultii* and the recently rediscovered *P. kwangtungensis*, will be essential. These taxa connect the Xishuangbanna endemics to the Indian-Himalayan and southern Chinese floras and may represent either unsampled parental lineages or surviving relatives of ancestral populations.

The network backbones that place *P. menghaiensis* relatively early within the sampled Asian radiation are compatible with its simple, thin, cartilaginous leaves and river-bank ecology, but this does not make the extant species an ancestor of the others (**Figure 4**). An extant terminal taxon can retain ancestral looking characters while also possessing its own derived history. Simple leaves may be plesiomorphic, secondarily derived or maintained by habitat-associated selection; only explicit ancestral state reconstruction with much broader taxon sampling can distinguish these alternatives. The same caution applies to the eight stamens and variable stigma or carpel number found in the *P. siamica*-*P. perakensis* group. Floral-organ number is a plastic character within Asian *Passiflora* (Krosnick et al., 2006), so it is not sufficient on its own to justify a new subsection. The stable genomic support for this lineage is taxonomically promising, but formal recognition would require expanded sampling, diagnosis and nomenclatural treatment beyond the scope of this study.

Ecological differentiation nevertheless offers a useful set of testable hypotheses. *P. menghaiensis* and morphologically similar species occupy sandy riverbanks and stream beds; *P. xishuangbannaensis* is rhizomatous and occurs in riverine woodland but generally farther from the active channel; *P. altebilobata* is associated with karst outcrops; and *P. siamica* and *P. sumatrana* are more vigorous vines of forest margins or scrub (Wang et al., 2007). Leaf dissection and rhizome development may therefore have evolved repeatedly in association with habitat, disturbance and seasonal persistence rather than tracking a simple linear series of species divergence. The unusual bilobed leaves in this group, often interpretable as trilobed leaves with a strongly reduced terminal lobe, are particularly suitable for comparative developmental study. Candidate-gene speculation alone would be premature; transcriptomic comparison of shoot apices, developing leaves and rhizomes across closely related simple and lobed leaves species would provide a stronger route to identifying the underlying developmental pathways.

### Population structure and demographic decline in *Passiflora xishuangbannaensis*

The population genomic results confirm that *P. xishuangbannaensis* is genetically structured despite its extremely narrow known distribution. PCA, ADMIXTURE and STRUCTURE independently recovered two principal populations, with K = 2 strongly preferred and K = 3 revealing finer substructure (**Figure 5 & 6)**. Genome-wide *F_ST_* was approximately 0.112, AMOVA attributed 21.04% of molecular variance to differences between populations, and population 2 retained lower gene diversity and fewer polymorphic loci (**Table 2 & 3)**. Agreement between the de novo Stacks analysis and mapping to *P. organensis* is particularly important because it shows that the main subdivision is not simply an artefact of mapping to a divergent reference.

Individual 28 and several mixed-ancestry individuals show that differentiation is incomplete. Because appreciable changes in ancestry occur over short geographic distances, the structure cannot be explained by isolation by distance alone. Restricted pollen or seed movement, fine-scale habitat discontinuities and historical fragmentation are all plausible. The present SNP dataset, however, cannot by itself partition pollen mediated from seed mediated gene flow. Such inference requires comparison of markers with contrasting modes of inheritance, typically biparentally inherited nuclear markers and uniparentally inherited organellar markers (Ennos, 1994; Petit et al., 2005). That comparison would require maternally or paternally inherited organellar haplotypes analysed alongside biparentally inherited nuclear markers, with the additional complication that plastid inheritance in *Decaloba* can be biparental. Chloroplast SNPs recovered from deeper sequencing could therefore be informative, but their inheritance mode should be verified in *P. xishuangbannaensis* before they are used as a purely seed dispersal marker.

The absence of identical multilocus genotypes is also biologically informative. All 39 sampled plants were distinct, even though some pairs were closely related and the species is rhizomatous. Vegetative spread therefore did not dominate the sampled stands to the point of producing repeated ramets with identical GBS profiles (**Figure 7**). This does not mean that rhizomatous propagation is unimportant: somatic mutation, genotyping error and unsampled connections between shoots can complicate clone detection, and the sampled plants may represent genets selected visually in the field. Nevertheless, sexual recruitment or historical sexual reproduction has contributed substantially to the extant genetic composition. The combination of distinct genotypes, close local relatives and strong population structure is consistent with restricted dispersal and clustering rather than pervasive clonality.

Both demographic methods indicate decline, but at different temporal scales. Stairway Plot suggested a long-term reduction beginning approximately 85 ka, whereas GONE2 inferred very recent declines in both populations, with a sharp contraction in population 1 during the last five to ten generations and a more gradual decline in population 2 (**Figure 8 & Supplemementary Figure S13)**. GONE was developed specifically to infer recent *N*_e_ from the distribution of linkage disequilibrium (Santiago et al., 2020), and the agreement among independent runs and sensitivity analyses strengthens the qualitative conclusion.

Absolute estimates should nonetheless be treated cautiously. The analysis relied on chromosome arm scaffolds rather than a complete chromosome assembly, an assumed recombination rate and generation time, and unequal population sample sizes. Population structure, linked marker placement and overlapping generations can also affect LD based inference. The most defensible conclusion is therefore not the exact *N*_e_ at a particular generation, but the concordant evidence that both populations have contracted and now retain low recent effective sizes.

Field observations provide an ecological context for this decline. Much of the valley landscape has been converted or fragmented by agriculture, roads and associated development. Population 2 occurred under dense cover and no flowering was observed, whereas larger and more floriferous plants were sometimes found near road or field edges in population 1. This pattern need not contradict habitat degradation.

Moderate canopy opening may temporarily increase light and flowering in an forest margin adapted vine, while the same land-use change reduces habitat continuity, pollinator movement, recruitment sites and long-term population size. A short-term reproductive response to increased light can therefore coexist with a long-term demographic cost of fragmentation. Quantitative monitoring of survival, flowering, fruit set, canopy openness and microclimate is needed before the forest margin habitat is interpreted as beneficial.

### Self-incompatibility and conservation management

The reproductive observations and genomic data identify mate availability as a major conservation concern, but they do not yet resolve the molecular SI system of *P. xishuangbannaensis*. Self pollination repeatedly failed, some controlled crosses produced fruit, and few fruits were seen in the field. In small SI populations, loss of S-allele diversity can reduce the number of compatible mates even when genome-wide diversity remains measurable, producing an S-Allee effect in which declining population size further reduces seed set (Busch & Schoen, 2008; Young & Pickup, 2010). The differentiation between the two populations makes interpopulation crosses a rational conservation experiment because they may combine compatibility classes that are rare within either stand. Such crosses should be reciprocal and replicated, however, and fruit and seed set should be compared with within-population crosses rather than assuming compatibility from genome-wide *F_ST_* alone.

Gene .2019 is a credible G-type lectin S-receptor-like kinase candidate because it is intact and contains the expected B-lectin, S-locus glycoprotein, PAN, transmembrane and kinase domains. The differentiated P736T substitution lies in the kinase domain and changes the local physicochemical environment without disrupting the predicted fold (**Figure 6**). This is an interesting functional candidate, not evidence that the substitution determines compatibility. Only 243 CDS positions met population-level coverage thresholds, principally within part of the kinase region, while the extracellular recognition domains were poorly recovered. Moreover, the male determinant remains unidentified, and the molecular architecture of SI in *Passiflora* has not been experimentally established (Suassuna et al., 2003; Madureira et al., 2014; Costa et al., 2021). Population differentiation at one receptor-like gene could reflect linked demographic history rather than balancing selection or S-haplotype function.

Resolving this system will require flower stage transcriptomics, phased long-read sequencing across the complete candidate interval and controlled diallel crosses among genotyped individuals. Transcriptomes from stigma/style and anthers or pollen should help identify co-expressed female and male recognition candidates, while long reads can recover highly divergent haplotypes that short-read mapping and GBS are likely to miss. Association between phased haplotypes and cross compatibility would then provide a direct functional test. Until those data are available, conservation crossing should maximize representation of both genomic populations and avoid selecting parents solely by the P736T genotype.

An integrated conservation strategy should combine in situ habitat protection with genetically informed ex situ propagation. Individuals from both populations, individual 28 and other admixed or genetically distinctive plants should be maintained as separate documented accessions. Reciprocal crosses between and within populations should be used to produce fertile fruits while retaining multiple parents, rather than creating a narrow set of highly related offspring. Because cultivated plants appear sensitive to heat and desiccation and may perform poorly in pots over long periods, propagation protocols should test cool, humid conditions and substrates that permit rhizome extension. Seed banking is especially valuable if dormancy and storage behaviour can be characterized, and in vitro culture or encapsulation may provide a complementary safeguard (Meng et al., 2021). Any translocation or reinforcement should retain provenance records and be preceded by crossing and common garden tests, because the observed genetic structure may include locally adapted variation as well as drift.

## CONCLUSION

Asian *Passiflora* diversification in supersection *Disemma* reflects the interaction of rapid lineage divergence, ILS, introgression and unusually dynamic plastome evolution. The *P. siamica*-*P. perakensis*-*P. cochinchinensis* lineage was stable across genomic compartments, whereas relationships among *P. xishuangbannaensis*, *P. menghaiensis*, *P. altebilobata* and *P. sumatrana* were strongly discordant and were better explained by a network than by a single fully bifurcating tree. The close relationship of *P. menghaiensis* to *P. napalensis* and its possible early position in alternative network backbones identify it as important for understanding the Asian radiation, but not as a demonstrated direct ancestor. Broader sampling across the Hua Line is required to distinguish ancient dispersal, organellar capture and reticulation.

The plastomes provide a second record of this history. Shared gene losses characterize *Disemma*, while repeated inversions and IR shifts generated exceptional structural variation, including the extreme expansion in *P. sumatrana*. These features are phylogenetically informative but, because the plastome behaves as a single inherited unit and plastid transmission is variable in *Passiflora*, they must be interpreted together with nuclear evidence.

For *P. xishuangbannaensis*, the conservation message is immediate. Thirty-nine sampled plants represented distinct genotypes divided between two differentiated populations, yet both demographic analyses indicated decline and recent *N*_e_ was low. Population 2 retained less diversity, while admixed individuals and individual 28 may preserve important connectivity. Controlled reciprocal crossing between populations, broad parental representation, habitat protection, seed banking and improved ex situ protocols should therefore be prioritized. The gene2019 P736T variant is a promising SI candidate but is not yet a functional marker of compatibility; transcriptomics, phased long-read sequencing and replicated crossing experiments are needed before molecularly guided mate selection is possible. Integrating these approaches with expanded sampling of *P. kwangtungensis*, *P. leschenaultii* and *P. napalensis* offers the clearest path toward resolving both the evolutionary history and the conservation requirements of this threatened Asian lineage.

## CRediT authorship contribution statement

Landrein Sven: Conceptualization, Methodology, Software, Validation, Formal analysis, Investigation, Data curation, Writing - original draft, Writing - review & editing, Visualization, Project administration. Zhou Zi Yu: Formal analysis, Investigation, Writing. Shook Ling Low: Methodology, Software, Formal analysis, Writing - review & editing. Niu Hong Bin: Investigation, Resources. Song Shi Jie: Investigation, Resources, Data curation. Wu Fu Chuan: Investigation, Resources, Project administration, Funding acquisition. Shen Jian Yong: Investigation, Resources, Project administration, Funding acquisition. Dong Hui: Writing - review & editing. Jiang Qiu Yu: Investigation, Resources, Data curation.

## Notes

### Competing Interest Statement

The authors have declared no competing interest.

