## Supplementary files for "Reticulate Evolution and Plastome Restructuring Shape Asian *Passiflora* Diversification"

| | Species | code | Locality | Voucher | Data/ Illumina | NCBI/SRA number |
|---|---|---|---|---|---|---|
| 1* | *Adenia cardiophylla* (Mast.) Engl. | Sven89 | Xishuangbanna, Yin Shang Shan | HITBC | 4.2 Gb | XXXXXX |
| 2* | *Adenia cordifolia* (Blume) Engl. | Bor9 | Borneo, Sabah, Pinangah Forest reserve | KFBG | 33.7 Gb | XXXXXX |
| 3* | *Adenia penangiana* (Wall. ex G.Don) W.J.de Wilde | Sven35 | Xishuangbanna, Mengla, Bu Beng Cun | HITBC | 5.2 Gb | XXXXXX |
| 4* | *Passiflora altebilobata* Hemsl. | Sven32 | Xishuangbanna, Meng Xing Pao Zhu Qing | HITBC | 5.3 Gb | XXXXXX |
| 5* | *Passiflora biflora* Lam. | Sven50 | Xishuangbanna Tropical Botanical Garden | HITBC | 5.8 Gb | XXXXXX |
| 7 | *Passiflora cinnabarina* Lindl. | PAH98 | | | 1.8 Gb | ERR5419864 |
| 8* | *Passiflora cochinchinensis* Spreng. | Kfbg21 | Hong Kong, Kadoorie Farm and Botanic Garden | KFBG | 10.6 Gb | XXXXXX |
| 9* | *Passiflora menghaiensis* X.D.Ma, L.C.Yan & J.Y.Shen | Sven31 | Xishuangbanna, Menghai | HITBC | 5.3 Gb | XXXXXX |
| 10 | *Passiflora papilio H.L.Li* | Fei Fei W. | Guangxi, Guilin | | | PZ899793 |
| 11* | *Passiflora perakensis* Hallier f. | Bor10 | Borneo, Sabah, Tambunan | KFBG | 39.4 Gb | SUB16508600 |
| 12* | *Passiflora siamica* Craib | Sven30 | Xishuangbanna, Ji Nuo Shan | HITBC | 5.3 Gb | XXXXXX |
| 13* | *Passiflora sumatrana* Blume | Sven33 | Xishuangbanna Tropical Botanical Garden | HITBC | 5.4 Gb | XXXXXX |
| 14 | *Passiflora xishuangbannaensis* Krosnick | | Xishuangbanna Tropical Botanical Garden | HITBC | 6.1 Gb | SRR13805787 |
| 15 | *Adenia mannii* (Mast.) Engl. | | NCBI assembled seq. | | N/A | MK651116 |
| 16 | *Passiflora actinia* Hook. | | NCBI assembled seq. | | N/A | MF807934 |
| 17 | *Passiflora affinis* Engelm. | | NCBI assembled seq. | | N/A | MK694930 |
| 18 | *Passiflora alata* Curtis | | NCBI assembled seq. | | N/A | MT525869 |
| 19 | *Passiflora arbelaezii* L.Uribe | | NCBI assembled seq. | | N/A | NC043819 |
| 20 | *Passiflora auriculata* Kunth | | NCBI assembled seq. | | N/A | MF807936 |
| 21 | *Passiflora biflora* Lam. | | NCBI assembled seq. | | N/A | MF807937 |
| 22 | *Passiflora candollei* Triana & Planch. | | NCBI assembled seq. | | N/A | MT525870 |
| 23 | *Passiflora capsularis* L. | | NCBI assembled seq. | | N/A | MT525883 |
| 24 | *Passiflora cerradensis* Sacco | | NCBI assembled seq. | | N/A | MT525871 |
| 25 | *Passiflora cincinnata* Mast. | | NCBI assembled seq. | | N/A | KY820583 |
| 25 | *Passiflora cirrhiflora* Juss. | | NCBI assembled seq. | | N/A | MN545921 |
| 27 | *Passiflora contracta* Vitta | | NCBI assembled seq. | | N/A | MK694925 |
| 28 | *Passiflora contracta* Vitta | | NCBI assembled seq. | | N/A | NC043818 |
| 29 | *Passiflora costaricensis* Killip | | NCBI assembled seq. | | N/A | MT473979 |

| | | | | |
|---|---|---|---|---|
| 30 | *Passiflora cristalina* Vanderpl. & Zappi | NCBI assembled seq. | N/A | MT525872 |
| 31 | *Passiflora deidamioides* Harms | NCBI assembled seq. | N/A | MT525873 |
| 32 | *Passiflora edmundoi* Sacco | NCBI assembled seq. | N/A | MT525874 |
| 33 | *Passiflora edulis* Sims | NCBI assembled seq. | N/A | KX290855 |
| 34 | *Passiflora edulis* Sims | NCBI assembled seq. | N/A | MT884000 |
| 35 | *Passiflora elegans* Mast. | NCBI assembled seq. | N/A | MN062356 |
| 36 | *Passiflora foetida* L. | NCBI assembled seq. | N/A | MK694932 |
| 37 | *Passiflora jatunsachensis* Schwerdtf. | NCBI assembled seq. | N/A | MK694920 |
| 38 | *Passiflora haematostigma* Mast. | NCBI assembled seq. | N/A | MT525875 |
| 39 | *Passiflora incarnata* L. | NCBI assembled seq. | N/A | MN062357 |
| 40 | *Passiflora laurifolia* L. | NCBI assembled seq. | N/A | MF807939 |
| 41 | *Passiflora ligularis* Juss. | NCBI assembled seq. | N/A | MF807940 |
| 42 | *Passiflora loefgrenii* Vitta | NCBI assembled seq. | N/A | MT525876 |
| 43 | *Passiflora lutea* L. | NCBI assembled seq. | N/A | MK694922 |
| 44 | *Passiflora malacophylla* Mast. | NCBI assembled seq. | N/A | MN062358 |
| 45 | *Passiflora maliformis* L. | NCBI assembled seq. | N/A | MN062359 |
| 46 | *Passiflora menispermifolia* Kunth | NCBI assembled seq. | N/A | MK694933 |
| 47 | *Passiflora microstipula* L.E.Gilbert & J.M.MacDougal | NCBI assembled seq. | N/A | MK694934 |
| 48 | *Passiflora miniata* Vanderpl. | NCBI assembled seq. | N/A | MT213977 |
| 49 | *Passiflora miniata* Vanderpl. | NCBI assembled seq. | N/A | MT525877 |
| 50 | *Passiflora misera* Kunth | NCBI assembled seq. | N/A | MK694928 |
| 51 | *Passiflora mucronata* Lam. | NCBI assembled seq. | N/A | MN062360 |
| 52 | *Passiflora nitida* Kunth | NCBI assembled seq. | N/A | MF807941 |
| 53 | *Passiflora obovata* Killip | NCBI assembled seq. | N/A | MK694931 |
| 54 | *Passiflora oerstedii* Mast. | NCBI assembled seq. | N/A | MF807942 |
| 55 | *Passiflora pittieri* Mast. | NCBI assembled seq. | N/A | MF807943 |
| 56 | *Passiflora quadrangularis* L. | NCBI assembled seq. | N/A | MF807944 |
| 57 | *Passiflora recurva* Mast. | NCBI assembled seq. | N/A | MT525879 |
| 58 | *Passiflora retipetala* Mast. | NCBI assembled seq. | N/A | MF807945 |
| 59 | *Passiflora rhamnifolia* Mast. | NCBI assembled seq. | N/A | MT525882 |
| 60 | *Passiflora rufa* Feuillet & J.M.MacDougal | NCBI assembled seq. | N/A | MK694924 |
| 61 | *Passiflora serratifolia* L. | NCBI assembled seq. | N/A | MF807948 |
| 62 | *Passiflora serratodigitata* L. | NCBI assembled seq. | N/A | MF807946 |
| 63 | *Passiflora serrulata* Jacq. | NCBI assembled seq. | N/A | MT677873 |
| 64 | *Passiflora suberosa* L. | NCBI assembled seq. | N/A | MT525868 |
| 65 | *Passiflora tenuiloba* Engelm. | NCBI assembled seq. | N/A | MK694923 |

| | | | | |
|---|---|---|---|---|
| 66 | *Passiflora tetrandra* Banks ex DC. | NCBI assembled seq. | N/A | MK694927 |
| 67 | *Passiflora tripartita var. mollissima* (Kunth) Holm-Niels. & P.M.Jørg. | NCBI assembled seq. | N/A | OQ910395 |
| 68 | *Passiflora vespertilio* L. | NCBI assembled seq. | N/A | MT525880 |
| 69 | *Passiflora vitifolia* Kunth | NCBI assembled seq. | N/A | MF807947 |
| 70 | *Passiflora watsoniana* Mast. | NCBI assembled seq. | N/A | MT525881 |

**Supplementary Table S1. List of samples used for plastid and phylogenomic analyses in this study.** Samples newly generated and sequenced for the present study are indicated in red and marked with an asterisk (*). All remaining samples represent previously published sequence data retrieved from NCBI/GenBank.

| sample | latitude | longitude | zone |
|---|---|---|---|
| s01 | 22.##### | 100.##### | Pop1 |
| s02 | 22.##### | 100.##### | Pop1 |
| s03 | 22.##### | 100.##### | Pop1 |
| s04 | 22.##### | 100.##### | Pop1 |
| s05 | 22.##### | 100.##### | Pop1 |
| s06 | 22.##### | 100.##### | Pop1 |
| s07 | 22.##### | 100.##### | Pop1 |
| s08 | 22.##### | 100.##### | Pop1 |
| s09 | 22.##### | 100.##### | Pop1 |
| s11 | 22.##### | 100.##### | Pop1 |
| s13 | 22.##### | 100.##### | Pop1 |
| s14 | 22.##### | 100.##### | Pop1 |
| s15 | 22.##### | 100.##### | Pop1 |
| s16 | 22.##### | 100.##### | Pop1 |
| s17 | 22.##### | 100.##### | Pop1 |
| s18 | 22.##### | 100.##### | Pop1 |
| s19 | 22.##### | 100.##### | Pop1 |
| s20 | 22.##### | 100.##### | Pop1 |
| s21 | 22.##### | 100.##### | Pop1 |
| s22 | 22.##### | 100.##### | Pop1 |
| s23 | 22.##### | 100.##### | Pop1 |
| s24 | 22.##### | 100.##### | Pop1 |
| s25 | 22.##### | 100.##### | Pop1 |
| s26 | 22.##### | 100.##### | Pop1 |
| s27 | 22.##### | 100.##### | Pop1 |
| s28 | 22.##### | 100.##### | |
| s29 | 22.##### | 100.##### | Pop2 |
| s30 | 22.##### | 100.##### | Pop2 |
| s31 | 22.##### | 100.##### | Pop2 |
| s32 | 22.##### | 100.##### | Pop2 |
| s33 | 22.##### | 100.##### | Pop2 |
| s34 | 22.##### | 100.##### | Pop2 |
| s35 | 22.##### | 100.##### | Pop2 |

| | | | |
|---|---|---|---|
| s36 | 22.##### | 100.##### | Pop2 |
| s38 | 22.##### | 100.##### | Pop2 |
| s40 | 22.##### | 100.##### | Pop2 |
| s41 | 22.##### | 100.##### | Pop2 |
| s42 | 22.##### | 100.##### | Pop2 |
| s43 | 22.##### | 100.##### | Pop2 |

**Supplementary Table S2. List of samples used for population genomic analyses of *Passiflora xishuangbannaensis*.** Precise GPS coordinates are withheld to protect the locations of this extremely rare species.

*atpA, atpB, atpE, atpF, atpH, atpI, ccsA, cemA, clpP, matK, ndhA, ndhB, ndhC, ndhD, ndhE, ndhF, ndhG, ndhH, ndhI, ndhJ, ndhK, petA, petB, petD, petG, petL, petN, psaA, psaB, psaC, psaI, psaJ, psbA, psbB, psbC, psbD, psbE, psbF, psbH, psbI, psbJ, psbK, psbL, psbM, psbN, psbT, psbZ, rbcL, rpl2, rpl14, rpl16, rpl23, rpl33, rpl36, rpoB, rpoC1, rpoC2, rps2, rps3, rps4, rps8, rps11, rps12, rps14, rps15, rps19, ycf3, ycf4.*

**Supplementary Table S3. Plastid loci used for phylogenetic analyses.** The 68 plastid protein-coding genes retained for phylogenetic reconstruction are listed above. Genes were extracted from the annotated plastomes, aligned individually, and concatenated for phylogenetic analysis.

***Passiflora altebilobata*** Hemsl. Krosnick 03: DQ458078/DQ458170/DQ458124/DQ458105/JX679732 ; ***P. aurantia*** G.Forst. AY632704/----/----/----/----; ***P. cinnabarina*** Lindl. : AY632706/----/----/----/----; ***P. cochinchinensis* var. *glaberrima*** (Gagnep.) G. Cusset : DQ284536/----/----/ ----/----; ***P. cupiformis*** Mast. Krosnick 253: AY632708/DQ458176/DQ458132/AY632733/JX679763; ***P. eberhardtii*** Gagnep. Krosnick 16 & 292: DQ458073/DQ458177/DQ458133/JX470877/JX679767; ***P. herbertiana*** Ker Gawl. AY632711/----/----/----/----; ***P. hollrungii*** K.Schum.: DQ45081/----/----/----/----; ***P. henryi*** Hemsl. Krosnick 08**:** AY632710/ DQ458180/ DQ458128/ AY632735/JX679777; **P. jianfengensis** S.M.Hwang & Q.Huang Krosnick 293: DQ458077/DQ458184/DQ458142/ DQ458113/----; ***P. jugorum*** W.W.Sm. Krosnick 15: AY632712/DQ458185/DQ458143/AY632737/JX679787 ; **P. kwangtungensis** Merr. : KF207865/----/----/----/----; ***P. leschenaultii*** DC. Krosnick 342 : DQ458079,KR350663/DQ458186/DQ458145/DQ458107/---- ; ***P. moluccana* var. *teysmanniana*** (Miq.) W.J.de Wilde Krosnick 326 & 327: DQ458080/DQ458189/DQ458151/ AY632739/JX679757; ***P. napalensis*** D.Don Krosnick 337: DQ458075/DQ458179/DQ458135/DQ458120/-----; **P. papilio** H.L.Li : DQ458074/----/ DQ458157/ DQ458119/----; ***P. perakensis*** Hallier f. Krosnick 314 : DQ087422/DQ458195/DQ458158/ DQ087431/JX679819 ; ***P. siamica*** Craib Krosnick 320 **:** DQ458215/ DQ458197/DQ458160/DQ458115/JX679838 ; ***P. sumatrana*** Blume J. Wen 5973 :DQ458072/DQ458201/DQ458165/DQ087434/ JX679856 ; ***P. tonkinensis*** W.J.de Wilde Krosnick 3248 : DQ087424/DQ45818/DQ458163/DQ087433/----; ***P. xishuangbannaensis*** Krosnick : DQ458071/DQ458203/DQ458167/DQ458109/----.

**Supplementary Table S4. GenBank accession numbers for the five molecular markers used in the comparison with previously published Asian *Passiflora* data.** For each taxon, accession numbers are presented in the following order: nuclear ribosomal ITS region (ITS1–5.8S–ITS2), cytosolic glutamine synthetase (*cytGS*), nuclear encoded chloroplast glutamine synthetase (*ncpGS*), *trnL-trnF*, and *ndhF*. Dashes indicate sequences that were unavailable for the corresponding taxon and marker.

Analysis locus \| P. organensis reference CDS/transcript ID

| | |
|---|---|
| gene_0001 | EVM%20prediction%20scaffold103_size347917.11.mRNA1 |
| gene_0005 | EVM%20prediction%20scaffold106_size336076.2.mRNA1 |
| gene_0009 | EVM%20prediction%20scaffold10_size6807042.140.mRNA1 |
| gene_0010 | EVM%20prediction%20scaffold10_size6807042.141.mRNA1 |
| gene_0012 | EVM%20prediction%20scaffold10_size6807042.270.mRNA1 |
| gene_0014 | EVM%20prediction%20scaffold10_size6807042.477.mRNA1 |
| gene_0016 | EVM%20prediction%20scaffold10_size6807042.552.mRNA1 |

| | |
|---|---|
| gene_0018 | EVM%20prediction%20scaffold10_size6807042.75.mRNA1 |
| gene_0019 | EVM%20prediction%20scaffold11_size6505685.110.mRNA1 |
| gene_0020 | EVM%20prediction%20scaffold11_size6505685.123.mRNA1 |
| gene_0021 | EVM%20prediction%20scaffold11_size6505685.136.mRNA1 |
| gene_0022 | EVM%20prediction%20scaffold11_size6505685.182.mRNA1 |
| gene_0023 | EVM%20prediction%20scaffold11_size6505685.188.mRNA1 |
| gene_0024 | EVM%20prediction%20scaffold11_size6505685.2.mRNA1 |
| gene_0025 | EVM%20prediction%20scaffold11_size6505685.238.mRNA1 |
| gene_0029 | EVM%20prediction%20scaffold11_size6505685.333.mRNA1 |
| gene_0030 | EVM%20prediction%20scaffold11_size6505685.340.mRNA1 |
| gene_0032 | EVM%20prediction%20scaffold11_size6505685.412.mRNA1 |
| gene_0033 | EVM%20prediction%20scaffold11_size6505685.433.mRNA1 |
| gene_0035 | EVM%20prediction%20scaffold11_size6505685.51.mRNA1 |
| gene_0036 | EVM%20prediction%20scaffold11_size6505685.518.mRNA1 |
| gene_0037 | EVM%20prediction%20scaffold11_size6505685.536.mRNA1 |
| gene_0041 | EVM%20prediction%20scaffold12_size3927828.210.mRNA1 |
| gene_0042 | EVM%20prediction%20scaffold12_size3927828.240.mRNA1 |
| gene_0044 | EVM%20prediction%20scaffold12_size3927828.57.mRNA1 |
| gene_0045 | EVM%20prediction%20scaffold12_size3927828.84.mRNA1 |
| gene_0047 | EVM%20prediction%20scaffold131_size248657.16.mRNA1 |
| gene_0049 | EVM%20prediction%20scaffold13_size3393329.95.mRNA1 |
| gene_0051 | EVM%20prediction%20scaffold14_size3361460.172.mRNA1 |
| gene_0052 | EVM%20prediction%20scaffold14_size3361460.27.mRNA1 |
| gene_0053 | EVM%20prediction%20scaffold14_size3361460.341.mRNA1 |
| gene_0055 | EVM%20prediction%20scaffold14_size3361460.41.mRNA1 |
| gene_0056 | EVM%20prediction%20scaffold14_size3361460.42.mRNA1 |
| gene_0057 | EVM%20prediction%20scaffold14_size3361460.98.mRNA1 |
| gene_0061 | EVM%20prediction%20scaffold162_size137447.4.mRNA1 |
| gene_0064 | EVM%20prediction%20scaffold16_size2738918.131.mRNA1 |
| gene_0067 | EVM%20prediction%20scaffold16_size2738918.30.mRNA2 |
| gene_0074 | EVM%20prediction%20scaffold176_size116420.5.mRNA1 |
| gene_0075 | EVM%20prediction%20scaffold176_size116420.8.mRNA1 |
| gene_0077 | EVM%20prediction%20scaffold18_size2475805.151.mRNA1 |
| gene_0078 | EVM%20prediction%20scaffold19_size2367737.35.mRNA1 |
| gene_0079 | EVM%20prediction%20scaffold1_size35943597.1035.mRNA1 |
| gene_0080 | EVM%20prediction%20scaffold1_size35943597.1052.mRNA1 |
| gene_0081 | EVM%20prediction%20scaffold1_size35943597.113.mRNA1 |
| gene_0082 | EVM%20prediction%20scaffold1_size35943597.1182.mRNA1 |
| gene_0088 | EVM%20prediction%20scaffold1_size35943597.148.mRNA1 |
| gene_0089 | EVM%20prediction%20scaffold1_size35943597.1533.mRNA1 |
| gene_0090 | EVM%20prediction%20scaffold1_size35943597.1582.mRNA1 |
| gene_0092 | EVM%20prediction%20scaffold1_size35943597.1659.mRNA1 |
| gene_0094 | EVM%20prediction%20scaffold1_size35943597.1708.mRNA1 |
| gene_0095 | EVM%20prediction%20scaffold1_size35943597.178.mRNA1 |
| gene_0096 | EVM%20prediction%20scaffold1_size35943597.1828.mRNA1 |
| gene_0097 | EVM%20prediction%20scaffold1_size35943597.183.mRNA1 |
| gene_0099 | EVM%20prediction%20scaffold1_size35943597.1869.mRNA1 |
| gene_0100 | EVM%20prediction%20scaffold1_size35943597.189.mRNA1 |
| gene_0101 | EVM%20prediction%20scaffold1_size35943597.1899.mRNA1 |
| gene_0103 | EVM%20prediction%20scaffold1_size35943597.1944.mRNA1 |
| gene_0105 | EVM%20prediction%20scaffold1_size35943597.2036.mRNA1 |
| gene_0106 | EVM%20prediction%20scaffold1_size35943597.2037.mRNA1 |
| gene_0107 | EVM%20prediction%20scaffold1_size35943597.2107.mRNA1 |
| gene_0108 | EVM%20prediction%20scaffold1_size35943597.2135.mRNA1 |
| gene_0109 | EVM%20prediction%20scaffold1_size35943597.2209.mRNA1 |
| gene_0110 | EVM%20prediction%20scaffold1_size35943597.222.mRNA1 |
| gene_0111 | EVM%20prediction%20scaffold1_size35943597.2229.mRNA1 |
| gene_0112 | EVM%20prediction%20scaffold1_size35943597.2250.mRNA1 |
| gene_0114 | EVM%20prediction%20scaffold1_size35943597.2281.mRNA1 |
| gene_0115 | EVM%20prediction%20scaffold1_size35943597.233.mRNA3 |
| gene_0116 | EVM%20prediction%20scaffold1_size35943597.2352.mRNA1 |
| gene_0122 | EVM%20prediction%20scaffold1_size35943597.245.mRNA1 |
| gene_0123 | EVM%20prediction%20scaffold1_size35943597.2596.mRNA1 |
| gene_0124 | EVM%20prediction%20scaffold1_size35943597.2601.mRNA1 |
| gene_0128 | EVM%20prediction%20scaffold1_size35943597.2636.mRNA1 |
| gene_0130 | EVM%20prediction%20scaffold1_size35943597.2693.mRNA1 |
| gene_0132 | EVM%20prediction%20scaffold1_size35943597.2777.mRNA1 |
| gene_0133 | EVM%20prediction%20scaffold1_size35943597.280.mRNA1 |
| gene_0134 | EVM%20prediction%20scaffold1_size35943597.2884.mRNA1 |
| gene_0136 | EVM%20prediction%20scaffold1_size35943597.2949.mRNA1 |

| | |
|---|---|
| gene_0137 | EVM%20prediction%20scaffold1_size35943597.2963.mRNA2 |
| gene_0138 | EVM%20prediction%20scaffold1_size35943597.2964.mRNA1 |
| gene_0140 | EVM%20prediction%20scaffold1_size35943597.3064.mRNA1 |
| gene_0141 | EVM%20prediction%20scaffold1_size35943597.3168.mRNA1 |
| gene_0142 | EVM%20prediction%20scaffold1_size35943597.3174.mRNA1 |
| gene_0144 | EVM%20prediction%20scaffold1_size35943597.3199.mRNA1 |
| gene_0147 | EVM%20prediction%20scaffold1_size35943597.3438.mRNA1 |
| gene_0149 | EVM%20prediction%20scaffold1_size35943597.3511.mRNA1 |
| gene_0150 | EVM%20prediction%20scaffold1_size35943597.3568.mRNA1 |
| gene_0151 | EVM%20prediction%20scaffold1_size35943597.3599.mRNA1 |
| gene_0152 | EVM%20prediction%20scaffold1_size35943597.3600.mRNA1 |
| gene_0153 | EVM%20prediction%20scaffold1_size35943597.3735.mRNA1 |
| gene_0154 | EVM%20prediction%20scaffold1_size35943597.38.mRNA1 |
| gene_0156 | EVM%20prediction%20scaffold1_size35943597.385.mRNA1 |
| gene_0161 | EVM%20prediction%20scaffold1_size35943597.4183.mRNA1 |
| gene_0163 | EVM%20prediction%20scaffold1_size35943597.423.mRNA1 |
| gene_0165 | EVM%20prediction%20scaffold1_size35943597.4242.mRNA1 |
| gene_0169 | EVM%20prediction%20scaffold1_size35943597.4603.mRNA1 |
| gene_0171 | EVM%20prediction%20scaffold1_size35943597.4837.mRNA1 |
| gene_0173 | EVM%20prediction%20scaffold1_size35943597.4898.mRNA1 |
| gene_0175 | EVM%20prediction%20scaffold1_size35943597.4909.mRNA1 |
| gene_0180 | EVM%20prediction%20scaffold1_size35943597.650.mRNA1 |
| gene_0181 | EVM%20prediction%20scaffold1_size35943597.666.mRNA1 |
| gene_0182 | EVM%20prediction%20scaffold1_size35943597.750.mRNA1 |
| gene_0183 | EVM%20prediction%20scaffold1_size35943597.782.mRNA1 |
| gene_0184 | EVM%20prediction%20scaffold1_size35943597.797.mRNA1 |
| gene_0185 | EVM%20prediction%20scaffold1_size35943597.825.mRNA1 |
| gene_0186 | EVM%20prediction%20scaffold1_size35943597.887.mRNA1 |
| gene_0187 | EVM%20prediction%20scaffold1_size35943597.922.mRNA1 |
| gene_0188 | EVM%20prediction%20scaffold1_size35943597.927.mRNA1 |
| gene_0190 | EVM%20prediction%20scaffold208_size76882.1.mRNA1 |
| gene_0195 | EVM%20prediction%20scaffold22_size2164848.47.mRNA1 |
| gene_0199 | EVM%20prediction%20scaffold24_size1974514.8.mRNA1 |
| gene_0204 | EVM%20prediction%20scaffold27_size1605964.5.mRNA1 |
| gene_0205 | EVM%20prediction%20scaffold27_size1605964.74.mRNA1 |
| gene_0206 | EVM%20prediction%20scaffold29_size1428910.118.mRNA1 |
| gene_0207 | EVM%20prediction%20scaffold29_size1428910.55.mRNA1 |
| gene_0208 | EVM%20prediction%20scaffold29_size1428910.57.mRNA1 |
| gene_0211 | EVM%20prediction%20scaffold29_size1428910.94.mRNA1 |
| gene_0212 | EVM%20prediction%20scaffold29_size1428910.97.mRNA1 |
| gene_0213 | EVM%20prediction%20scaffold2_size22687060.1034.mRNA1 |
| gene_0214 | EVM%20prediction%20scaffold2_size22687060.1036.mRNA1 |
| gene_0216 | EVM%20prediction%20scaffold2_size22687060.1120.mRNA1 |
| gene_0217 | EVM%20prediction%20scaffold2_size22687060.1172.mRNA1 |
| gene_0218 | EVM%20prediction%20scaffold2_size22687060.1182.mRNA1 |
| gene_0219 | EVM%20prediction%20scaffold2_size22687060.1195.mRNA1 |
| gene_0220 | EVM%20prediction%20scaffold2_size22687060.1244.mRNA1 |
| gene_0221 | EVM%20prediction%20scaffold2_size22687060.1288.mRNA1 |
| gene_0222 | EVM%20prediction%20scaffold2_size22687060.1290.mRNA1 |
| gene_0223 | EVM%20prediction%20scaffold2_size22687060.1305.mRNA1 |
| gene_0226 | EVM%20prediction%20scaffold2_size22687060.1514.mRNA1 |
| gene_0228 | EVM%20prediction%20scaffold2_size22687060.1580.mRNA1 |
| gene_0229 | EVM%20prediction%20scaffold2_size22687060.162.mRNA1 |
| gene_0231 | EVM%20prediction%20scaffold2_size22687060.1862.mRNA1 |
| gene_0233 | EVM%20prediction%20scaffold2_size22687060.1936.mRNA1 |
| gene_0234 | EVM%20prediction%20scaffold2_size22687060.1982.mRNA1 |
| gene_0235 | EVM%20prediction%20scaffold2_size22687060.199.mRNA1 |
| gene_0238 | EVM%20prediction%20scaffold2_size22687060.2088.mRNA1 |
| gene_0240 | EVM%20prediction%20scaffold2_size22687060.2130.mRNA1 |
| gene_0241 | EVM%20prediction%20scaffold2_size22687060.2137.mRNA1 |
| gene_0242 | EVM%20prediction%20scaffold2_size22687060.2157.mRNA1 |
| gene_0244 | EVM%20prediction%20scaffold2_size22687060.2190.mRNA1 |
| gene_0247 | EVM%20prediction%20scaffold2_size22687060.2280.mRNA1 |
| gene_0249 | EVM%20prediction%20scaffold2_size22687060.2319.mRNA1 |
| gene_0250 | EVM%20prediction%20scaffold2_size22687060.232.mRNA1 |
| gene_0253 | EVM%20prediction%20scaffold2_size22687060.2423.mRNA1 |
| gene_0254 | EVM%20prediction%20scaffold2_size22687060.2488.mRNA1 |
| gene_0255 | EVM%20prediction%20scaffold2_size22687060.2572.mRNA1 |
| gene_0257 | EVM%20prediction%20scaffold2_size22687060.2619.mRNA1 |
| gene_0265 | EVM%20prediction%20scaffold2_size22687060.279.mRNA1 |

| | |
|---|---|
| gene_0266 | EVM%20prediction%20scaffold2_size22687060.280.mRNA1 |
| gene_0268 | EVM%20prediction%20scaffold2_size22687060.2815.mRNA1 |
| gene_0269 | EVM%20prediction%20scaffold2_size22687060.2881.mRNA1 |
| gene_0270 | EVM%20prediction%20scaffold2_size22687060.2911.mRNA1 |
| gene_0275 | EVM%20prediction%20scaffold2_size22687060.2975.mRNA1 |
| gene_0280 | EVM%20prediction%20scaffold2_size22687060.321.mRNA1 |
| gene_0281 | EVM%20prediction%20scaffold2_size22687060.323.mRNA1 |
| gene_0282 | EVM%20prediction%20scaffold2_size22687060.3237.mRNA1 |
| gene_0286 | EVM%20prediction%20scaffold2_size22687060.331.mRNA1 |
| gene_0289 | EVM%20prediction%20scaffold2_size22687060.3568.mRNA1 |
| gene_0290 | EVM%20prediction%20scaffold2_size22687060.3679.mRNA1 |
| gene_0291 | EVM%20prediction%20scaffold2_size22687060.3680.mRNA1 |
| gene_0292 | EVM%20prediction%20scaffold2_size22687060.3717.mRNA1 |
| gene_0293 | EVM%20prediction%20scaffold2_size22687060.3728.mRNA1 |
| gene_0294 | EVM%20prediction%20scaffold2_size22687060.3742.mRNA4 |
| gene_0295 | EVM%20prediction%20scaffold2_size22687060.3745.mRNA1 |
| gene_0297 | EVM%20prediction%20scaffold2_size22687060.381.mRNA1 |
| gene_0299 | EVM%20prediction%20scaffold2_size22687060.3850.mRNA2 |
| gene_0300 | EVM%20prediction%20scaffold2_size22687060.3852.mRNA1 |
| gene_0303 | EVM%20prediction%20scaffold2_size22687060.439.mRNA1 |
| gene_0305 | EVM%20prediction%20scaffold2_size22687060.831.mRNA1 |
| gene_0307 | EVM%20prediction%20scaffold2_size22687060.887.mRNA1 |
| gene_0308 | EVM%20prediction%20scaffold2_size22687060.899.mRNA1 |
| gene_0309 | EVM%20prediction%20scaffold2_size22687060.914.mRNA1 |
| gene_0310 | EVM%20prediction%20scaffold30_size1410943.43.mRNA1 |
| gene_0311 | EVM%20prediction%20scaffold31_size1301498.148.mRNA1 |
| gene_0312 | EVM%20prediction%20scaffold31_size1301498.174.mRNA1 |
| gene_0313 | EVM%20prediction%20scaffold31_size1301498.177.mRNA2 |
| gene_0314 | EVM%20prediction%20scaffold31_size1301498.178.mRNA1 |
| gene_0315 | EVM%20prediction%20scaffold31_size1301498.185.mRNA1 |
| gene_0316 | EVM%20prediction%20scaffold31_size1301498.36.mRNA1 |
| gene_0319 | EVM%20prediction%20scaffold31_size1301498.88.mRNA1 |
| gene_0321 | EVM%20prediction%20scaffold35_size1092366.20.mRNA1 |
| gene_0323 | EVM%20prediction%20scaffold3_size12432716.1008.mRNA1 |
| gene_0324 | EVM%20prediction%20scaffold3_size12432716.1037.mRNA2 |
| gene_0325 | EVM%20prediction%20scaffold3_size12432716.1054.mRNA1 |
| gene_0328 | EVM%20prediction%20scaffold3_size12432716.116.mRNA1 |
| gene_0330 | EVM%20prediction%20scaffold3_size12432716.1231.mRNA1 |
| gene_0331 | EVM%20prediction%20scaffold3_size12432716.1239.mRNA1 |
| gene_0332 | EVM%20prediction%20scaffold3_size12432716.1246.mRNA1 |
| gene_0333 | EVM%20prediction%20scaffold3_size12432716.1259.mRNA1 |
| gene_0335 | EVM%20prediction%20scaffold3_size12432716.1359.mRNA1 |
| gene_0339 | EVM%20prediction%20scaffold3_size12432716.1491.mRNA1 |
| gene_0340 | EVM%20prediction%20scaffold3_size12432716.1516.mRNA1 |
| gene_0342 | EVM%20prediction%20scaffold3_size12432716.165.mRNA1 |
| gene_0344 | EVM%20prediction%20scaffold3_size12432716.1736.mRNA1 |
| gene_0345 | EVM%20prediction%20scaffold3_size12432716.1754.mRNA1 |
| gene_0349 | EVM%20prediction%20scaffold3_size12432716.1933.mRNA1 |
| gene_0350 | EVM%20prediction%20scaffold3_size12432716.1939.mRNA4 |
| gene_0352 | EVM%20prediction%20scaffold3_size12432716.1964.mRNA1 |
| gene_0355 | EVM%20prediction%20scaffold3_size12432716.2053.mRNA1 |
| gene_0356 | EVM%20prediction%20scaffold3_size12432716.218.mRNA1 |
| gene_0357 | EVM%20prediction%20scaffold3_size12432716.256.mRNA1 |
| gene_0358 | EVM%20prediction%20scaffold3_size12432716.277.mRNA1 |
| gene_0360 | EVM%20prediction%20scaffold3_size12432716.374.mRNA1 |
| gene_0365 | EVM%20prediction%20scaffold3_size12432716.44.mRNA1 |
| gene_0366 | EVM%20prediction%20scaffold3_size12432716.45.mRNA1 |
| gene_0370 | EVM%20prediction%20scaffold3_size12432716.583.mRNA1 |
| gene_0372 | EVM%20prediction%20scaffold3_size12432716.615.mRNA3 |
| gene_0373 | EVM%20prediction%20scaffold3_size12432716.766.mRNA1 |
| gene_0377 | EVM%20prediction%20scaffold3_size12432716.958.mRNA1 |
| gene_0378 | EVM%20prediction%20scaffold3_size12432716.981.mRNA1 |
| gene_0379 | EVM%20prediction%20scaffold40_size983166.17.mRNA1 |
| gene_0380 | EVM%20prediction%20scaffold40_size983166.9.mRNA1 |
| gene_0381 | EVM%20prediction%20scaffold41_size972950.37.mRNA1 |
| gene_0382 | EVM%20prediction%20scaffold41_size972950.45.mRNA1 |
| gene_0384 | EVM%20prediction%20scaffold44_size936324.20.mRNA1 |
| gene_0389 | EVM%20prediction%20scaffold45_size927446.79.mRNA1 |
| gene_0390 | EVM%20prediction%20scaffold48_size872349.18.mRNA1 |
| gene_0391 | EVM%20prediction%20scaffold48_size872349.25.mRNA1 |

| | |
|---|---|
| gene_0394 | EVM%20prediction%20scaffold49_size841913.7.mRNA1 |
| gene_0396 | EVM%20prediction%20scaffold4_size11498481.1094.mRNA1 |
| gene_0397 | EVM%20prediction%20scaffold4_size11498481.1101.mRNA1 |
| gene_0398 | EVM%20prediction%20scaffold4_size11498481.1140.mRNA1 |
| gene_0399 | EVM%20prediction%20scaffold4_size11498481.1175.mRNA1 |
| gene_0401 | EVM%20prediction%20scaffold4_size11498481.1239.mRNA1 |
| gene_0403 | EVM%20prediction%20scaffold4_size11498481.1343.mRNA1 |
| gene_0407 | EVM%20prediction%20scaffold4_size11498481.1411.mRNA1 |
| gene_0408 | EVM%20prediction%20scaffold4_size11498481.1443.mRNA1 |
| gene_0410 | EVM%20prediction%20scaffold4_size11498481.214.mRNA1 |
| gene_0412 | EVM%20prediction%20scaffold4_size11498481.304.mRNA1 |
| gene_0415 | EVM%20prediction%20scaffold4_size11498481.378.mRNA1 |
| gene_0416 | EVM%20prediction%20scaffold4_size11498481.379.mRNA1 |
| gene_0418 | EVM%20prediction%20scaffold4_size11498481.471.mRNA1 |
| gene_0419 | EVM%20prediction%20scaffold4_size11498481.483.mRNA1 |
| gene_0422 | EVM%20prediction%20scaffold4_size11498481.531.mRNA1 |
| gene_0424 | EVM%20prediction%20scaffold4_size11498481.576.mRNA1 |
| gene_0426 | EVM%20prediction%20scaffold4_size11498481.594.mRNA1 |
| gene_0427 | EVM%20prediction%20scaffold4_size11498481.623.mRNA1 |
| gene_0428 | EVM%20prediction%20scaffold4_size11498481.680.mRNA1 |
| gene_0429 | EVM%20prediction%20scaffold4_size11498481.682.mRNA1 |
| gene_0430 | EVM%20prediction%20scaffold4_size11498481.69.mRNA1 |
| gene_0432 | EVM%20prediction%20scaffold4_size11498481.763.mRNA1 |
| gene_0433 | EVM%20prediction%20scaffold4_size11498481.821.mRNA1 |
| gene_0434 | EVM%20prediction%20scaffold4_size11498481.833.mRNA1 |
| gene_0444 | EVM%20prediction%20scaffold4_size11498481.982.mRNA1 |
| gene_0448 | EVM%20prediction%20scaffold58_size703183.8.mRNA1 |
| gene_0449 | EVM%20prediction%20scaffold5_size10758477.1074.mRNA1 |
| gene_0450 | EVM%20prediction%20scaffold5_size10758477.1080.mRNA1 |
| gene_0452 | EVM%20prediction%20scaffold5_size10758477.109.mRNA1 |
| gene_0453 | EVM%20prediction%20scaffold5_size10758477.1119.mRNA1 |
| gene_0455 | EVM%20prediction%20scaffold5_size10758477.1170.mRNA1 |
| gene_0456 | EVM%20prediction%20scaffold5_size10758477.1179.mRNA1 |
| gene_0457 | EVM%20prediction%20scaffold5_size10758477.1188.mRNA1 |
| gene_0458 | EVM%20prediction%20scaffold5_size10758477.1221.mRNA1 |
| gene_0459 | EVM%20prediction%20scaffold5_size10758477.1230.mRNA1 |
| gene_0464 | EVM%20prediction%20scaffold5_size10758477.1393.mRNA1 |
| gene_0465 | EVM%20prediction%20scaffold5_size10758477.1408.mRNA1 |
| gene_0466 | EVM%20prediction%20scaffold5_size10758477.141.mRNA1 |
| gene_0467 | EVM%20prediction%20scaffold5_size10758477.1644.mRNA1 |
| gene_0471 | EVM%20prediction%20scaffold5_size10758477.292.mRNA1 |
| gene_0473 | EVM%20prediction%20scaffold5_size10758477.294.mRNA1 |
| gene_0477 | EVM%20prediction%20scaffold5_size10758477.45.mRNA1 |
| gene_0479 | EVM%20prediction%20scaffold5_size10758477.471.mRNA1 |
| gene_0480 | EVM%20prediction%20scaffold5_size10758477.482.mRNA1 |
| gene_0482 | EVM%20prediction%20scaffold5_size10758477.484.mRNA1 |
| gene_0484 | EVM%20prediction%20scaffold5_size10758477.552.mRNA1 |
| gene_0485 | EVM%20prediction%20scaffold5_size10758477.632.mRNA1 |
| gene_0486 | EVM%20prediction%20scaffold5_size10758477.699.mRNA1 |
| gene_0489 | EVM%20prediction%20scaffold5_size10758477.782.mRNA1 |
| gene_0493 | EVM%20prediction%20scaffold5_size10758477.800.mRNA1 |
| gene_0497 | EVM%20prediction%20scaffold5_size10758477.910.mRNA1 |
| gene_0499 | EVM%20prediction%20scaffold5_size10758477.979.mRNA1 |
| gene_0500 | EVM%20prediction%20scaffold65_size636131.25.mRNA1 |
| gene_0501 | EVM%20prediction%20scaffold65_size636131.81.mRNA1 |
| gene_0502 | EVM%20prediction%20scaffold66_size618479.25.mRNA1 |
| gene_0503 | EVM%20prediction%20scaffold67_size617749.11.mRNA1 |
| gene_0504 | EVM%20prediction%20scaffold67_size617749.32.mRNA1 |
| gene_0505 | EVM%20prediction%20scaffold67_size617749.34.mRNA1 |
| gene_0507 | EVM%20prediction%20scaffold6_size10383153.110.mRNA1 |
| gene_0509 | EVM%20prediction%20scaffold6_size10383153.220.mRNA1 |
| gene_0512 | EVM%20prediction%20scaffold6_size10383153.297.mRNA1 |
| gene_0514 | EVM%20prediction%20scaffold6_size10383153.332.mRNA1 |
| gene_0516 | EVM%20prediction%20scaffold6_size10383153.359.mRNA1 |
| gene_0518 | EVM%20prediction%20scaffold6_size10383153.498.mRNA1 |
| gene_0520 | EVM%20prediction%20scaffold6_size10383153.727.mRNA1 |
| gene_0521 | EVM%20prediction%20scaffold6_size10383153.83.mRNA1 |
| gene_0523 | EVM%20prediction%20scaffold6_size10383153.880.mRNA1 |
| gene_0525 | EVM%20prediction%20scaffold6_size10383153.907.mRNA1 |
| gene_0528 | EVM%20prediction%20scaffold79_size476836.6.mRNA1 |

| | |
|---|---|
| gene_0529 | EVM%20prediction%20scaffold7_size10217246.1052.mRNA1 |
| gene_0537 | EVM%20prediction%20scaffold7_size10217246.1264.mRNA1 |
| gene_0545 | EVM%20prediction%20scaffold7_size10217246.128.mRNA1 |
| gene_0546 | EVM%20prediction%20scaffold7_size10217246.1360.mRNA1 |
| gene_0547 | EVM%20prediction%20scaffold7_size10217246.1367.mRNA1 |
| gene_0548 | EVM%20prediction%20scaffold7_size10217246.140.mRNA1 |
| gene_0550 | EVM%20prediction%20scaffold7_size10217246.1417.mRNA1 |
| gene_0554 | EVM%20prediction%20scaffold7_size10217246.1528.mRNA1 |
| gene_0555 | EVM%20prediction%20scaffold7_size10217246.155.mRNA1 |
| gene_0556 | EVM%20prediction%20scaffold7_size10217246.1577.mRNA1 |
| gene_0557 | EVM%20prediction%20scaffold7_size10217246.1600.mRNA1 |
| gene_0558 | EVM%20prediction%20scaffold7_size10217246.1639.mRNA1 |
| gene_0559 | EVM%20prediction%20scaffold7_size10217246.1645.mRNA1 |
| gene_0560 | EVM%20prediction%20scaffold7_size10217246.1652.mRNA1 |
| gene_0562 | EVM%20prediction%20scaffold7_size10217246.1725.mRNA1 |
| gene_0563 | EVM%20prediction%20scaffold7_size10217246.174.mRNA3 |
| gene_0564 | EVM%20prediction%20scaffold7_size10217246.1752.mRNA1 |
| gene_0566 | EVM%20prediction%20scaffold7_size10217246.1941.mRNA1 |
| gene_0567 | EVM%20prediction%20scaffold7_size10217246.1991.mRNA1 |
| gene_0568 | EVM%20prediction%20scaffold7_size10217246.2016.mRNA1 |
| gene_0569 | EVM%20prediction%20scaffold7_size10217246.2017.mRNA1 |
| gene_0570 | EVM%20prediction%20scaffold7_size10217246.2040.mRNA1 |
| gene_0571 | EVM%20prediction%20scaffold7_size10217246.2072.mRNA1 |
| gene_0572 | EVM%20prediction%20scaffold7_size10217246.230.mRNA1 |
| gene_0573 | EVM%20prediction%20scaffold7_size10217246.243.mRNA5 |
| gene_0574 | EVM%20prediction%20scaffold7_size10217246.258.mRNA1 |
| gene_0577 | EVM%20prediction%20scaffold7_size10217246.340.mRNA1 |
| gene_0578 | EVM%20prediction%20scaffold7_size10217246.440.mRNA1 |
| gene_0581 | EVM%20prediction%20scaffold7_size10217246.575.mRNA1 |
| gene_0583 | EVM%20prediction%20scaffold7_size10217246.650.mRNA1 |
| gene_0584 | EVM%20prediction%20scaffold7_size10217246.657.mRNA1 |
| gene_0585 | EVM%20prediction%20scaffold7_size10217246.667.mRNA1 |
| gene_0590 | EVM%20prediction%20scaffold7_size10217246.902.mRNA1 |
| gene_0591 | EVM%20prediction%20scaffold7_size10217246.923.mRNA1 |
| gene_0594 | EVM%20prediction%20scaffold8_size8616334.1024.mRNA1 |
| gene_0596 | EVM%20prediction%20scaffold8_size8616334.1049.mRNA1 |
| gene_0597 | EVM%20prediction%20scaffold8_size8616334.1072.mRNA1 |
| gene_0598 | EVM%20prediction%20scaffold8_size8616334.1089.mRNA1 |
| gene_0599 | EVM%20prediction%20scaffold8_size8616334.117.mRNA1 |
| gene_0603 | EVM%20prediction%20scaffold8_size8616334.144.mRNA1 |
| gene_0604 | EVM%20prediction%20scaffold8_size8616334.203.mRNA1 |
| gene_0607 | EVM%20prediction%20scaffold8_size8616334.29.mRNA1 |
| gene_0608 | EVM%20prediction%20scaffold8_size8616334.309.mRNA1 |
| gene_0610 | EVM%20prediction%20scaffold8_size8616334.382.mRNA1 |
| gene_0612 | EVM%20prediction%20scaffold8_size8616334.426.mRNA1 |
| gene_0613 | EVM%20prediction%20scaffold8_size8616334.441.mRNA1 |
| gene_0614 | EVM%20prediction%20scaffold8_size8616334.465.mRNA1 |
| gene_0617 | EVM%20prediction%20scaffold8_size8616334.534.mRNA1 |
| gene_0618 | EVM%20prediction%20scaffold8_size8616334.606.mRNA1 |
| gene_0621 | EVM%20prediction%20scaffold8_size8616334.67.mRNA1 |
| gene_0623 | EVM%20prediction%20scaffold8_size8616334.730.mRNA1 |
| gene_0625 | EVM%20prediction%20scaffold8_size8616334.778.mRNA1 |
| gene_0626 | EVM%20prediction%20scaffold8_size8616334.798.mRNA1 |
| gene_0633 | EVM%20prediction%20scaffold8_size8616334.890.mRNA1 |
| gene_0634 | EVM%20prediction%20scaffold8_size8616334.894.mRNA1 |
| gene_0636 | EVM%20prediction%20scaffold8_size8616334.93.mRNA1 |
| gene_0640 | EVM%20prediction%20scaffold8_size8616334.999.mRNA1 |
| gene_0641 | EVM%20prediction%20scaffold95_size369776.4.mRNA1 |
| gene_0642 | EVM%20prediction%20scaffold9_size8259397.19.mRNA1 |
| gene_0643 | EVM%20prediction%20scaffold9_size8259397.234.mRNA1 |
| gene_0644 | EVM%20prediction%20scaffold9_size8259397.27.mRNA1 |
| gene_0647 | EVM%20prediction%20scaffold9_size8259397.369.mRNA1 |
| gene_0650 | EVM%20prediction%20scaffold9_size8259397.429.mRNA1 |
| gene_0651 | EVM%20prediction%20scaffold9_size8259397.435.mRNA1 |
| gene_0652 | EVM%20prediction%20scaffold9_size8259397.466.mRNA1 |
| gene_0653 | EVM%20prediction%20scaffold9_size8259397.472.mRNA1 |
| gene_0660 | EVM%20prediction%20scaffold9_size8259397.62.mRNA1 |
| gene_0661 | EVM%20prediction%20scaffold9_size8259397.674.mRNA1 |
| gene_0662 | EVM%20prediction%20scaffold9_size8259397.717.mRNA1 |

**Supplementary Table S5. Nuclear loci used for phylogenomic analyses.** Mapping of the 356 nuclear loci retained for concatenated, multispecies-coalescent and phylogenetic-network analyses to their corresponding Passiflora organensis reference annotation identifiers. Analytical locus identifiers (gene_XXXX) were assigned sequentially during construction of the ≥6-of-8-taxa dataset. For genes with multiple annotated transcripts, the longest coding-sequence isoform was retained.

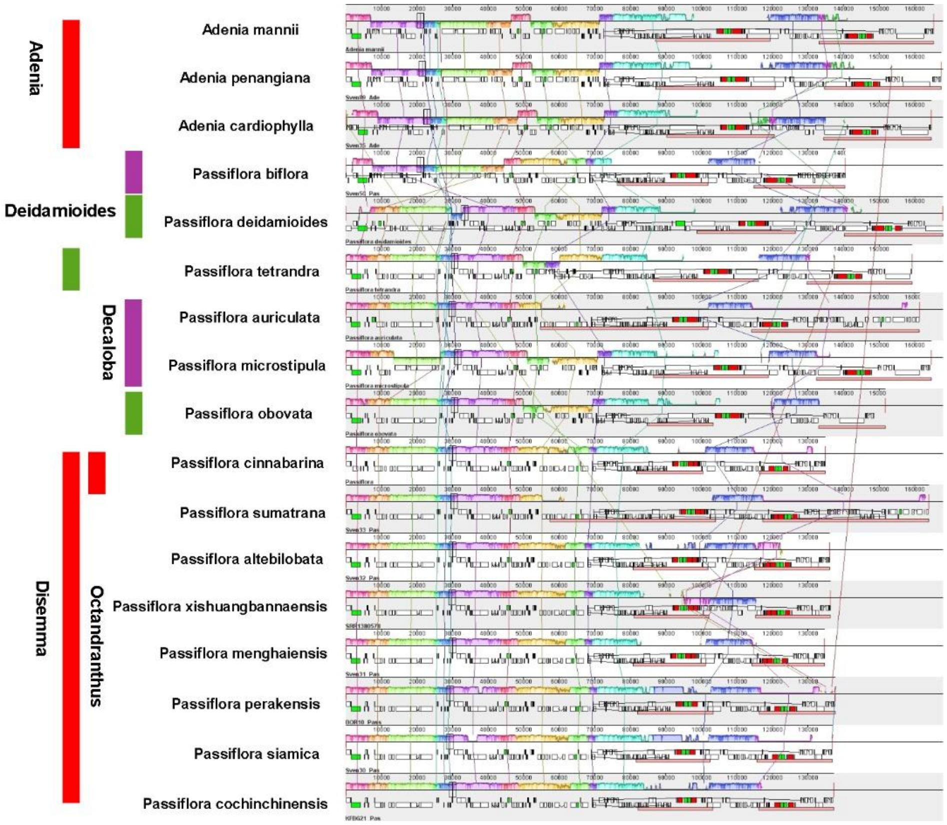


**Supplementary figure S6 ProgressiveMauve Comparative plastome structure across *Adenia* and *Passiflora*.** Linearized plastome maps showing gene order and structural variation among three *Adenia* species and representatives of *Passiflora* supersect. *Disemma*. Plastomes were standardized to a common orientation to facilitate comparison.

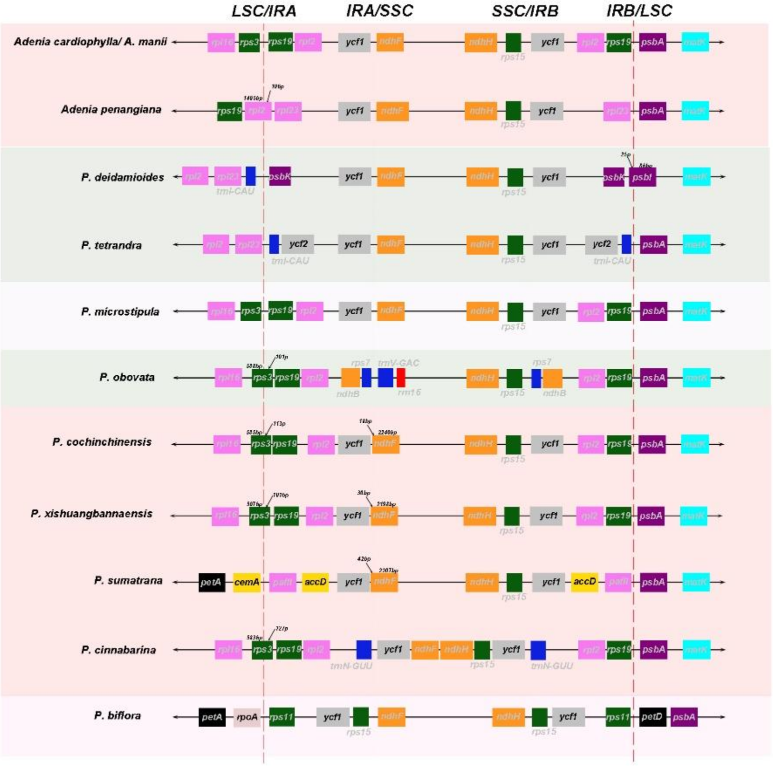


**Supplementary Figure S7. Comparison of inverted-repeat (IR) boundaries among selected *Adenia* and *Passiflora* plastomes.** Positions of genes adjacent to or spanning the four junctions between the large single-copy (LSC), small single-copy (SSC), and inverted-repeat regions (IRA and IRB) are shown for each taxon. Dashed vertical lines indicate the approximate positions of the LSC/IRA and IRB/LSC boundaries. Differences in gene position across the junctions illustrate lineage-specific expansions and contractions of the inverted repeats.

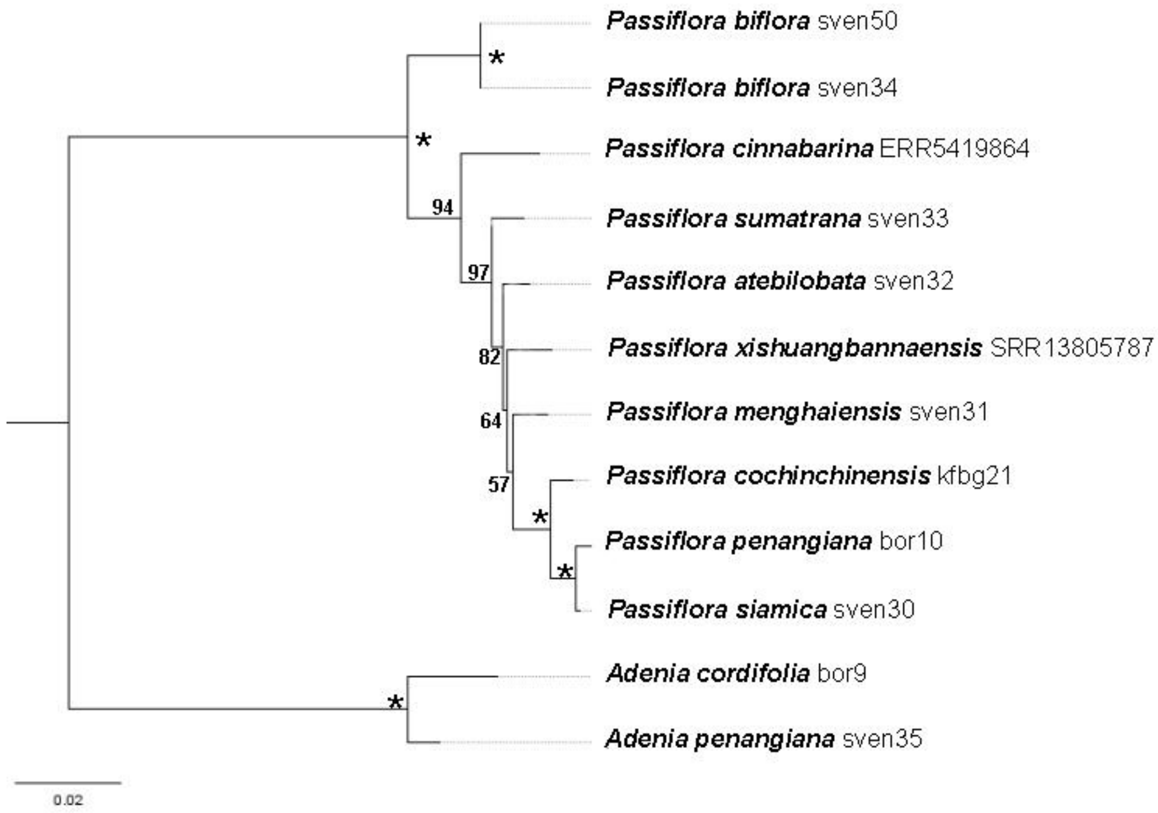


**Supplementary figure S8. Nuclear ribosomal cistron phylogeny of Asian *Passiflora* and *Adenia***. Maximum-likelihood phylogeny inferred from the nuclear ribosomal cistron sequences of the sampled taxa. Numbers at nodes indicate bootstrap support values; asterisks indicate nodes with maximal (100%) bootstrap support. Branch lengths are proportional to the number of nucleotide substitutions per site, as indicated by the scale bar. Two *Adenia* accessions were included as the outgroup.

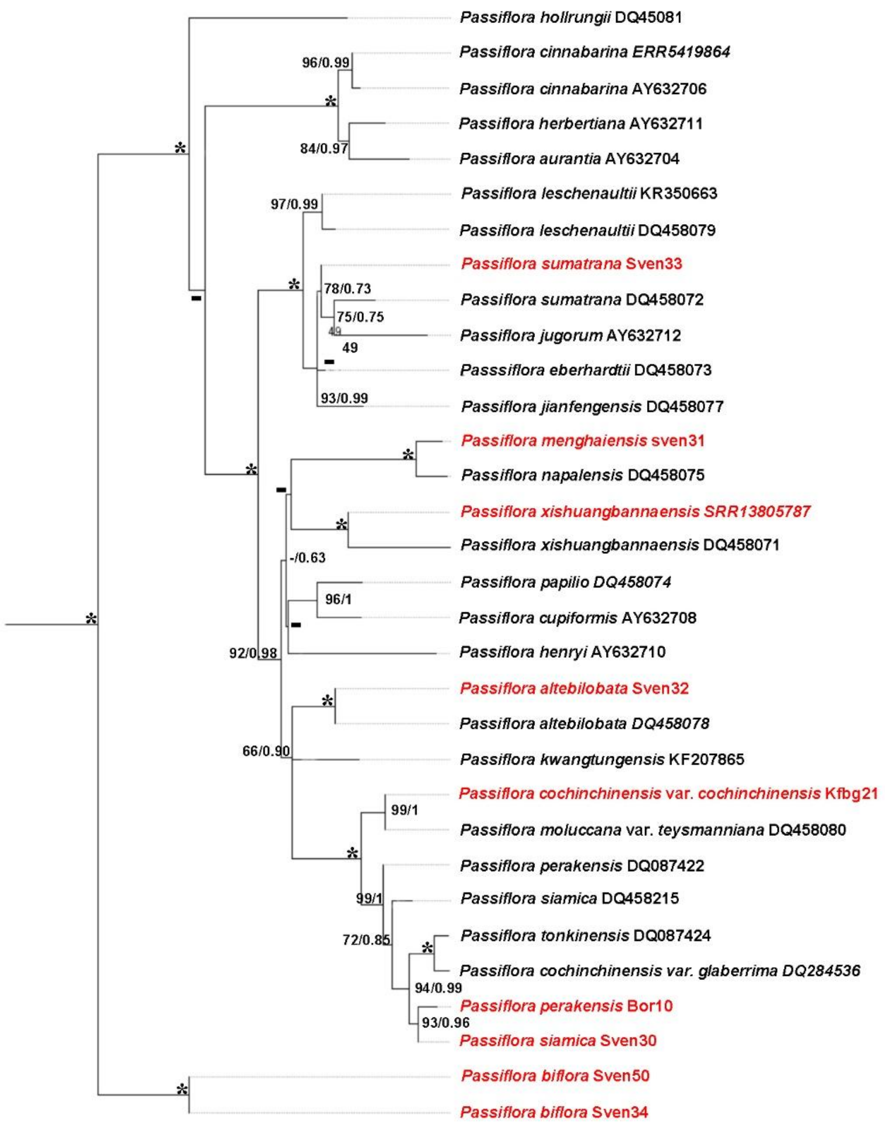


**Supplementary figure S9. ITS phylogeny of Asian *Passiflora*.** Maximum-likelihood phylogeny inferred from the nuclear ribosomal ITS1–5.8S–ITS2 region using an expanded taxonomic sampling of Asian *Passiflora*. Newly generated sequences are highlighted in red and were analysed together with previously published GenBank accessions. Node labels indicate maximum-likelihood bootstrap support/Bayesian posterior probability; * indicate maximally supported nodes, - indicate no supported node. The expanded ITS sampling was used to assess the placement of the newly sampled taxa relative to previously described Asian *Passiflora* species.

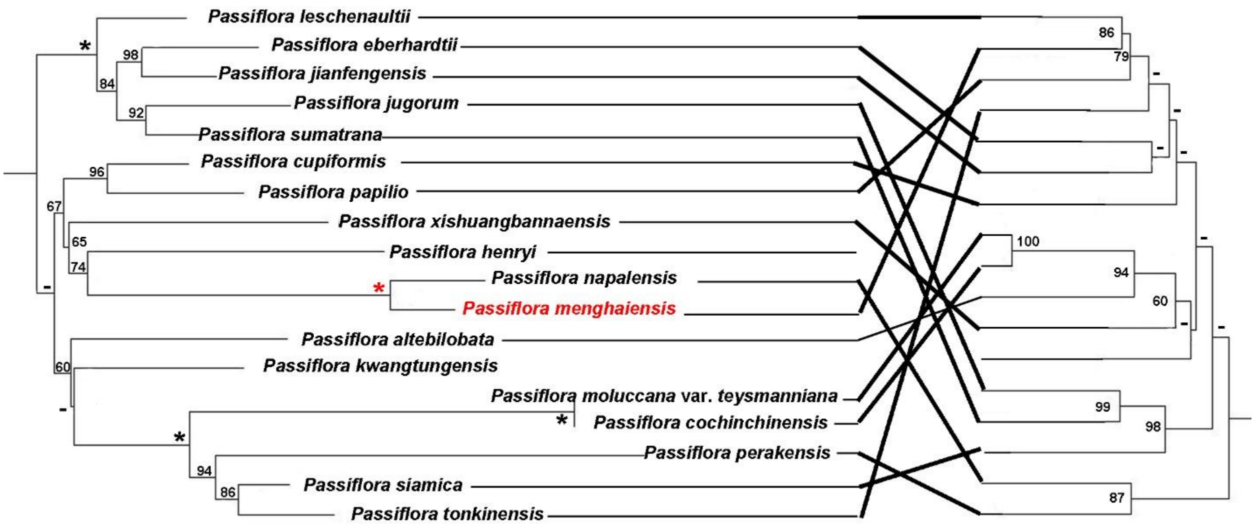


**Supplementary figure S10. Comparison of phylogenetic relationships inferred from nuclear and plastid markers within Asian Passiflora supersection *Disemma***. Tanglegram comparing phylogenetic relationships inferred from the nuclear 18S–ITS1–5.8S–ITS2–26S ribosomal cistron, low-copy nuclear cytGS and ncpGS (GS2) genes, and plastid trnL–trnF and ndhF regions. Phylogenetic trees were reconstructed using maximum likelihood in IQ-TREE, and numbers adjacent to nodes indicate bootstrap support values; asterisks indicate nodes with maximal bootstrap support. Lines connect identical taxa across trees and highlight topological congruence and discordance among nuclear and plastid markers. Taxon sampling incorporates sequences available from NCBI GenBank supplemented with sequences generated in the present study.

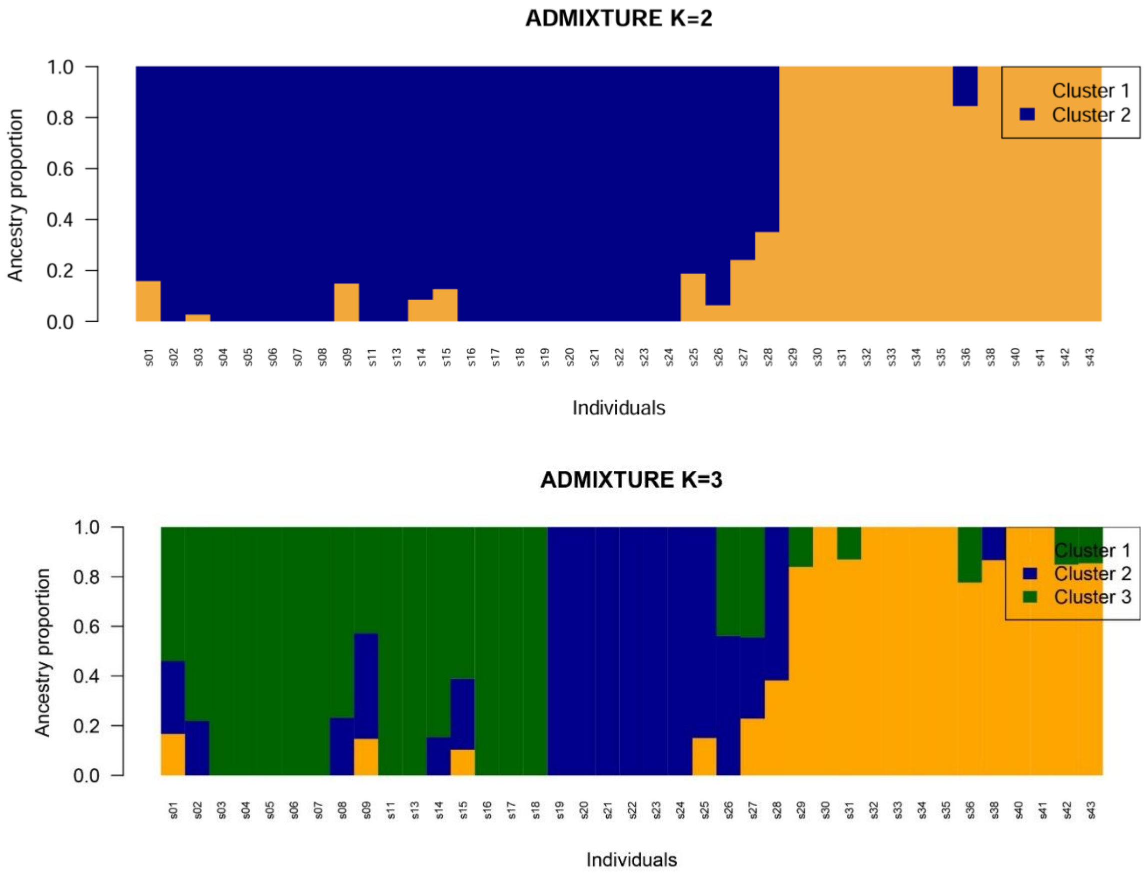


**Supplementary Figure S11. Population structure of *Passiflora xishuangbannaensis* inferred using ADMIXTURE.** Individual ancestry proportions are shown for K = 2 (upper panel) and K = 3 (lower panel) based on the LD-pruned nuclear SNP dataset. Each vertical bar represents one individual, and colours indicate the estimated proportion of ancestry assigned to each genetic cluster. At K = 2, individuals separate predominantly into two major genetic groups, with individual 28 showing intermediate ancestry. At K = 3, additional substructure is apparent within the first group, while the distinction between the two principal population groups is retained.

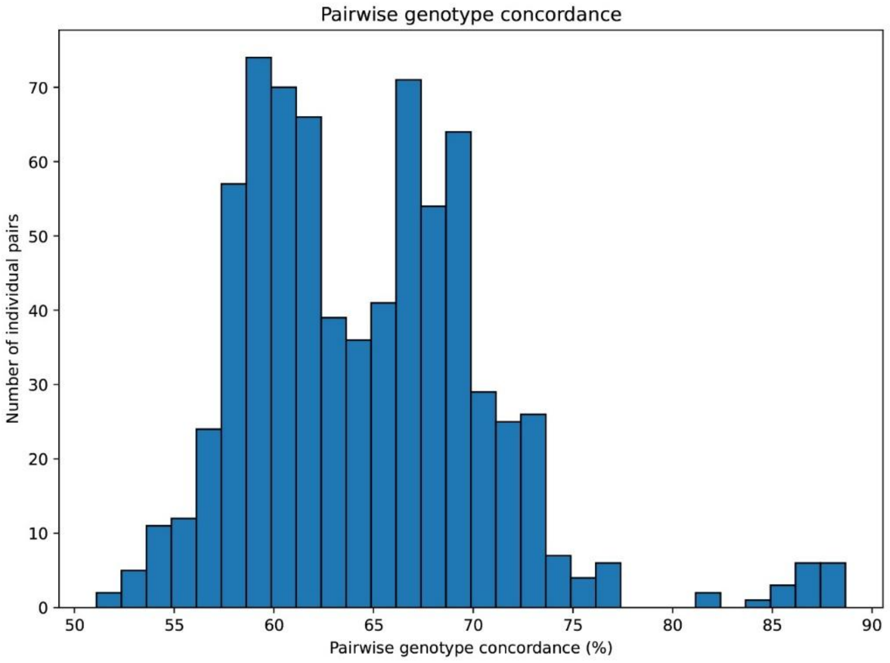


**Supplementary Table S12. Pairwise genotype concordance (%) among the 39 *P. xishuangbannaensis* individuals**. Concordance was calculated independently for each pair using only SNP loci for which genotype calls were available in both individuals; loci missing in either member of a pair were excluded from that comparison.

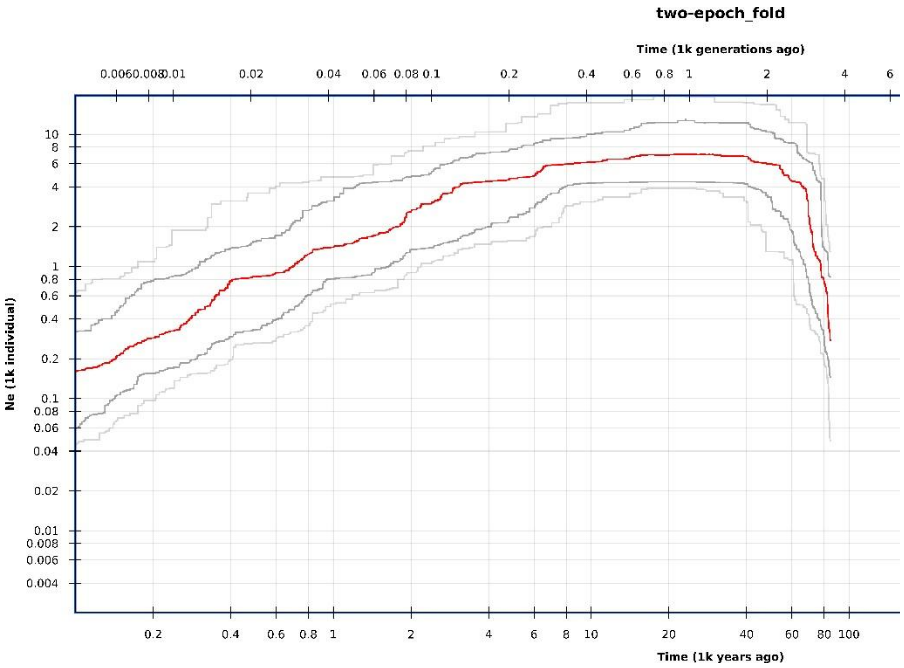

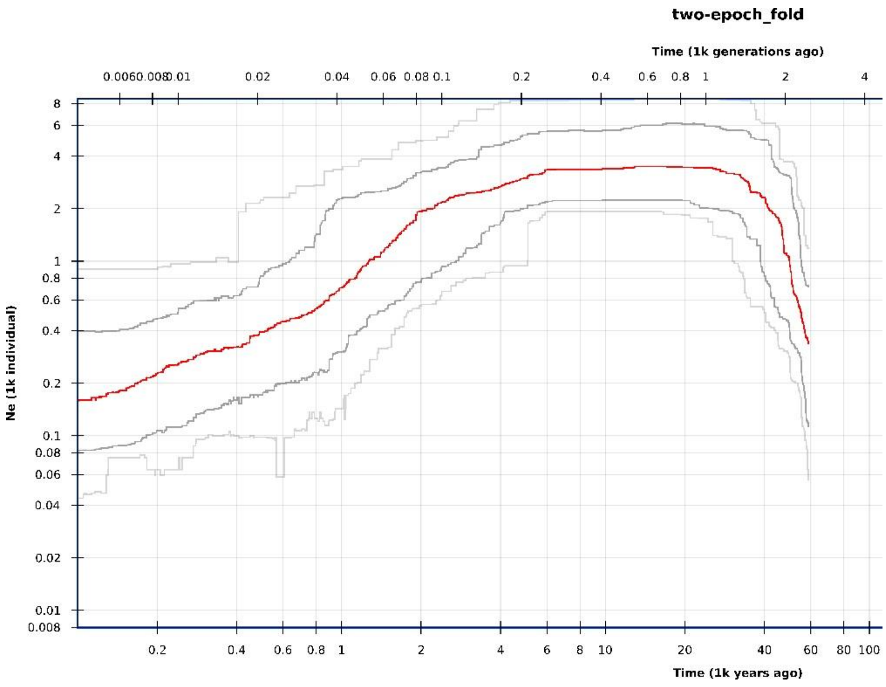


**Supplementary Figure S13. Historical demographic trajectories of the two principal populations of *Passiflora xishuangbannaensis* inferred using Stairway Plot 2.** Changes in effective population size (Ne) through time are shown separately for Pop1 (upper panel) and Pop2 (lower panel) based on the folded site-frequency spectrum. Red lines represent the median estimated Ne trajectory, while black and grey lines delimit the associated confidence intervals. Time is shown in thousands of years before present on the lower x-axis and thousands of generations on the upper x-axis. Demographic scaling assumed a generation time of 24 years.

### Supplementary Methods S14 - Exploratory screening of candidate self-incompatibility regions

To determine whether population differentiation in *Passiflora xishuangbannaensis* extended more broadly to genomic regions potentially associated with self-incompatibility (SI), an exploratory analysis was performed using candidate loci reported from the *P. organensis* genome annotation (Costa et al., 2021). These candidates were originally identified through similarity searches against Brassicaceae S-locus sequences and previously characterized P. edulis sequences, together with examination of protein-domain architectures characteristic of S-locus glycoproteins (SLG) and S-locus receptor kinases (SRK). Thus, SI candidates were not identified de novo from the *P. xishuangbannaensis* GBS data but were defined a priori from the *P. organensis* reference annotation. Following reconciliation of gene and scaffold identifiers with those used in the reference-based GBS dataset, 12 candidate annotation intervals were retained.

Because GBS provides sparse and discontinuous genomic coverage, SNPs were first intersected directly with the annotated candidate intervals and subsequently with windows extending ±10 kb and ±50 kb from each candidate gene. The extended windows were included to capture potentially linked variation surrounding SI candidates while accommodating the incomplete genomic representation inherent to GBS. SNPs associated with overlapping candidate intervals were subsequently identified by genomic position to avoid treating repeated SNP–annotation associations as independent variants.

Population differentiation at SI-associated variants was evaluated between Pop1 and Pop2 by matching candidate-region SNPs to locus-specific ($F_{st}$) estimates. The distribution of ($F_{st}$) among SI-associated variants was compared with the genome-wide distribution using a Wilcoxon test. Individual variants were additionally screened for elevated differentiation, with ($F_{st}$ >0.20) used to identify strongly differentiated loci and population-differentiation tests with (P<0.01) used to identify particularly strong candidates. Within-population variation was also examined from heterozygosity/homozygosity and inbreeding coefficients (F) calculated separately for Pop1 and Pop2, with SI-region values compared both between populations and against the corresponding genome-wide estimates.

Supplementary Results - Exploratory SI-region screening

Only three SNPs were recovered directly within the annotated SI candidate genes, illustrating the limited coverage of these regions by the GBS dataset. No SNPs were recovered in the ±10-kb dataset, whereas extension to ±50 kb produced 98 SNP-annotation associations. After accounting for variants associated with more than one overlapping candidate interval, these corresponded to 65 unique SNP positions. Of these SI-associated variants, 29 SNPs could be matched to loci for which Pop1-Pop3 ($F_{st}$) estimates were available and were compared with a genome-wide background of 2119 SNPs.

Candidate SI regions showed particularly limited variation within populations. Two examined SI candidate regions contained markedly different numbers of variants, with approximately 3 SNPs in one region and 98 SNPs in the other before the final population-level filtering. After filtering for variants suitable for differentiation analyses, approximately 10 informative SNPs were retained for the principal $F_{st}$ comparison.

Within-population heterozygosity across the SI candidate regions was low. In contrast, allele frequencies at informative SI-associated SNPs differed strongly between the two major populations, producing elevated between-population differentiation relative to the limited variation observed within populations.

These results indicate that the two genetically differentiated populations also carry differentiated allelic variation at loci potentially associated with self-incompatibility. The combination of low within-population SI diversity and high between-population differentiation is particularly important for conservation because genetically isolated populations may retain only subsets of the species-wide diversity at reproductive compatibility loci.

The SI results should nevertheless be interpreted as evidence of differentiation in candidate incompatibility-associated genes, rather than direct demonstration of functional incompatibility between individuals. Functional crosses or direct characterization of the incompatibility system would be required to establish whether the observed genomic differentiation translates into reduced reproductive compatibility between or within populations.
